# Platform-specific count-matrix preprocessing workflows for spatial transcriptomics data analysis

**DOI:** 10.64898/2026.09.24.753981

**Authors:** Yixuan Du, Sijie Li, Xinwang Yang, Han Shu, Zhikang Wang, Daoliang Zhang, Qi Zou, Zhiyuan Yuan

## Abstract

Preprocessing of spatial transcriptomics (ST) count matrices is critical for removing technical variation while preserving biological signals, but optimal strategies remain unclear across diverse platforms. We systematically benchmarked 371 preprocessing workflows across 45 data spanning 11 mainstream ST platforms, evaluating their performance using specifically designed complementary metrics. To further assess preprocessing factors upstream of count-matrix generation, we additionally evaluated image preprocessing, including cell segmentation and post-segmentation transcript processing, across representative imaging-based ST platforms. We found that no universal count-matrix preprocessing workflow performed optimally across all platforms, while data from the same platforms exhibited striking consistency in optimal count-matrix preprocessing workflows. We identified the key data structure characteristics and count-matrix preprocessing steps that critically influence preprocessing performance. We developed platform-specific and data structure-guided recommendations to assist users in selecting the optimal count-matrix preprocessing workflow, validated across a broader range of downstream tasks. We demonstrated that our data-driven recommendations uncover subtle biologically meaningful spatial patterns in new data that were obscured by suboptimal count-matrix preprocessing. Our findings suggest best practices for ST data count-matrix preprocessing.

## Introduction

The rapid diversification of ST platforms has created a complex landscape of data types with fundamentally different characteristics, ranging from sequencing-based platforms (sST) to imaging-based platforms (iST). These platform differences manifest as distinct data structures varying in gene panel size, transcripts per cell, spatial resolution, and data scale (Fig. 1a). However, ST count matrices are inherently challenging to analyze, as they are characterized by substantial technical noise, heteroscedasticity, and sparsity that can confound downstream analyses if not properly addressed. Therefore, count-matrix preprocessing serves as a critical bridge between ST measurements and biological interpretation, yet optimal count-matrix preprocessing workflows remain poorly defined across diverse ST platforms.

**Fig. 1.**
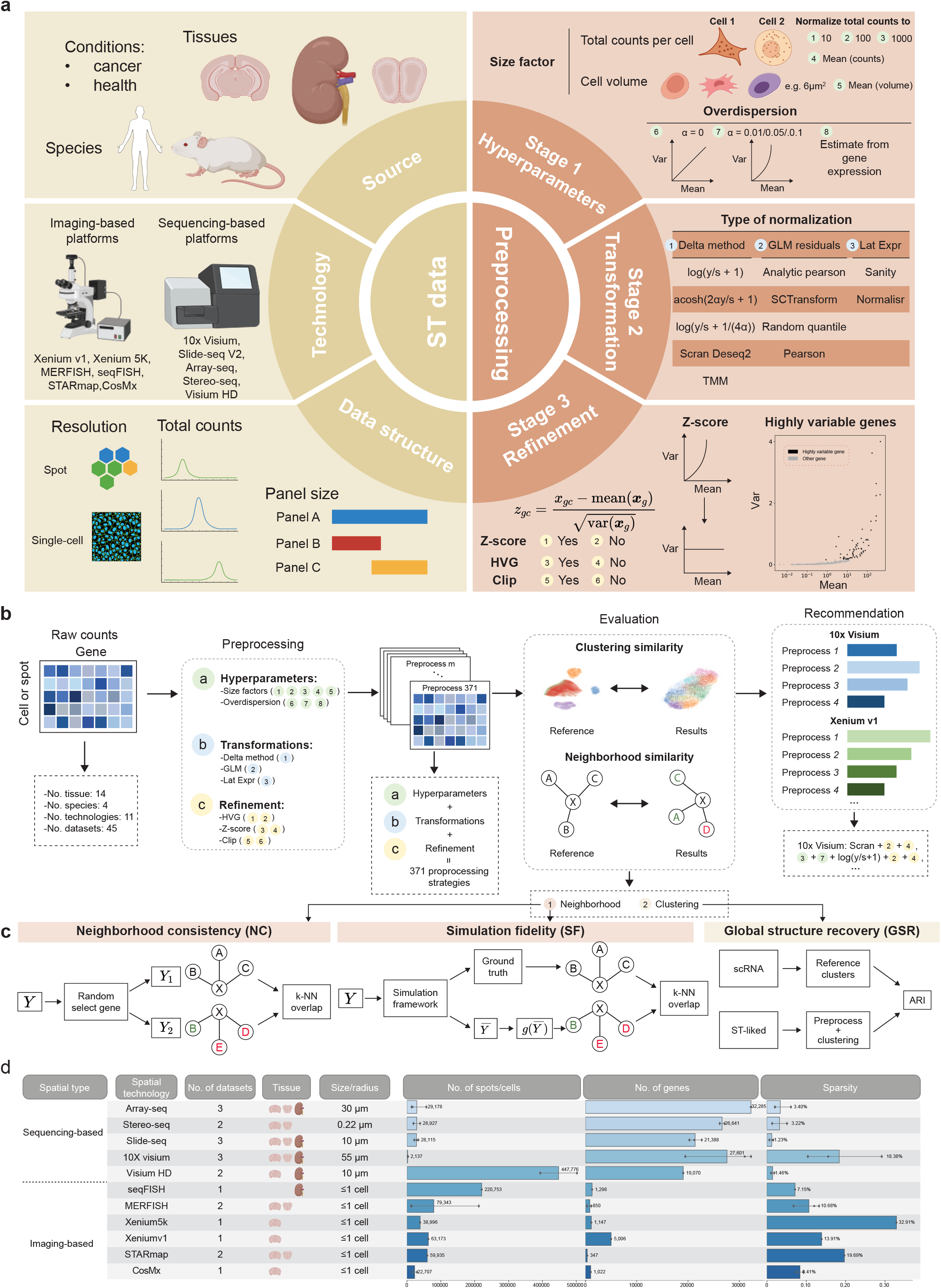
Overview of preprocessing workflow and study design. **a**, The motivation of our study. The left part illustrates diversity in ST data, including: data source, spatial resolution, and platforms. The right part illustrates preprocessing workflows. Our preprocessing workflow included three stages. Stage 1 involves setting the hyperparameters for the subsequent transformations, specifically the size factor and overdispersion. Stage 2 is transformation. We considered 12 distinct transformations, categorized into three types, or alternatively, no transformation was applied. Stage 3 consists of refinement operations, including HVG selection, Z-score normalization and clipping. Some elements in this figure are sourced from BioRender. **b**, The benchmarking framework for preprocessing, including Input data, Preprocessing, Evaluation, and Recommendation. **c**, Overview of the evaluation metrics, including NC, SF and GSR (See Methods for details). **d**, Overview of data from same tissue type (MCH, MOB, and MKD) in this study, including platform, number of data, tissue type, spatial resolution, number of spots/cells, number of genes, and data sparsity. Bar lengths represent mean values for all slices, with exact numbers labeled. The number of scatter points corresponds to the number of slices.

Current ST count-matrix preprocessing workflows largely adopt methods developed for single-cell RNA sequencing (scRNA-seq). While extensive work has optimized scRNA-seq preprocessing^7^ (e.g., studies on transformation methods^2^, overdispersion modeling^8,9^, and clustering performance^10^), ST data possess unique characteristics that may violate assumptions underlying these approaches^3,4,59^. For example, sST data often suffer from cellular heterogeneity within spots (multiplexing of cells), spatial autocorrelation introduces dependencies absent in traditional scRNA-seq data, and the vastly different transcript capture efficiencies across ST platforms suggest that scRNA-seq best practices cannot be directly transposed to ST data without rigorous validation. For iST platforms, the count matrix is additionally influenced by upstream image preprocessing, particularly cell segmentation and post-segmentation transcript assignment or purification, which directly determine how detected transcripts are assigned to individual cells. Thus, ST preprocessing involves multiple interconnected steps, highlighting the need for a systematic evaluation framework to comprehensively assess these procedures and establish robust preprocessing workflows for ST data.

Here, we address this gap through comprehensive benchmarking of 371 count-matrix preprocessing workflows across 45 data spanning 11 ST platforms. Our framework systematically evaluates combinations of hyperparameter settings (size factors and overdispersion), transformation methods (12 approaches spanning delta method^11^, Generalized Linear Model (GLM) residuals^12^, and latent expression models^13,14^), and refinement operations (highly variable gene selection, Z-score normalization and clipping, see Methods “Preprocessing” for more details, Supplementary Fig. 2). In addition, for representative imaging-based ST platforms, we extended the framework upstream to evaluate image preprocessing procedures, including cell segmentation and post-segmentation transcript processing, that directly determine count-matrix construction. We assess count-matrix preprocessing quality through metrics capturing both local graph structure and global organization (Fig. 1c), while controlling for biological variables by using matched tissue types across platforms. Finally, to ensure the robustness of our conclusions, we validated our findings on 24 additional data and assessed the impact of preprocessing on downstream analysis performance.

Our analysis reveals that while no universal count-matrix preprocessing workflow performs optimally across all scenarios, distinct patterns emerge. Specifically, delta method-based transformations provided a consistently strong foundation across data. However, the optimal hyperparameters and refinement operations were highly platform-dependent. Importantly, the relative performance of count-matrix preprocessing workflows remained highly consistent across data generated using different upstream image preprocessing procedures, including different cell-segmentation and post-segmentation transcript-processing strategies. Formal variance-partitioning analyses further showed that platform remained the strongest measured factor associated with variation in preprocessing preferences, exceeding the contributions of the measured biological factors, image preprocessing workflows. At the same time, we identified gene panel size and gene expression level per unit (GELPU) as important intrinsic data-structure characteristics associated with preprocessing performance. Consequently, we developed platform-specific recommendations for technologies represented in our benchmark, complemented by data structure-guided strategies based on panel size and GELPU. This approach ensures our guidelines are not only effective for current technologies but remain extensible to novel ST platforms developed in the future.

## Results

### Comprehensive benchmarking framework for ST count-matrix preprocessing workflow

We designed a three-stage count-matrix preprocessing framework encompassing all major strategic decisions in current ST count-matrix preprocessing workflows (Fig. 1a). The first stage involves setting hyperparameters of transformation, including size factors (counts-based^15^ and spatially-aware cell volume-based^3^) for normalizing total counts and overdispersion (*α*) parameters for Negative Binomial (NB) model-based transformations^16-18^, with specific values of *α* (0, 0.01, 0.05, 0.1 or gene-specific estimation) tested. The second stage applies transformations to normalize counts and stabilize variance^2^. We considered 12 distinct transformations (Methods; Fig. 1a), spanning three conceptual categories: delta method, GLM residuals, and latent expression models (Methods). The third stage, refinement, involves three critical operations: Highly Variable Genes (HVG) selection (top 20%) for initial dimensionality reduction, Z-score normalization to standardize gene expression, ensuring all genes contribute equally to the definition of cell state, and residual clipping for GLM residual-based transformations to limit extreme residual values and reduce the influence of outliers. The systematic combination of these three stages resulted in the evaluation of 371 distinct preprocessing workflows (see Methods “Preprocessing” for more details).

To ensure our benchmark provides robust and generalizable insights, we curated a comprehensive collection of ST data. Our collection encompasses 11 platforms, including 10x Visium, Stereo-seq (8µm bin), Slide-seq V2, Array-seq, Visium HD, MERFISH, STARmap, Xenium v1, Xenium 5K, CosMx and seqFISH+ (Fig. 1d)^5,19-23,26,56-58^. To enable a systematic comparison across platforms while minimizing biological confounders, we focused on three tissue types: mouse brain sections encompassing cortex and hippocampus (MCH), mouse olfactory bulb (MOB), and kidney (MKD), each of which has been profiled by multiple ST platforms in our panel. This study design allows us to hold the tissue type constant and primarily attribute observed differences to the underlying platforms. The collection encompasses both major categories of spatial platforms: iST and sST platforms. They vary significantly across key dimensions such as the gene panel size, data scale (number of spots/cells), and spatial resolution (from low-resolution spots to subcellular resolution), as well as underlying tissue architecture. Throughout this manuscript, “cells” refers to the analytical units of iST platforms, whereas “spots” refers to the spots or bins used as analytical units in sST platforms; when both platform types are discussed together, we use “spots/cells”. Furthermore, to provide even more reliable and generalizable results, we conducted extensive validation on 24 additional data (Supplementary Fig. 1). This comprehensive validation set ensures the robustness of our conclusions across a wider range of biological contexts and experimental variations. For clarity, throughout this study, “Data” denotes one independently analyzed cell/spot-by-gene count matrix used as a basic unit in the benchmark, with each data entry assigned a unique identifier (e.g., Data 1, Data 2, …). A “tissue section” refers specifically to an individual physical section subjected to ST profiling, whereas a “biological sample” refers to the underlying biological specimen or source from which one or more tissue sections or measurements were obtained. Multiple Data entries generated using the same ST technology are collectively referred to as data from the same platform.

Following the extensive collection of diverse ST data, we designed an evaluation framework with three complementary metrics: neighborhood consistency (NC), simulation fidelity (SF), and global structure recovery (GSR). NC metric quantifies local graph quality using *k*-nearest neighbor (*k*-NN) graph consistency, adapted from the *k*-NN graph-based benchmarking framework introduced for single-cell RNA-seq data^2^, by measuring the agreement of neighborhood structure across random gene subsets. SF metric assesses how well expected relationships are preserved in controlled simulations, following the *k*-NN graph-based principles as in Ref^2^. GSR evaluates global structure quality using a recovery strategy conceptually related to a recent Xenium benchmarking study^6^: we collected matched scRNA-seq data with ground-truth cell type labels, transformed them to mimic ST characteristics through downsampling and gene filtering, applied each preprocessing strategy, performed clustering, and computed the Adjusted Rand Index (ARI) between inferred clusters and true cell types (more details see Methods “Benchmark Metrics”). For each platform, we calculated overall ranking scores by averaging rank-normalized values across the three metrics, and validation on the above mentioned 24 additional data confirmed the generalizability of these rankings (see Methods “Benchmark Metrics” for more details).

### Optimal count-matrix preprocessing workflows Across iST Platforms

We first evaluated the performance of count-matrix preprocessing workflows on iST platforms. Our benchmark included representative iST technologies MERFISH, STARmap, Xenium v1, Xenium 5K, CosMx and seqFISH+. For each iST platform, count-matrix preprocessing workflows were compared within the same tissue type, thereby holding tissue type constant so that observed differences can be primarily attributed to preprocessing rather than biological variation. We evaluated each count-matrix preprocessing workflow using the three metrics detailed previously: NC, SF, GSR. The overall ranking scores were calculated by averaging rank-normalized values across the three metrics (Methods). According to the overall ranking score, the top-performing and bottom-performing count-matrix preprocessing workflows are visualized for comparison. Figure 2a and Supplementary Figure 3a present the results for iST platforms on MCH tissue. Correspondingly, Supplementary Figure 4a and Supplementary Figure 5a present the results derived from the MOB and MKD tissues. We first analyzed the benchmark results derived from the MCH tissue for demonstration (Fig. 2a-c, Fig. 2e, Fig. 2f). Unless otherwise specified, the following analyses focus on the iST Data entries derived from MCH tissue.

**Fig. 2.**
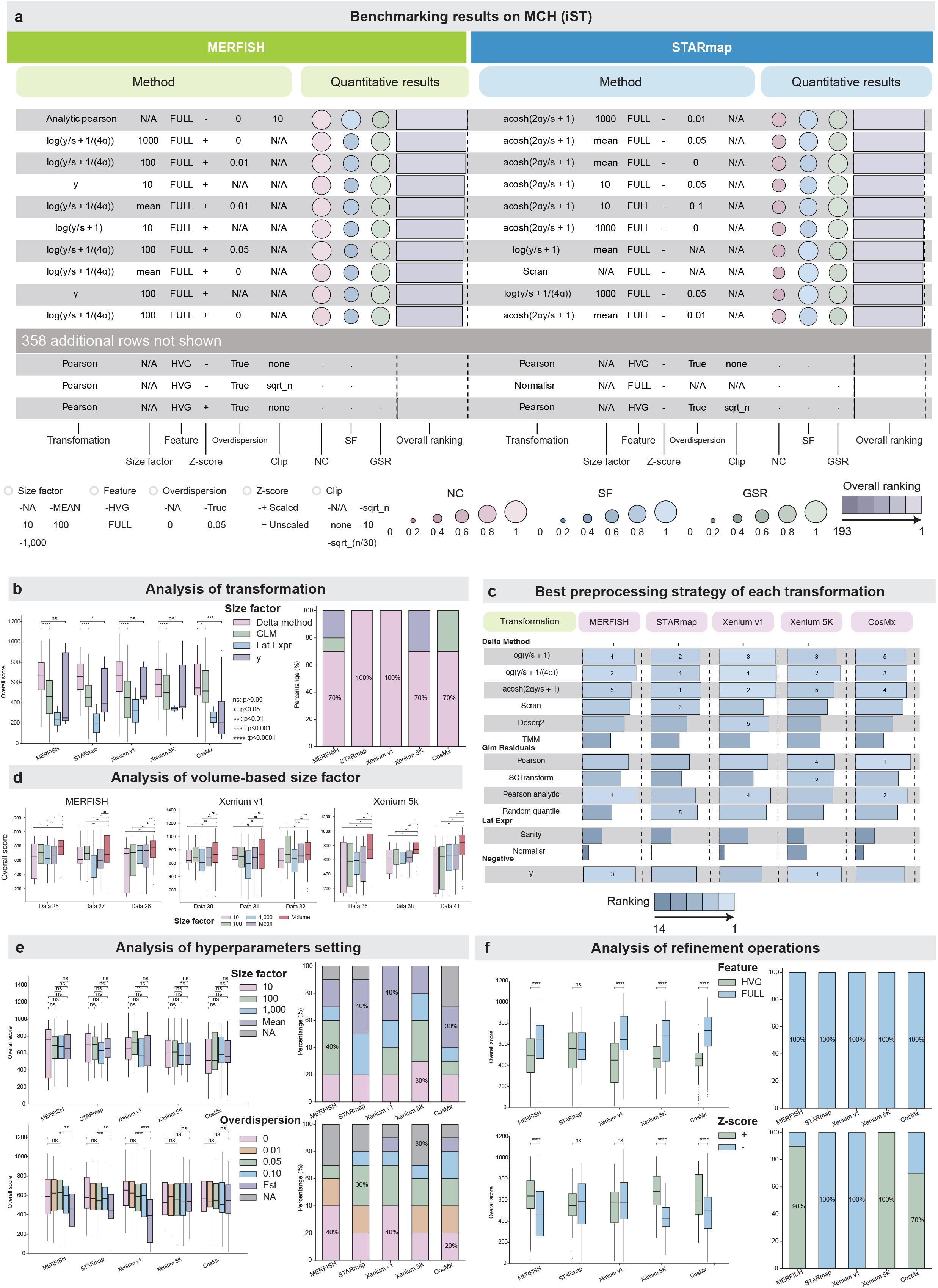
Benchmarking results for iST platforms on MCH tissue. **a**, Overview of top 10 and bottom 3 ranked preprocessings by overall ranking score. Metrics included NC (pink), SF (blue) and GSR (green). Overall ranking scores are computed as the average of the three rank-normalized metrics. Figure 2a presents the results for MERFISH and STARmap platforms on MCH tissue. Supplementary Figure 3a presents the results for Xenium v1 and Xenium 5K platforms on MCH tissue. This figure was generated using the R package funkyheatmap^48^. **b**, Analysis of transformation across different iST MCH data. The grouped boxplot in the left part shows the distribution of the overall ranking score across 4 transformation types, with ‘y’ representing the no-transformation. Each group in the plot represents the individual iST platform MCH data, and the different columns within each group correspond to the distinct transformation types. For all boxplots in this manuscript, center line, median; box limits, upper and lower quartiles; whiskers, 1.5x interquartile range; points, outliers. *P*-values were calculated using a two-sided Mann-Whitney U test. **\*p** < 0.05; **\*\*p** < 0.01; **\*\*\*p** < 0.001; **\*\*\*\*p** < 0.0001; ns, not significant. The stacked bar plot in the right part shows the composition of the top 10 performing preprocessing workflows. Each bar represents an individual iST platform MCH data, with the stacked segments within the bar showing the frequency of each transformation type present in that data’s top-performing subset. **c**, Overview of the best-performing preprocessing strategies for 13 transformations across different iST MCH data. The lengths of the bars represent the magnitude of the rank-based overall scores. **d**, Analysis of volume-based size factor. The three grouped boxplots, presented from left to right, display the results for the MERFISH, Xenium v1, and Xenium 5K platforms. Within each plot, each group represents an individual data corresponding to that specific platform (the data IDs for data are detailed in Supplementary Fig. 1). The columns within each group correspond to preprocessings that employed different size factors. **e, f**, Analysis of hyperparameters settings and refinement operations. The grouped boxplot in the left part of each figure shows the distribution of the overall ranking score across different hyperparameters settings or refinement operations (**e** for hyperparameters settings and **f** for refinement operations). Each group in the plot represents the individual iST platform MCH data, and the different columns within each group correspond to preprocessings that employ distinct hyperparameters or refinement operations. *P*-values were calculated using a two-sided Mann-Whitney U test. *p < 0.05; **p < 0.01; ***p < 0.001; ****p < 0.0001; ns, not significant. The stacked bar plot in the right part of each figure shows the composition of the top 10 performing preprocessing workflows. Each bar represents an individual iST platform MCH data, with the stacked segments within the bar showing the frequency of each hyperparameter setting or refinement operation in that data’s top 10 performing subset.

Regarding the transformation step, delta method-based transformations (a transformation category, Supplementary Fig. 2) excel across all iST platforms. For the top-performing count-matrix preprocessing workflows, we observed that the delta method accounted for over 70% in the top 10 count-matrix preprocessing workflows across all the iST platforms (Fig. 2b, right). Notably, GLM-based transformations (a transformation category; Supplementary Fig. 2) were represented among the top 10 count-matrix preprocessing workflows only for MERFISH and CosMx, accounting for 10% and 30% of the top-performing workflows, respectively (Fig. 2b, right). Additionally, simply applying Z-score normalization to raw counts also yielded competitive results on the MERFISH and Xenium 5K data, where they accounted for 20% and 30% among the top 10 count-matrix preprocessing workflows (Fig. 2b, right). This finding suggests that the raw counts in some iST data inherently preserve highly meaningful biological information, which may be attributable to the high specificity and detection efficiency of these iST platforms, as suggested by Wang et al^1^. Conversely, the latent expression methods (a transformation category, Supplementary Fig. 2) exhibited suboptimal performance across all iST platforms, as none of the top-performing count-matrix preprocessing workflows employed them (Fig. 2b, right). When considering the performance of all count-matrix preprocessing workflows, the delta method-based transformations achieved a higher performance than other transformations across all the iST data (Fig. 2b, left). To further investigate the differences between the various transformation methods, we focused on the best-performing count-matrix preprocessing workflows corresponding to each transformation method (Fig. 2c). We found that 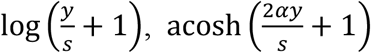 and 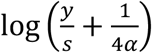 exhibited consistently superior performance on all the iST data (Fig. 2c). In contrast, on the MERFISH and CosMx data, the GLM-based transformations (Pearson^12^ and Analytic pearson^9^) also exhibited strong performance (Fig. 2c). We performed the same analysis on the other two tissues, and the conclusions are consistent (Supplementary Fig. 4, Supplementary Fig. 5).

Hyperparameter analysis revealed distinct patterns. For size factors, focusing solely on the top-performing workflows suggested that the choice of size factors did not yield significant differences in benchmark results, as almost all tested values were well-represented among the top 10 workflows (Fig. 2e, upper right). However, the analysis across all tested strategies demonstrated that setting the size factors to 10 or 100 exhibited superior performance for the MERFISH and STARmap and Xenium 5K data, while mean or 1,000 exhibited superior performance for the CosMx data, mean or 100 exhibited superior performance for the Xenium v1 data. This difference may be attributable to the varying capture efficiencies of the different iST platforms (Fig. 2d, upper left). Beyond simple count-based size factor normalization, we also tested the performance of mean cell volume as an alternative size factor^3^. Due to the absence of cell volume information in the STARmap and seqFISH+ platforms, our analysis focused on the MERFISH, Xenium v1, and Xenium 5K platforms, utilizing three data from each platform (Fig. 2d; data IDs are listed in Supplementary Fig. 1). Since the GSR metric requires ground truth labels derived from paired scRNA-seq data, which lack cell volume information, we were restricted to evaluating performance using only the *k*-NN Graph-based metrics (NC and SF) for these data. We found that using cell volume achieved marginally better results on all the data, while the difference in performance compared to the counts-based size factor was not significant in some data (Fig. 2d). Overall, these results are broadly consistent with the findings of Jean Fan et al., who suggested that cell volume-based normalization can reduce error rates in gene differential expression analysis, while further indicating that the magnitude of this improvement may vary across iST data^3^.

In contrast to the size factor, the optimal setting for overdispersion (*α*) was consistent across most iST platforms. On MERFISH, STARmap and Xenium 5K data, we observed that among the top-performing workflows that utilized overdispersion, none employed the strategy of estimating overdispersion individually for each gene (Fig. 2e, lower right). Instead, they all used fixed values for the overdispersion setting (Fig. 2e, lower right). A notable exception was the Xenium v1 and CosMx data, where 10% of the top workflows selected the estimation from data approach (Fig. 2e, lower right). But when focusing on the complete set of all workflows, fixed values setting of overdispersion have better performance than estimation from the data across all the iST platforms (Fig. 2e, lower left). This result notably contrasts with findings reported in the scRNA-seq field^2^, where estimating overdispersion from data or using *α* = 0.05 is typically preferred over *α* = 0. This divergence suggests that the phenomenon of overdispersion might be less pronounced or statistically less reliable to estimate in iST data compared to scRNA-seq data. The conclusions remained consistent across the other two tissues (Supplementary Fig. 4e, Supplementary Fig. 5e).

Analysis of refinement operations revealed divergent optimal settings across iST data. Regarding HVG selection, we observed that all the top-performing preprocessings utilized all features without performing HVG selection across MERFISH, STARmap, Xenium v1, Xenium 5K and CosMx data (Fig. 2f, upper right). This pattern was maintained when focusing on all workflows, we found that workflows utilized the full feature consistently and significantly outperformed those that performed HVG selection. (Fig. 2f, upper left). This result is likely attributable to the fact that the panel sizes of iST platforms are generally much smaller than those of scRNA-seq, and HVG selection on iST data tends to cause an undesirable loss of critical biological information necessary for defining cellular states. Conversely, Z-score normalization revealed different preferences. On MERFISH, Xenium5K, and CosMx data, the majority (> 70%) of top-performing workflows performed the Z-score normalization (Fig. 2f, lower right). However, the opposite trend was observed on STARmap and Xenium v1 data, where all the top-performing pipelines omitted this standardization step (Fig. 2f, lower right). When considering all tested workflows, count-matrix preprocessing that employed Z-score normalization generally outperformed those that did not on most iST data (Fig. 2f, lower left). The conclusions remained mostly consistent across the other two tissues (Supplementary Fig. 4f, Supplementary Fig. 5f).

### Optimal count-matrix preprocessing workflows Across sST Platforms

Following our analysis of iST platforms, this module shifts the focus to the benchmarking on sST platforms. Our benchmark included representative sST technologies 10x Visium, Slide-seq V2, Stereo-seq, Visium HD and Array-seq, and the count-matrix preprocessing workflows performance was compared across the same tissue types, which is consistent with the iST benchmark. We evaluated count-matrix preprocessing workflows based on NC and SF metrics. GSR metric was not utilized for evaluation because some sST data exhibit very high total counts per spot, often exceeding those in scRNA-seq data, due to a spot potentially covering multiple cells. This data characteristic prevents effective downsampling to create sST-like data from corresponding scRNA-seq data. The overall ranking score was calculated by averaging rank-normalized values across the two metrics (Methods). The top-performing and bottom-performing count-matrix preprocessing workflows, determined by the overall ranking score, are visualized for comparison. Figure 3a and Supplementary Figure 3b present the results for sST platforms on MCH tissue. Correspondingly, Supplementary Figure 6a and Supplementary Figure 8a present the results for sST platforms derived from the MOB and MKD tissues. We first analyzed the benchmark results derived from the MCH tissue for demonstration (Fig. 3a-f). Unless otherwise specified, the following analyses focus on the sST Data entries derived from MCH tissue.

**Fig. 3.**
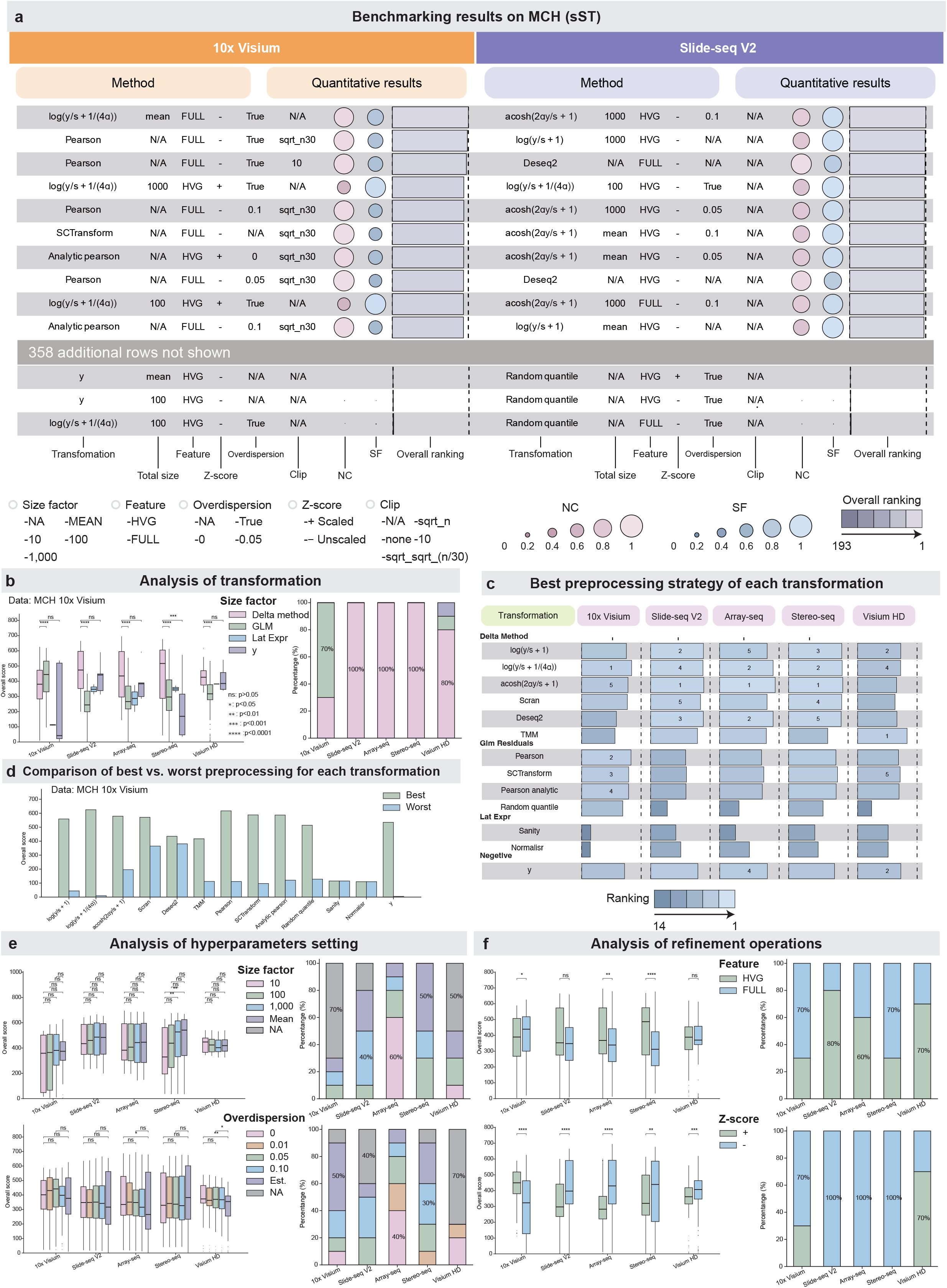
Benchmarking results for sST platforms on MCH tissue. **a**, Overview of top 10 and bottom 3 ranked preprocessings by overall ranking score. Metrics included NC (pink) and SF (blue). Overall ranking scores are computed as the average of the two rank-normalized metrics. Figure 3a presents the results for 10x Visium and Slide-seq V2 platforms on MCH tissue. Supplementary Figure 3b presents the results for Array-seq and Stereo-seq platforms on MCH tissue. **b**, Analysis of transformation across different sST MCH data. The grouped boxplot in the left part shows the distribution of the overall ranking score across 4 transformation types, with ‘y’ representing the no-transformation. Each group in the plot represents the individual sST platform MCH data, and the different columns within each group correspond to the distinct transformation types. *P*-values were calculated using a two-sided Mann-Whitney U test. **\*p** < 0.05; **\*\*p** < 0.01; **\*\*\*p** < 0.001; **\*\*\*\*p** < 0.0001; ns, not significant. The stacked bar plot in the right part shows the composition of the top 10 performing preprocessing workflows. Each bar represents an individual sST platform MCH data, with the stacked segments within the bar showing the frequency of each transformation type present in that data’s top-performing subset. **c**, Overview of the best-performing preprocessing strategies for 13 transformations across different sST MCH data. The lengths of the bars represent the magnitude of the rank-based overall scores. **d**, Comparison of the best- and worst-performing preprocessings for each transformation on 10x Visium MCH data. The grouped boxplot shows the overall ranking score of the best- and worst-performing preprocessings for each transformation. Each group in the plot represents a transformation. For each group, the first column shows the best preprocessing’s overall ranking score for this transformation, and the second column shows the worst preprocessing’s overall ranking score for this transformation. **e, f**, Analysis of hyperparameters settings and refinement operations. The grouped boxplot in the left part of each figure shows the distribution of the overall ranking score across different hyperparameters settings or refinement operations (**e** for hyperparameters settings and **f** for refinement operations). Each group in the plot represents the individual sST platform MCH data, and the different columns within each group correspond to preprocessings that employ distinct hyperparameters or refinement operations. *P*-values were calculated using a two-sided Mann-Whitney U test. *p < 0.05; **p < 0.01; ***p < 0.001; ****p < 0.0001; ns, not significant. The stacked bar plot in the right part of each figure shows the composition of the top 10 performing preprocessing workflows. Each bar represents an individual sST platform MCH data, with the stacked segments within the bar showing the frequency of each hyperparameter setting or refinement operation in that data’s top 10 performing subset.

Regarding the transformation step, delta method-based transformations excel across most sST platforms. For the top-performing workflows, we observed that the delta method accounted for over 80% in the top 10 workflows across almost all the sST platforms, with 10x Visium being the only exception, where it accounted for 30% (Fig. 3b, right). Additionally, applying HVG selection and Z-score normalization to raw counts also yielded competitive results on the Visium HD data, where they accounted for 10% among the top 10 workflows (Fig. 3b, right). The GLM-based transformations have good performance only on 10x Visium data, where it accounted for 30%. The latent expression methods (Supplementary Fig. 2) exhibited suboptimal performance across all sST data, as none of the top-performing preprocessing pipelines employed them (Fig. 3b, right). When considering the performance of all workflows, the delta method achieved a higher average performance than other transformations across the most sST data (Fig. 3b, left). Consistent with iST benchmark, we then focused on the best-performing workflows corresponding to each transformation method (Fig. 3c). We found that 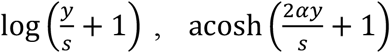, and 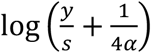 exhibited consistently superior performance across all the sST data (Fig. 3c). In contrast, on the 10x Visium and Visium HD data, the GLM-based transformations also exhibited strong performance (Fig. 3c). We performed the same analysis on the other two tissues, and the conclusions are consistent (Supplementary Fig. 7, Supplementary Fig. 8).

The optimal hyperparameter settings derived for sST platforms are consistent. First, we began our analysis of the size factors. Among the top-performing workflows, we found that size factors set at 10, 100, 1,000, or the mean resulted in similar performance, as all tested values were observed among the top 10 workflows, which is consistent with the iST conclusions (Fig. 3e, upper right). However, when considering all workflows, the optimal setting for sST data was consistently found to be size factors of 1,000 or mean (Fig. 3e, upper left). Second, regarding the overdispersion (*α*), the optimal setting was consistent with almost all the sST data, where a fixed value performed better (Fig. 3e, lower left), which is also consistent with the iST conclusions. The conclusions remained mostly consistent across the other two tissues (Supplementary Fig. 7, Supplementary Fig. 8).

In terms of refinement operations, the top-performing workflows showed divergent preferences across different sST data. For HVG selection, we observed that the majority (> 70%) of top-performing workflows utilized all features on 10x Visium and Stereo-seq data, while the majority (> 60%) of top-performing workflows performed HVG selection on Slide-seq V2, Array-seq and Visium HD data (Fig. 3f, upper right). When considering the performance of all workflows, we found that workflows performed HVG selection achieved superior average performance compared to those omitting HVG selection across most sST data (Fig. 3f, upper left). For Z-score normalization, we observed that the majority (> 70%) of top-performing workflows omitted Z-score normalization on most sST data (Fig. 3f, lower right). When considering the performance of all workflows, we found that workflows that omitted Z-score normalization achieved higher average performance than those that included it across most sST data (Fig. 3f, lower left). The conclusions remained mostly consistent across the other two tissues (Supplementary Fig. 7e, Supplementary Fig. 8f).

Interestingly, we found that the same transformations can yield significantly divergent performance based on their specific hyperparameter settings and refinement operations. For instance, in the Array-seq MOB data, the raw counts without transformation appeared in both the top-performing and bottom-performing groups due to variations in its accompanying settings (Supplementary Fig. 6a). By comparing the best and worst workflows for each transformation method across various data, we found that even the same transformation method combined with different combinations of hyperparameters and refinement operations can lead to substantially different results for both iST and sST platforms (Fig. 3d, Supplementary Fig. 4d, Supplementary Fig. 5d, Supplementary Fig. 7c, Supplementary Fig. 8d). This underscores the critical importance of hyperparameters and refinement operations in defining the efficacy of a preprocessing workflow.

### Z-score Normalization Exerts the Strongest Influence on Count-matrix Preprocessing Workflows

Given the consistently strong performance of the delta method-based transformations across both iST and sST platforms, and recognizing that count-matrix preprocessing performance is critically influenced by the choice of hyperparameter settings and refinement operations (as previously noted), this section specifically focuses on analyzing the influence of those hyperparameter settings and refinement operations. This analysis is important for identifying critical tuning parameters and establishing practical guidelines for robust ST data preprocessing. We investigated this influence through two aspects: the *k*-NN graph structure of the preprocessed data, and the overall ranking score derived from our benchmark. Both aspects consistently revealed a highly similar hierarchy of influence among preprocessing components across iST and sST data.

The difference between two preprocessed data (from the same raw counts) can be quantified by the overlap of their *k*-NN graphs^2,24^ (see Methods “Influence of hyperparameter settings and refinement operations” for more details, Fig. 4a). A higher overlap indicates that the two count-matrix preprocessing workflows yield comparable data structures. For the overall score, the difference between two count-matrix preprocessing workflows is calculated as the absolute difference of their scores. A lower score difference suggests that the two preprocessing strategies yield comparable data quality. We calculated the *k*-NN graph overlap between all preprocessed data for both Slide-seq V2 and Xenium 5K MCH data (Fig. 4b). Interestingly, we discovered a consistent pattern on two data: preprocessings that exhibited a high overlap with the majority of other preprocessings also tended to achieve a higher overall ranking score (Fig. 4c). Specifically, the number of preprocessings with a *k*-NN overlap exceeding a certain number (15 for Xenium 5K, 35 for Slide-seq V2) showed a significant linear correlation with the overall score (r = 0.400 for Xenium 5K, r = 0.424 for Slide-seq V2). This suggests that high-quality preprocessings tend to produce a consensus data structure. A high *k*-NN graph overlap with the majority of preprocessings implies that the effective preprocessing effectively mitigates noise and preserves the intrinsic gene expression manifold of the data, leading to embeddings that are robustly captured across the majority of preprocessings.

**Fig. 4.**
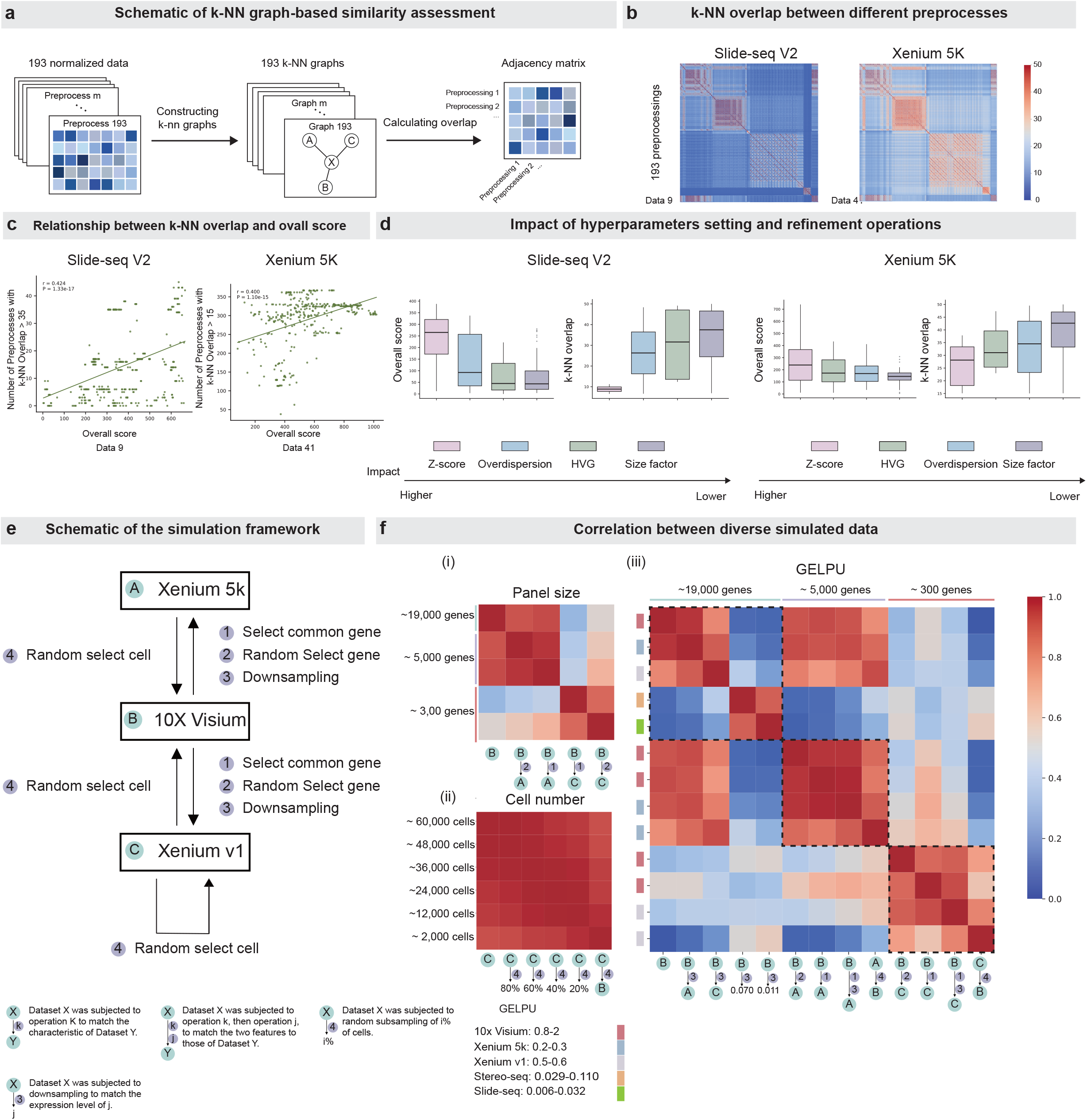
The influence of hyperparameter settings, refinement operations and data structure. **a**, Schematic of *k*-NN graph-based similarity assessment. **b**, Heatmaps of the *k*-NN overlap adjacency matrix for all transformation pairs. **c**, Scatter plot showing the correlation between overall ranking score for each preprocessing and number of preprocessing workflows with *k*-NN overlap > k for each preprocessing (k = 15 for Xenium 5K, k = 35 for Slide-seq V2). Each point represents an individual preprocessing workflow. **d**, Impact of hyperparameters settings and refinement operations. Boxplots show the distribution of the *k*-NN overlap or overall ranking score across different hyperparameters settings and refinement operations. Each box in the plot represents the distribution of minimum *k*-NN overlap change or maximum overall score change after altering only one parameter setting or refinement operation at a time. **e**, Schematic of the simulation framework. **f**, Heatmaps of PCC between benchmark results of different simulated data. On the heatmap, the entry of row i and column j is computed by the PCC of two vectors. One is formed by the overall ranking score of all preprocessings on data i, another is that on data j. The bottom legend of each heatmap denotes the data generation method, and the left-side legend indicates the corresponding data structure. The black dashed box in heatmap (iii) was the correlations between data from same panel size.

Regarding the influence of hyperparameter settings and refinement operations, analysis of the overall score and *k*-NN overlap (Methods) consistently yielded the same conclusions. We calculated minimum k-NN overlap and the maximum overall score change after altering only one parameter setting or refinement operation at a time (Methods). Lower *k*-NN overlap and a higher score gap signify a greater influence. We found that the two aspects yielded consistent conclusions. Across both iST and sST data, Z-score normalization consistently exerted the strongest influence on preprocessing outcomes, whereas size-factor setting showed the weakest influence. The effects of the remaining components, including HVG selection and overdispersion, were intermediate. Importantly, this hierarchy was reproducible across data and was highly consistent between different ST technologies (Fig. 4d, Supplementary Fig. 10).

Therefore, Z-score normalization represents the most influential preprocessing component across both iST and sST data, consistently producing the largest changes in the inferred k-NN graph structure and overall benchmarking performance. Conversely, size-factor setting showed the smallest influence, indicating that moderate changes in size-factor configuration have comparatively limited effects on the resulting data representation. HVG selection and overdispersion showed intermediate effects, and the overall ranking of preprocessing-component influence was consistent across data and ST technologies.

### Panel Size and Gene Expression Level per Unit are Key Drivers of Count-matrix Preprocessing Efficacy

Understanding the influence of inherent data structure on preprocessing performance is important for developing generalizable preprocessing pipelines across diverse ST platforms. Therefore, we investigated the relationship between data structure and preprocessing performance, focusing on three features: the number of cells, gene panel size, and gene expression level per unit (GELPU). We define gene expression level per unit (GELPU) as mean RNA counts captured for each gene within a unit (see Methods “Influence of data structures” for details). For simplified presentation, this data structure will be referred to as “GELPU” throughout the remainder of this text and its accompanying figures.

We generated 17 simulated data from the original three MCH data (10x Visium, Xenium v1, and Xenium 5K) using strategies such as random cell sampling, random gene selection, matched gene selection, and downsampling (Fig. 4e). We then benchmarked the preprocessings across 19 data (17 simulated data and 2 real data) and computed the Pearson correlation coefficient (PCC) of the overall scores (Methods). GSR metric was not utilized for evaluation because, as previously stated, it cannot be assessed with 10x Visium platform.

For panel size, we assessed the PCC between the overall scores of the native 10x Visium MCH data and those of simulated data created by random gene selection and matched gene selection. This comparison ensured that the cell number and GELPU were virtually identical across the compared data. We found that data with identical panel size exhibited highly correlated overall scores, irrespective of the specific genes included. Conversely, a low PCC was observed between data with different panel sizes (Fig. 4f).

For cell number, we followed the same principle of controlled variables: we assessed the PCC between the overall scores of the native Xenium v1 MCH data and those of simulated data created by random cell sampling. We found that data with different cell numbers exhibited a high PCC, suggesting that cell number does not substantially impact preprocessing performance (Fig. 4f).

For GELPU, we followed the same principle of controlled variables, comparing its influence on preprocessing efficacy across three panel sizes: 19,000, 5,000, and 300. In all three scenarios, we first applied gene selection to the native Visium MCH data, followed by downsampling to match the GELPU of Xenium v1 and Xenium 5K. Specifically for the panel size of 19,000 scenario, we further downsampled the 10x Visium MCH data to simulate the GELPU of Array-seq and Slide-seq V2. Our findings revealed that at the panel size of 19,000, data with differing GELPU exhibited substantially lower PCC. This PCC difference weakened for the panel size of 5,000 and became even less pronounced at the panel size of 300 (Fig. 4f).

### Count-matrix preprocessing rankings are robust to upstream image-processing choices

Cell segmentation and transcript assignment directly determine the cell-by-gene count matrix in imaging-based ST data^62,63^. We therefore next investigated whether variation in these upstream procedures substantially alters the relative performance of count-matrix preprocessing workflows. To minimize biological confounding, we constructed a matched benchmark using mouse hippocampal data generated by CosMx, Xenium v1, and Xenium 5K (Supplementary Fig. 13a). We evaluated six segmentation settings, including vendor-provided segmentation, BIDCell^50^, Cellpose^51^, and QuPath WatershedCellDetection^52^ with cell-expansion settings of 0, 5, and 10. We further considered six post-segmentation states, including three transcript-purification strategies (CellAdmix^53^, Ovrlpy^54^, and SPLIT^55^), an unpurified control, and two perturbed conditions representing segmentation error and transcript-assignment noise (Supplementary Fig. 12a, see Methods for more details). For each resulting cell-by-gene matrix, we applied all 371 count-matrix preprocessing workflows and evaluated their performance using the same NC, SF, and GSR framework as in the main benchmark. This factorial design comprised 13,356 distinct combinations of segmentation, post-segmentation processing, and count-matrix preprocessing settings (Fig S12a).

Importantly, although different segmentation and post-segmentation procedures altered cellular boundaries and transcript assignment, the relative performance of count-matrix preprocessing workflows remained highly consistent across upstream processing conditions. Workflows that ranked highly using vendor-provided segmentation generally remained among the top-performing workflows after Cellpose, BIDCell, or WatershedCellDetection segmentation, as well as after transcript purification or simulated-noise perturbations (Supplementary Fig. 12a). We further represented the performance of the 371 workflows under each condition as a preprocessing-ranking profile and quantified the relative contributions of platform, segmentation method, and post-segmentation state to variation in these profiles. Ordination showed substantially clearer grouping by platform than by segmentation method or post-segmentation state (Supplementary Fig. 12b). Consistently, variance partitioning showed that platform explained the largest unique fraction of ranking variation (partial R^2^ = 0.201), compared with segmentation method (partial R^2^ = 0.106) and post-segmentation state (partial R^2^ = 0.103 ) (Supplementary Fig. 12c). Thus, the unique contribution of platform was approximately twice that of either upstream factor. These results indicate that, although upstream image processing influences the composition and purity of the resulting count matrix, it has a substantially smaller effect on the relative preference among count-matrix preprocessing workflows than the underlying ST platform.

The robustness of count-matrix preprocessing rankings, however, does not imply that upstream image-processing procedures themselves perform equivalently. We therefore independently benchmarked segmentation and transcript-purification workflows using three biologically motivated metrics: marker-purity F1 score, negative-marker fraction, and neighborhood contamination^50^ (Supplementary Fig. 12d). Distinct preferences for cell-segmentation strategies were observed across the three platforms: Cellpose and WatershedCellDetection without additional cell expansion performed best for CosMx and Xenium v1, Cellpose and vendor-provided segmentation performed best for Xenium 5K (Supplementary Fig. 12d, Supplementary Fig. 13d). Notably, the favorable performance of Cellpose on Xenium data is consistent with the findings of Marco Salas et al.^6^, who similarly reported that Cellpose outperformed the standard Xenium segmentation in their benchmarking analysis. In contrast, SPLIT consistently achieved the highest overall quality among transcript-purification strategies across all three platforms (Supplementary Fig. 13d). These results demonstrate that upstream image preprocessing should be explicitly optimized, but also support treating upstream image processing and count-matrix preprocessing as complementary but distinct analytical layers in the ST preprocessing workflow.

### Platform- and Data Structure-based Count-Matrix Preprocessing Recommendations

To extract generalized insights from our benchmark results and provide users with actionable and robust guidelines for ST count-matrix preprocessing, we first performed a correlation analysis on the overall scores derived from the initial iST and sST data (Fig. 5a, b). We observed a high PCC between the overall ranking scores of data generated by the same ST platform (average PCC = 0.759; Fig. 5b), but a lower correlation between data from the same tissue profiled using different technologies (average PCC = 0.462; Fig. 5b). These results indicate that data generated by the same ST platform tend to share similar preferences among count-matrix preprocessing workflows.

**Fig. 5.**
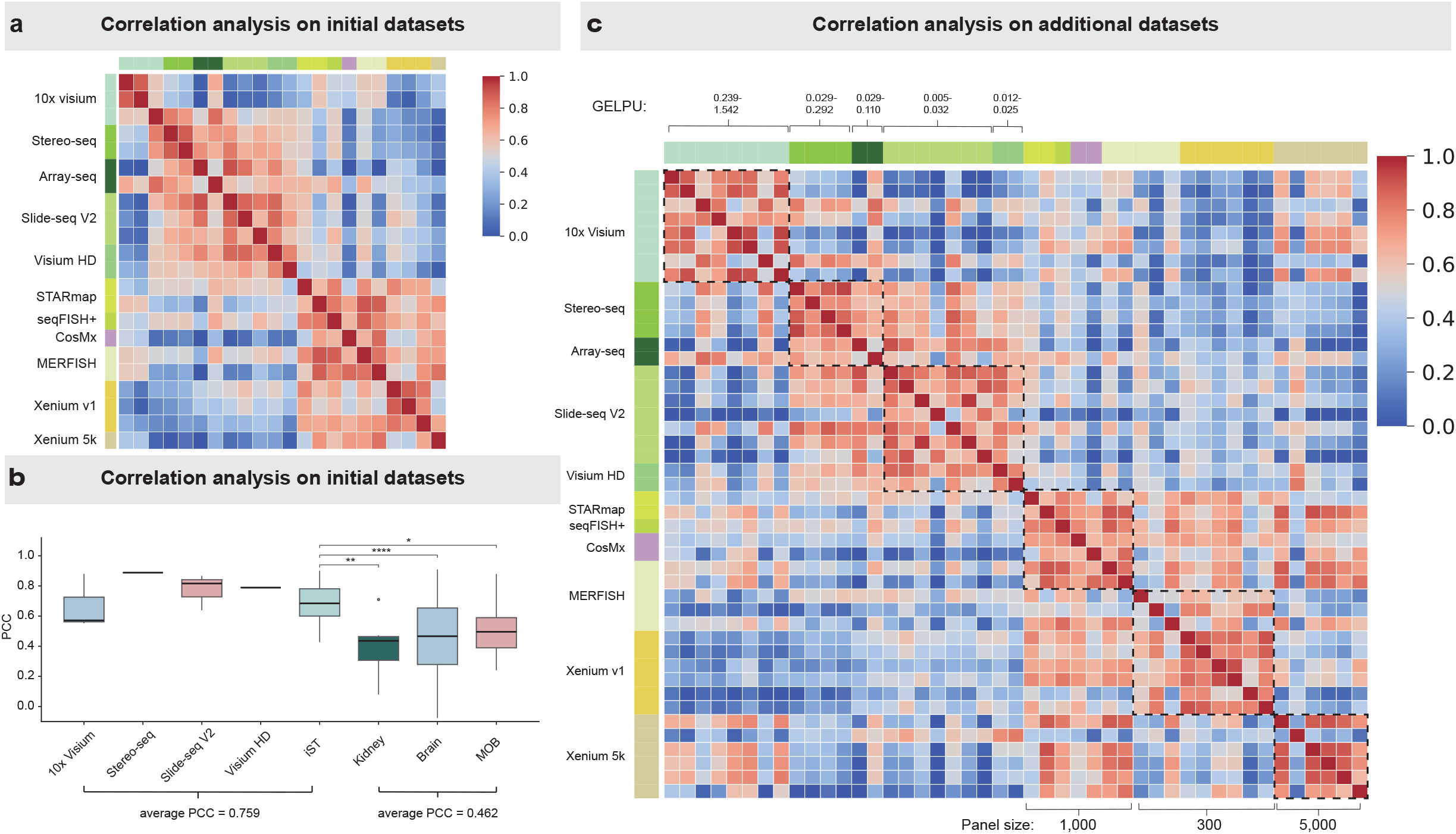
Correlation analysis of benchmark results. **a**, Heatmap of PCC between benchmark results of initial data (data from MCH, MOB and MKD, see Fig. 1d). The left-side legend of the heatmap indicates the corresponding ST platform. **b**, Boxplot of PCC between benchmark results within data groups. Each box represents the PCC between benchmark results of data from the data groups indicated by the x-axis label. **c**, Heatmap of PCC between benchmark results of all data included in the study (see Supplementary Fig. 1). The left-side legend of the heatmap indicates the corresponding ST platform. The black dashed box was the correlations between data that was from the same platform or has similar data structure. Our subsequent preprocessing recommendations were derived from the benchmark results of the data grouped within each black dashed box.

To further evaluate the generalizability of this pattern, we extended the benchmark to 45 data spanning diverse tissue types, species, disease states, and ST technologies. Across this expanded collection, data generated by the same platform generally exhibited highly correlated count-matrix preprocessing-ranking profiles, whereas substantial differences were observed between platforms (Fig. 5c). The MERFISH platform represented a notable exception due to its flexible panel size configuration. MERFISH data with a 1,000-gene panel exhibited high PCCs with seqFISH+, CosMx and STARmap data with similar panel sizes, whereas 300-gene MERFISH data showed substantially different preprocessing-ranking profiles (Fig. 5c). Similarly, among whole-transcriptome sST data, platforms with comparable GELPU, such as Array-seq and Stereo-seq, showed similar preprocessing preferences. These observations further suggested that gene panel size and GELPU, in addition to platform, are important determinants of count-matrix preprocessing performance.

We further quantified these relationships through variance-partitioning analysis using platform, tissue, species, gene panel size, GELPU, and cell number as explanatory variables (Supplementary Fig. 11a). In single-predictor models, platform showed the strongest association with variation in preprocessing-ranking profiles, followed by gene panel size and GELPU, whereas cell number showed only a minor effect (Supplementary Fig. 11d). This pattern independently recapitulated our controlled data-structure experiments, in which panel size and GELPU strongly influenced preprocessing performance whereas cell number had comparatively little impact. When all variables were considered simultaneously, platform retained the largest unique contribution (Supplementary Fig. 11d); the contributions of panel size and GELPU were reduced because part of their explanatory variation was shared with platform. Together, the controlled experiments and variance-partitioning analyses consistently identify platform, gene panel size, and GELPU as the major factors associated with count-matrix preprocessing preferences, with platform capturing the strongest overall technology-specific effect.

Based on these results, we established a hierarchical recommendation framework integrating platform identity and intrinsic data structure (Fig. 6). If ST data are generated by a platform represented in our benchmark, we recommend platform-based selection of count-matrix preprocessing workflows. For MERFISH, which exhibits substantial variation in panel size, recommendations are further stratified into 300-gene and 1,000-gene panel groups. If the ST platform is not represented in our benchmark, we recommend data structure-based selection: for whole-transcriptome data, workflows should primarily be selected according to GELPU, whereas for targeted or limited-panel data, panel size provides the primary matching criterion. Accordingly, we defined seven recommendation groups based on platform or data-structure characteristics. Rather than selecting a single workflow as uniquely optimal, we recommend the top 20 count-matrix preprocessing workflows within each group as a candidate set, reflecting the similar performance of multiple highly ranked workflows while providing users with a robust range of preprocessing choices (Fig. 6).

**Fig. 6.**
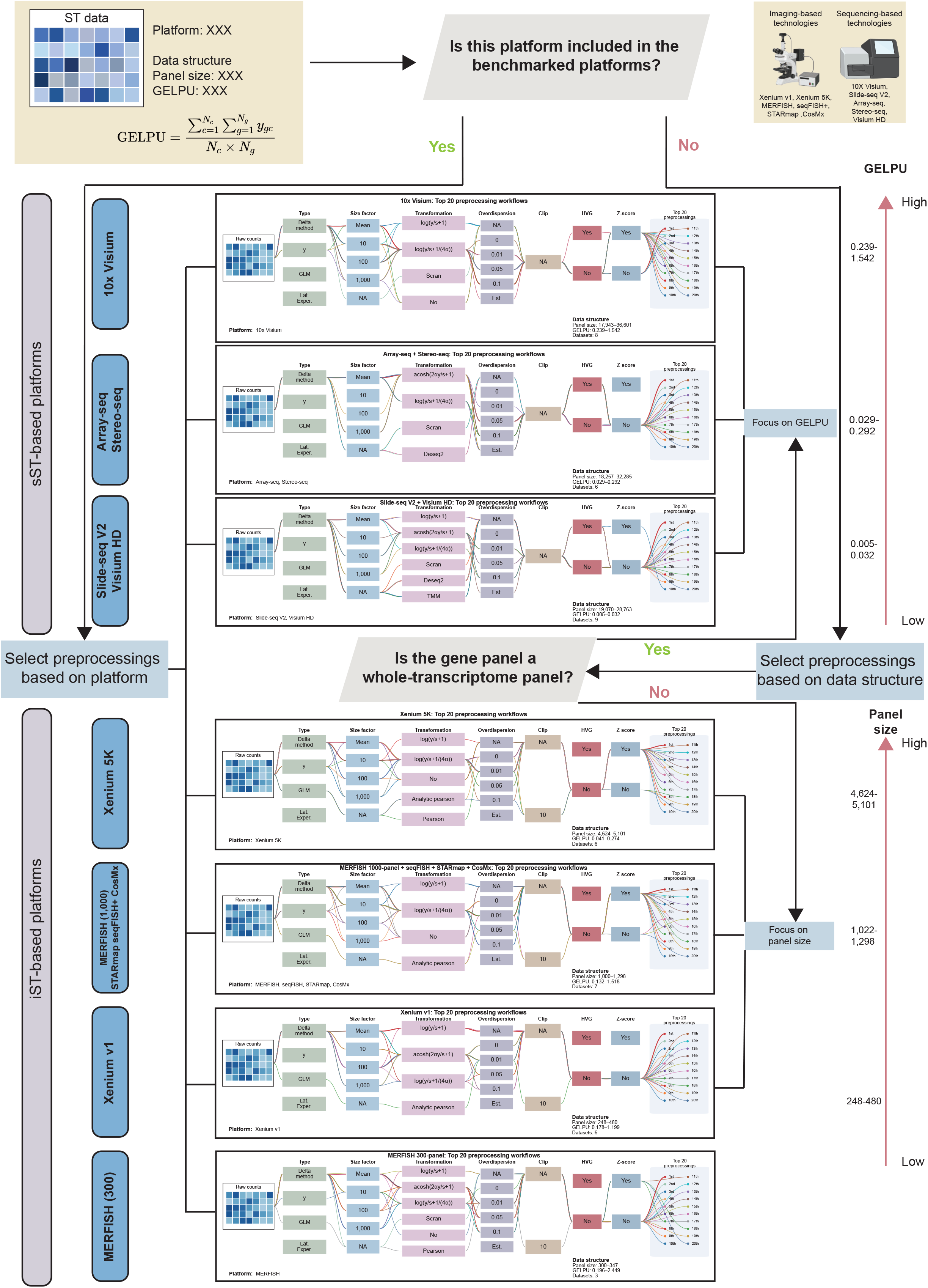
A practical guideline for preprocessing. A step-by-step guideline for users to select optimal preprocessing workflow. We provide seven recommended preprocessing sets, each containing 20 specific preprocessing workflows. For any ST data requiring preprocessing, users should begin by evaluating the data’s critical characteristics, including its originating platform, panel size, and GELPU. For GELPU calculation in figure, *y*_*gc*_ is the element in the row *g* and column *c* of raw counts matrix, *N*_*g*_ and *N*_*c*_ are the panel size and cell number of raw counts matrix. If the ST data platform is included in our benchmark study, we advise selecting the optimal preprocessing workflow corresponding to that specific platform’s best-performing set. If the platform is not included, the selection should be based on the data’s structure: for a whole-transcriptome panel, choose the workflow based on the data’s GELPU; otherwise, for limited panels, select the scheme based on the data’s panel size.

We next examined the robustness of these recommendations across the complete benchmark. Visualization of all 371 count-matrix preprocessing workflows across all 45 data revealed substantial between-platform variation but pronounced within-platform consistency in preprocessing preferences (Supplementary Fig. 22). No single count-matrix preprocessing workflow consistently performed well across all data and platforms; workflows that ranked highly for one platform could perform substantially worse on another. In contrast, independent data generated by the same platform showed highly similar ranking patterns, and the recommended workflows consistently remained within the top-performing range across data corresponding to their target platforms. These results simultaneously demonstrate the absence of a universally optimal count-matrix preprocessing workflow and the reproducibility of platform-specific preprocessing preferences across independent data.

The stability of highly ranked workflows was further confirmed under several perturbation settings. Across Xenium v1 and Xenium 5K data spanning different tissues and species, workflows with high platform-level aggregate rankings consistently showed the highest frequencies of remaining among the top-performing workflows (Supplementary Fig. 23). Similarly, after independently removing 10% of cells/spots or 10% of genes, highly ranked workflows generally remained within the top-performing range (Supplementary Fig. 24, 25). Their relative performance was also stable when varying key evaluation parameters, including the number of nearest neighbors, the number of principal components, and the distance metric (Supplementary Fig. 26, 32). Although the exact ordering among closely performing workflows changed modestly under these perturbations, the separation between consistently high- and low-performing workflows remained substantially more stable.

Together, these analyses support a practical recommendation framework in which platform serves as the primary criterion for technologies represented in our benchmark, while gene panel size and GELPU provide complementary guidance for data with atypical structures or from emerging and unbenchmarked platforms. The absence of a universally high-performing workflow, together with the reproducible performance of the recommended Top-20 workflow sets across data and perturbations, supports the robustness of these platform- and data structure-specific recommendations.

### Recommended Count-Matrix Preprocessing Workflows Excel Across Diverse Downstream Tasks

To rigorously assess the generalizability and practical utility of our proposed count-matrix preprocessing recommendations, we evaluated their performance across four representative downstream tasks in ST analysis: cell type identification, spatial domain identification, spatially variable gene (SVG) detection, and marker recovery. These tasks span multiple levels of biological information, including cell identity, tissue organization, spatial gene-expression patterns, and gene-expression relationships.

For cell type identification, we collected two data from MERFISH and Xenium v1 platforms and then quantitatively and qualitatively demonstrated that our recommended count-matrix preprocessing workflows significantly improve performance on this task (Fig. 7b). The first data, derived from human cortical areas at gestational week 20 (GW20)^25^ (Fig. 7a), was generated using the MERFISH platform. In this data, Qian et al. identified a distinct cell type composition demarcating the border between the primary visual cortex (V1) and secondary visual cortex (V2)^25^. To evaluate the performance of our recommended count-matrix preprocessing workflows for cell type identification, we applied the top three and bottom three count-matrix preprocessing workflows, which were selected based on their overall score across the 300-panel size MERFISH data (Fig. 7a). Subsequently, unsupervised clustering via the Leiden algorithm^61^ was performed to reveal the cell type distributions (Fig. 7a). We observed that while all six count-matrix preprocessing workflows yielded data successfully capturing the prominently layered structure characteristic of the cerebral cortex, their ability to define the subtle V1/V2 boundary varied substantially (Fig. 7c). Specifically, only the data processed by the top three workflows clearly resolved the distinct V1/V2 border (Fig. 7c). The bottom three workflows failed entirely, as no discernible V1/V2 difference was evident in their resulting cell type distributions (Fig. 7c). This V1/V2 border remained unidentifiable even at higher clustering resolutions (Supplementary Fig. 29a). The findings demonstrate that count-matrix preprocessing is a critical determinant for resolving fine biological structures in ST data. In particular, the improved resolution of the V1/V2 boundary indicates that highly ranked count-matrix preprocessing workflows better preserve subtle biological differences in cellular states.

**Fig. 7.**
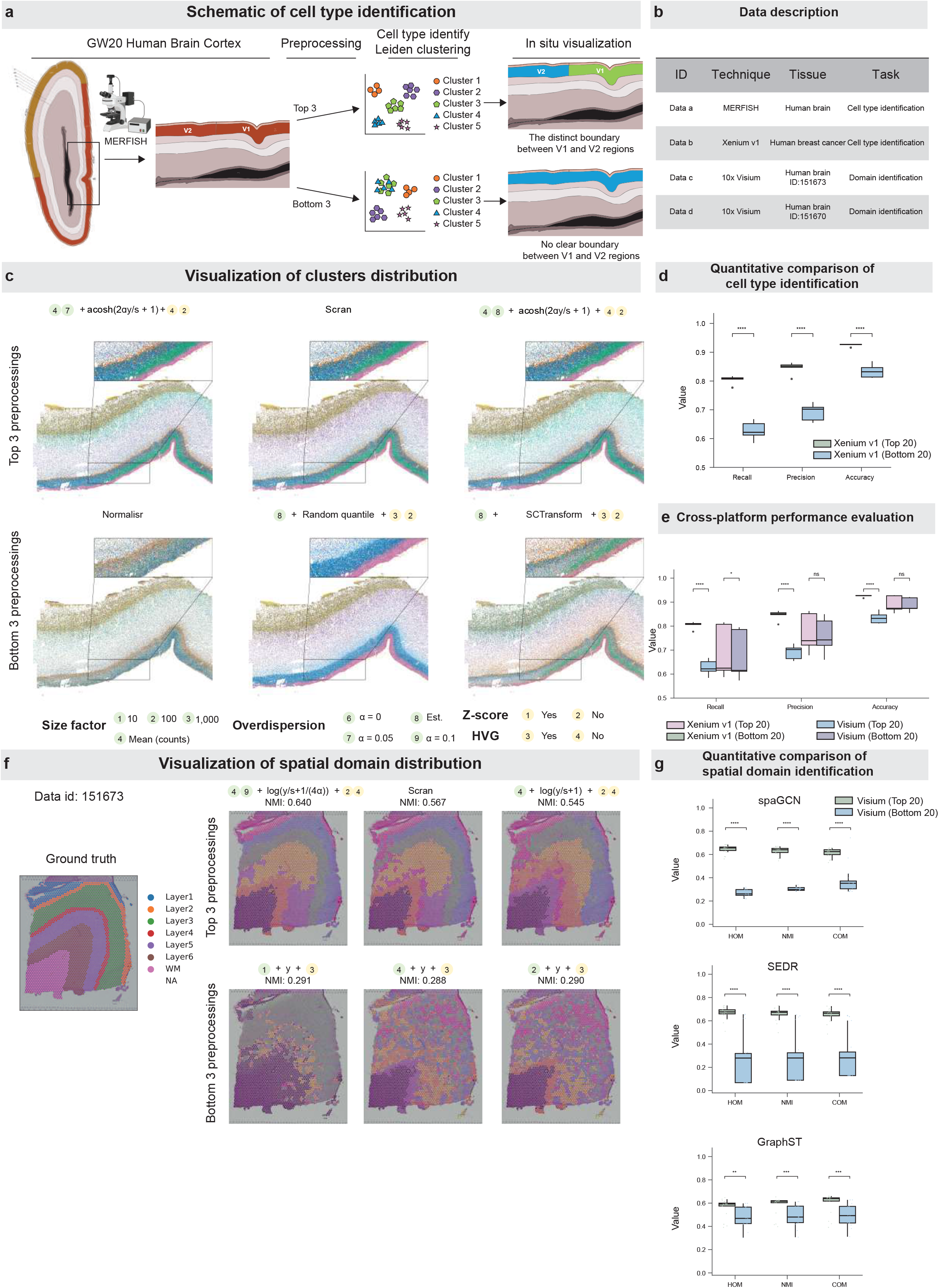
Validation on Downstream Task. **a**, The pipeline of the validation workflow on MERFISH GW20 human brain cortex data. Schematics are taken from the Allen Reference Atlas for GW21 human fetal brain^49^. Some elements in this figure are sourced from BioRender. **b**, Description of data used in this section. **c**, Visualization of cell clustering results obtained with the top 3 preprocessings and bottom 3 preprocessings, respectively, as determined by their overall score on the MERFISH (300-panel size) data (Data 27 to Data 29). **d**, Quantitative comparison of cell type identification. The grouped boxplot shows the distribution of metric values obtained through 5-fold cross-validation. Within each group, the two columns represent the corresponding metric scores achieved on this task by the top 20 and bottom 20 preprocessings, respectively, as determined by their overall score on the Xenium v1 data (Data 30 to Data 35). **e**, Cross-Platform performance evaluation. The grouped boxplot shows the distribution of metric values obtained through 5-fold cross-validation. Within each group, the four columns represent the corresponding metric scores achieved on this task by the Xenium v1 top 20, Xenium v1 bottom 20, 10x Visium top 20, and 10x Visium bottom 20 preprocessings, respectively, as determined by their overall score on the Xenium v1 data (Data 30 to Data 35) and 10x Visium data (Data 1 to Data 8). **f**, Visualization of the spatial domain clustering results on Data c (Fig. 7b) achieved using the SpaGCN, comparing the outcomes obtained with the top 3 and bottom 3 performing preprocessings, respectively, as determined by their overall score on the 10x Visium data (Data 1 to Data 8). **g**, Quantitative comparison of spatial domain identification. The grouped boxplot shows the distribution of metric values through quantifying the agreement between the spatial cluster labels obtained by each algorithm and the ground truth labels. Within each group, the two columns represent the corresponding metric scores achieved on this task by the 10x Visium top 20 and 10x Visium bottom 20 preprocessings.

Next, we quantitatively evaluated the recommended count-matrix preprocessing workflows using an externally annotated data with cell type labels as ground truth. This validation data, generated by the Xenium v1 platform, was derived from human breast cancer^26^ (Fig. 7b). Crucially, this data included annotated cell labels that served as the reliable ground truth for validation. To assess the performance of each count-matrix preprocessing workflow, we trained a support vector machine (SVM) for multiclass cell type classification on each preprocessed data and measured its performance on an independent test set. To obtain robust performance metrics, we implemented five-fold cross-validation^27^ (Methods). We quantified the discrepancies between the predicted and true cell labels using three standard metrics: recall, precision, and accuracy (Methods). We found that the top 20 count-matrix preprocessing workflows, based on their overall ranking score across the Xenium v1 data (Data 30 to Data 35), significantly outperformed the bottom 20 workflows across all three metrics: recall, precision, and accuracy^28^ (Fig. 7d). These quantitative results further demonstrate that highly ranked count-matrix preprocessing workflows better preserve information relevant to cell type identification.

Intriguingly, when we evaluated the performance of the best and worst 20 count-matrix preprocessing workflows derived from the 10x Visium data (Data 1 to Data 8) on this Xenium data, we observed a striking reversal (Fig. 7e): the 20 worst workflows from the 10x Visium ranking actually outperformed its 20 best workflows across the three metrics (recall, precision, and accuracy). This finding indicates that count-matrix preprocessing preferences derived from one ST platform do not necessarily transfer to another platform. Consequently, our analysis highlights the critical conclusion that no universal count-matrix preprocessing workflow can effectively handle all types of ST data. These results further support the importance of selecting count-matrix preprocessing workflows according to the characteristics of individual ST platforms rather than directly transferring commonly used workflows across technologies.

For spatial domain identification, we focused on 10x Visium human postmortem tissue sections data (DLPFC)^29^ and then quantitatively and qualitatively demonstrated that our recommended count-matrix preprocessing workflows significantly improve performance on this task. This data is the most widely used benchmark data among spatial clustering methods and contains 12 tissue sections from three donors. We validated the performance of the count-matrix preprocessing workflows on two sections from two distinct donors (Fig. 7b). We employed a suite of popular spatial domain identification methods, including SpaGCN^30^, GraphST^31^, and SEDR^32^ (Methods). Across all tested methods, the results consistently demonstrated a clear performance disparity. Specifically, the top 20 count-matrix preprocessing workflows (based on the average overall score across all tested 10x Visium data) significantly outperformed the bottom 20 workflows across all metrics: completeness score (COM), normalized mutual information (NMI), and homogeneity score (HOM) (Fig. 7g, Supplementary Fig. 29c, Methods)^33^. Importantly, the superior performance of the top-ranked count-matrix preprocessing workflows remained stable across a broader range of hyperparameter settings for SpaGCN, SEDR, and GraphST (Supplementary Fig. 33). Furthermore, we visualized the SpaGCN spatial domain results generated by the top three and bottom three count-matrix preprocessing workflows, clearly showing the enhanced performance of the top three workflows (Fig. 7f, Supplementary Fig. 29b). Together, these results demonstrate that highly ranked count-matrix preprocessing workflows improve the recovery of biologically meaningful tissue-domain organization across multiple spatial clustering approaches.

We next evaluated the performance of count-matrix preprocessing workflows in SVG detection, an important downstream task for identifying genes with spatially structured expression patterns. We compared the top 30 and bottom 30 count-matrix preprocessing workflows across five representative data generated by Array-seq, Slide-seq V2, Stereo-seq, Visium HD, and Xenium 5K. For each count-matrix preprocessing workflow, SVGs were first identified using the complete data. We then randomly removed 20% of the cells or spots, repeated SVG detection on the perturbed data, and quantified SVG stability by calculating the Jaccard similarity between the top 10% SVG sets identified before and after perturbation (Methods). To ensure that the observed trends were not dependent on a single SVG detection method, we performed the analysis using both Moran’s I and SPARK-X. Across nearly all platform–method combinations, the top-ranked count-matrix preprocessing workflows showed higher SVG stability than the bottom-ranked workflows (Supplementary Fig. 30a). These results indicate that highly ranked count-matrix preprocessing workflows better preserve reproducible spatial gene-expression patterns under moderate perturbation of the input data.

Finally, we evaluated whether count-matrix preprocessing workflows preserved gene-expression relationships using a marker-recovery task. We compared the top 30 and bottom 30 count-matrix preprocessing workflows across data generated by CosMx, MERFISH, Slide-seq V2, Stereo-seq, Visium HD, Xenium v1, and Xenium 5K. For each data, the top 20 highly variable genes were selected as target genes for recovery, and their expression values were completely masked from the preprocessed expression matrix before feature construction. PCA was then performed using the remaining genes, and the resulting principal components were used to predict the binary expression status (detected versus undetected) of each target gene in individual cells or spots. Cells/Spots were randomly divided into equally sized training and test sets, with 50% used for model training and 50% held out for evaluation. The same train/test partition was maintained across all count-matrix preprocessing workflows. To reduce dependence on a particular prediction model, we evaluated three classifiers—k-nearest neighbors, logistic regression, and a multilayer perceptron (MLP)—and quantified prediction performance using AUROC for each target gene and macro-AUROC across the 20 target genes. Across data and classifier architectures, top-ranked count-matrix preprocessing workflows consistently achieved higher marker-recovery performance than bottom-ranked workflows (Supplementary Fig. 30b). These results demonstrate that highly ranked count-matrix preprocessing workflows more effectively preserve gene-expression relationships that support the recovery of masked gene signals.

Overall, highly ranked count-matrix preprocessing workflows consistently showed superior performance across four complementary downstream tasks, encompassing cell type identification, spatial domain identification, SVG detection, and marker recovery. These analyses demonstrate that count-matrix preprocessing quality has broad consequences for the preservation and recovery of biologically meaningful information at both the cellular and gene-expression levels. Together, the consistent advantages of highly ranked count-matrix preprocessing workflows across diverse platforms and analytical tasks support the practical utility and robustness of our platform-specific count-matrix preprocessing recommendations for downstream ST analysis.

## Discussion

In this study, we performed a comprehensive benchmark analysis of 371 count-matrix preprocessing workflows. Our framework spanned the spectrum of current count-matrix preprocessing workflows, encompassing mainstream transformations, varying hyperparameter settings, and crucial refinement operations. To ensure the robust generalizability of our findings while minimizing the influence of confounding factors, we curated an extensive collection of data, including data derived from 11 different ST platforms across diverse tissue types and biological contexts. We established a comprehensive evaluation framework incorporating metrics for NC, SF, and GSR, with a focus on assessing the quality of the fundamental cell graph structure at both local and global levels. Our analysis demonstrates that no single count-matrix preprocessing workflow consistently performs optimally across all ST data. Although delta method-based transformations consistently showed strong performance across different ST platforms, we found that the optimal hyperparameter configurations and specific refinement operations differ significantly between platforms. We further quantified the impact of parameter choices and refinement operations and found that Z-score normalization exerted the largest overall influence on preprocessing performance, whereas size-factor setting showed the smallest influence across both iST and sST data. Data-estimated overdispersion generally performed less favorably than fixed settings across platforms. We also identified platform, gene panel size, and GELPU as major factors associated with preprocessing preference, with panel size and GELPU representing key intrinsic data-structure characteristics. We then collected and benchmarked a more comprehensive range of data to expand the generalizability of our findings. Based on these extended analyses, we provide users with actionable and robust guidelines tailored to specific ST platforms and data structures, with the top-performing workflows presented as recommended candidate sets rather than a single universally optimal solution (Fig. 6). Finally, we performed extensive validation across cell type identification, spatial domain identification, SVG detection, and marker recovery, consistently showing that highly ranked count-matrix preprocessing workflows improve performance across diverse downstream tasks.

Our study addresses a critical gap in ST count-matrix preprocessing by establishing a comprehensive benchmark and offering detailed preprocessing guidelines tailored to different ST technologies and data structures. These guidelines are designed to directly assist researchers in achieving more accurate and reliable downstream analyses.

Beyond the strategic recommendations, we have curated and released a valuable resource: a collection of ST data derived from identical tissue types measured by different ST platforms, which serves as an essential resource for future benchmarking studies. In addition, to facilitate the calculation of GSR metrics, we gathered matched scRNA-seq data for the iST data included in our benchmark, which subsequently supports common downstream analytical tasks, such as accurate cell type identification. This rich collection of readily usable resources will significantly benefit the wider research community.

Furthermore, the results of our comprehensive benchmarking are highly generalizable. Our data structure analysis reveals that gene panel size and GELPU are key intrinsic data characteristics influencing count-matrix preprocessing efficacy. This conclusion was supported by both controlled data-structure perturbation experiments and variance-partitioning analyses across real data. Together with the strong platform-associated variation observed across data, these results provide a practical basis for selecting count-matrix preprocessing workflows according to platform when available, or according to panel size and GELPU for emerging or unbenchmarked technologies.

These findings are also relevant to emerging spatial foundation models^64,65^, multimodal learning frameworks^66–68^, and AI-based ST analysis pipelines^69^. Many such approaches rely on transcriptomic features or count-derived representations as model inputs, making the quality and consistency of count-matrix preprocessing important for subsequent representation learning and multimodal integration^70,71^. Our platform- and data structure-aware recommendations therefore provide a model-agnostic starting point for constructing transcriptomic inputs across heterogeneous ST technologies^72–74^. Moreover, the recommended sets of consistently high-performing workflows can substantially reduce the preprocessing search space for computationally intensive AI models and provide practical candidates for sensitivity analyses, allowing model development to focus on representation learning and downstream optimization rather than exhaustive preprocessing searches. The same principle also applies to multiview graph-based spatial-domain models that use gene-expression-derived features^44^.

Our benchmarking also serves to inspire the development of novel preprocessing strategies and downstream analysis models for ST. Specifically, our results consistently demonstrate that raw count data remains highly informative, and when combined with strategic refinement steps, it achieves notably strong performance. This suggests that effective ST preprocessing does not necessarily require a global variance-stabilizing transformation in all settings. Future research should also explore developing count-based models that can directly handle raw counts, thereby avoiding potential information loss and statistical artifacts introduced by traditional variance-stabilizing transformations.

## Methods

### 1. Preprocessing

Our preprocessing framework consisted of three stages: setting hyperparameters, transformation, and refinement operations. The following sections detail each of these steps.

#### 1.1 Stage 1: Setting hyperparameters

##### Size factors

The size factors, often referred to as library size, originated in scRNA-seq preprocessing with the primary goal of removing technical artifacts stemming from variable RNA capture efficiency and sequencing depth among individual cells^15^. However, the application of size factor normalization in ST is highly debated due to the confounding effect of varying cell density or spot size^3,4^. The current definitions of size factors mainly fall into two categories: counts-based and volume-based. Conventionally, the counts-based size factor *s*_*c*_ ∈ ℝ^+^ for an individual cell or spot *c* is defined as:

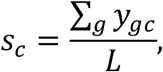

where *y*_*gc*_ ∈ ℕ is the expression count of gene *g* in cell *c*, the numerator is the total counts of the cell *c* and *L* ∈ ℝ^+^ is a constant. In our benchmark, we tested various choices for the constant *L*. Specifically, we set *L* to 10, 100, 1000, or the overall mean total count per cell. The volume-based size factors are defined as:

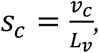

where *v*_*c*_ ∈ ℝ^+^ is the volume of cell *c* and *L*_*v*_ ∈ ℝ^+^ is the mean volume of all cells. Then the raw counts are divided by size factors to account for differences in RNA yield or cell volume among cells.

##### Overdispersion

The concept of overdispersion originates from the statistical modeling of scRNA-seq data^16-18^, where gene expression variance is significantly greater than the mean. Given this characteristic, many studies advocate for modeling scRNA-seq counts using the Negative Binomial (NB) distribution, and the overdispersion is an important parameter in this distribution (Methods). This statistical foundation has led to the development of several NB-based transformation methods^9,12,13^. In the NB distribution, the overdispersion accounts for the additional, higher variance beyond what is predicted by the Poisson model. The mean-variance relationship in the NB distribution is defined as:

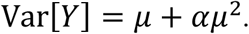

Here, *Y* ∈ ℕ represents the random variable, *μ* ∈ ℝ^+^ is the mean value for *Y*, Var[*Y*] ∈ ℝ^+^ is the variance of the *Y*, and *α* ∈ ℝ^+^ is the overdispersion in NB distribution. When the *α* is 0, the NB distribution degenerates into the Poisson distribution.

#### 1.2 Stage 2: Transformation

The second stage is transformation. In this study, we investigated 12 widely used transformation methods, which can be grouped into three approaches (delta method, GLM residuals, and latent expression models). The primary objectives of these transformations are twofold. First, they involve the normalization of counts using the predetermined size factors^2,3,12^. Specifically, some delta method-based transformations directly divide raw counts by size factors, unifying the total counts across cells^34-36^. In contrast, GLM-based methods model the size factors as covariates, integrating them directly into the statistical framework to account for technical variation^9,12^. Second, transformation aims to address heteroscedasticity, the confounding phenomenon where the variance of gene expression is highly dependent on its abundance^2,12^. Specifically, these transformations seek to decouple the variance of gene from its abundance. This ensures that the biological signal from both lowly and highly expressed genes can contribute to the definition of cellular state.

##### The delta method

The delta method is a mathematical technique used to approximate the variance of a transformed random variable^11^. If the variance Var[*X*] and mean *μ* of a random variable *X* are known, the variance of the transformed random variable *g*(*X*) ∈ ℝ can be approximated by:

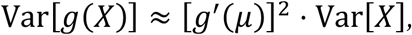

where *g*^′^ (*μ*) is the derivative of the function *g*(·) evaluated at *μ*. The delta method resolves the variance stabilization issue by applying transformation *g*(·) to random variable *X*, resulting in a transformed variable *g*(*X*) with approximately constant variance. This is possible because the random variable *X*, when following a NB distribution, is known to exhibit a functional relationship between its variance Var[*X*] and its mean *μ*. This relationship is typically expressed as:

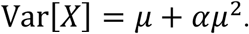

We can set the variance of the transformed variable to a constant, SD[*g*(*X*)] = *const*., then the required derivative of the variance-stabilizing transformation *g*(·) can be derived using the delta method:

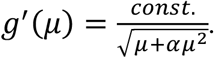

Finally, the transformation *g*(·) can be obtained through integration. Using this approach, Ahlmann-Eltze et al.^2^ derived the transformation 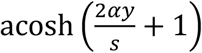 and 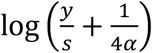.

In our benchmark, the delta method-based transformations were: 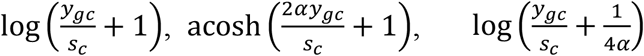, Scran, TMM^37^ and Deseq2^34^. For these transformations, *y*_*gc*_ ∈ ℕ is the expression count of gene *g* in cell *c, s*_*c*_ ∈ ℝ^+^ denotes size factor of cell *c*, and *α* ∈ ℝ^+^ denotes overdispersion. The 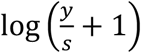 is the most commonly used transformation^35,36^. The 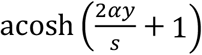 and 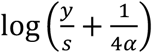 were suggested by Ahlmann-Eltze et al^2^. These transformations can be implemented in the transformGamPoi package^2^. The Scran, TMM and Deseq2 were originally developed for bulk RNA and scRNA-seq data, and are commonly adapted for ST data preprocessing. They follow the shifted logarithm 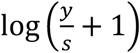, but estimate the size factors based on different ways.

The Scran transformation leverages a deconvolution approach to estimate the size factors based on a linear regression over genes for pools of cells, and is implemented in the Scran package^15^. The TMM (Trimmed Mean of M-values) transformation estimates size factors by calculating a trimmed mean of expression ratios between each cell and a reference cell, and is implemented in the edgeR package^38^. The Deseq2 transformation estimates the size factors using the median of ratios method, and is implemented in the DESeq2 package^34^.

##### GLM residuals

Hafemeister and Satija modeled the raw counts using a NB generalized linear model^12^,

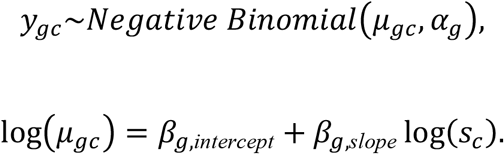

Here, *y*_*gc*_ ∈ ℕ is the raw counts of cell *c* and gene *g, s*_*c*_ ∈ ℝ^+^ is the size factor of cell *c, α*_*g*_ ∈ ℝ^+^ is the overdispersion of gene *g*, and *β*_*g,intercept*_ ∈ ℝ and *β*_*gc,slope*_ ∈ ℝ are intercept and slope parameters for gene *g*. They then calculated the Pearson residuals from this model:

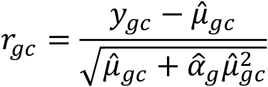

where 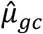 and 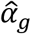 are estimated parameters derived from the NB model. The *r*_*gc*_ are the resulting transformed data.

In our benchmark, GLM-based transformations were: Pearson, SCTransform, Analytic Pearson and Random Quantile. The Pearson residuals are calculated directly as *r*_*gc*_, implemented in the transformGamPoi package. The Pearson residuals are initially calculated as *r*_*gc*_. Each residual is then clipped to be within 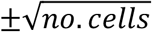, a step suggested by Hafemeister and Satija^12^. The SCTransform also utilizes Pearson residuals but incorporates additional heuristics, and is implemented by the SCTransform package^12^. The Analytic Pearson represents an analytic approximation of the Pearson residuals, suggested by Lause et al., and is implemented by the transformGamPoi package^9^. The Random Quantile uses a randomization technique to convert the data into standard normal quantiles, resulting in transformed data that more closely approximates a normal distribution, and is implemented by the transformGamPoi package^2^.

##### The latent expression models

These transformations infer the parameters of a postulated generative model, aiming to estimate latent gene expression values based on the observed counts. Two prominent examples of this approach are Sanity^13^ and Normalisr^14^. Sanity specifically is a Bayesian normalization procedure derived from first principles that estimates expression values and associated error bars directly from raw scRNA-seq UMI counts without any tunable parameters. We apply this method to ST datasets preprocessing to empirically assess its performance. Normalisr is primarily a tool designed for frequentist hypothesis testing. However, because it infers logarithmic latent gene expression, it can also serve as a generic preprocessing method. Normalisr returns the minimum mean square error estimate for each count, assuming an underlying binomial generative model. It is implemented by the Normalisr package.

##### Spatially aware normalization

SpaNorm was considered directly relevant to our benchmark because it explicitly incorporates spatial information during normalization. However, SpaNorm showed limited scalability for the large ST datasets included in our benchmark. On Data 34 (∼300,000 cells), a single SpaNorm run did not complete within 6 h under our computational environment. Because the benchmark included 45 datasets, several containing hundreds of thousands to nearly one million cells/spots, and each normalization method needed to be evaluated together with multiple refinement settings and benchmarking metrics, a complete SpaNorm evaluation was computationally prohibitive. We therefore excluded SpaNorm from the final quantitative benchmark.

#### 1.3 Stage 3: Refinement operations

The third stage is refinement operations, which includes two critical operations: highly variable gene (HVG) selection, Z-score normalization and clipping.

##### HVG selection

HVG selection acts as a form of initial dimensionality reduction after transformation^39^. We computed the variance of each gene across cells in the transformed data. Genes were then selected as HVGs if their variance fell within the top 20% of all calculated variances.

##### Z-score normalization

Z-score normalization was evaluated as an optional refinement after HVG selection^40^. This operation scales the expression of each gene, where its variance becomes 1, and its mean becomes 0. Crucially, this standardization forces all genes to contribute equally to the definition of the cellular state. We take the transformed expression values *x*_*gc*_ = *g*(*y*_*gc*_ ), and compute the Z-score *z*_*gc*_ ∈ ℝ for gene *g* in cell *c*. The Z-score *z*_*gc*_ is defined by the formula:

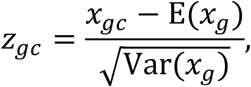

where mean E(·) and variance Var(·) are the empirical mean and variance taken across cells. Final Z-scores are then clipped to a maximum absolute value 10 to mitigate the influence of extreme outliers after scaling.

##### Clipping

For Pearson-residual-based transformations, residual clipping was evaluated as an additional refinement to limit the influence of extreme residual values. Given a Pearson residual *r*_*cg*_ and a clipping threshold *c*, the clipped residual was defined as

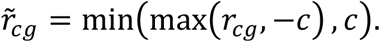

We explicitly evaluated multiple residual-clipping strategies, including no clipping, a fixed clipping threshold of 10, and two additional clipping thresholds included in the benchmark. No clipping retained the original residual values, whereas finite clipping thresholds truncated residuals whose absolute values exceeded the specified threshold.

### 2. Image preprocessing

We evaluated cell segmentation and post-segmentation transcript processing on matched mouse hippocampal datasets generated by three imaging-based spatial transcriptomics platforms: Xenium v1, Xenium 5K, and CosMx. For each platform, six segmentation settings were considered: the platform-provided vendor segmentation, Cellpose, BIDCell, and QuPath WatershedCellDetection with cell-expansion distances of 0, 5, or 10 µm. Each segmentation result was retained as an unpurified baseline and further processed with three transcript-purification methods (CellAdmix, Split, and Ovrlpy) or two simulated perturbations representing segmentation error and transcript-assignment noise. The resulting 36 image-processing variants per platform (108 variants in total) were used both to evaluate upstream image-processing quality and to test the robustness of the 371 count-matrix preprocessing workflows. The full factorial analysis comprised 6 segmentation settings × 6 post-segmentation states × 371 count-matrix preprocessing workflows = 13,356 processing combinations, evaluated separately for each of the three platforms.

#### 2.1 Input standardization and spatial coordinates

All data variants were represented as AnnData/H5AD objects containing a non-negative cell-by-gene raw-count matrix, read preferentially from layers[‘counts’] and otherwise from X, and two-dimensional global cell centroids in obsm[‘spatial’]. Non-vendor segmentations used obs[‘seg_label’] for mask labels, whereas vendor data retained the native cell_id. For Xenium, transcripts with quality value <20 and negative-control, blank, antisense, Unassigned, or Deprecated features were excluded. For CosMx, only transcripts with codeclass = Endogenous were retained.

Transcript coordinates were transformed to a common global coordinate system before assignment. Pixel sizes were 0.2125 µm/pixel for Xenium, 0.1203 µm/pixel for CosMx, and 1.0 µm/pixel for BIDCell masks. CosMx field-of-view origins were obtained from latest.fovs.csv, and local y coordinates were transformed consistently with image stitching.

#### 2.2 Cell segmentation and transcript quantification

##### Vendor segmentation

Platform-provided cell boundaries and transcript-to-cell assignments were used directly as the native segmentation baseline. Native cell identifiers were retained to restore spatial coordinates and cell-area metadata after purification and to support paired comparisons.

##### Cellpose

Single-channel DAPI images were segmented with the CPSAM model in Cellpose 4.x using GPU inference and adaptively estimated diameter. Images were normalized and processed in tiles using batch size 32, flow threshold 0.4, cell-probability threshold 0, and tile overlap 0.1. The output was a uint32 label mask, from which centroids and pixel areas were calculated.

##### QuPath

DAPI nuclei were detected with WatershedCellDetection in QuPath using a requested pixel size of 0.5 µm, background radius of 8 µm, Gaussian sigma of 1.5 µm, and accepted nuclear area of 10–400 µm^2^. Watershed post-processing and boundary smoothing were enabled. The DAPI threshold was 100 for Xenium and 30 for CosMx. Detected nuclei were expanded by 0, 5, or 10 µm to define QuPath 0, QuPath 5, and QuPath 10. Exported GeoJSON polygons were rasterized into label masks.

##### BIDCell

BIDCell jointly used DAPI, transcript coordinates, a platform-matched expression reference, and positive and negative markers. Target resolution was 1.0 µm/pixel and patch size was 64. The custom model assigned weight 1 to nuclear exclusion, over-segmentation, cell connectivity, overlap, positive-marker, and negative-marker losses. Xenium v1 was trained for one epoch and 1,500 steps; Xenium 5K and CosMx were trained for one epoch and 1,000 steps. Connected output labels were converted into a common mask, centroid, and area representation.

##### Transcript re-quantification

For non-vendor segmentations, each filtered transcript was mapped to the mask pixel at floor (x/pixel size) and floor (y/pixel size). Transcripts outside the image or on label 0 were left unassigned; all other transcripts were aggregated by cell label and gene into a sparse cell-by-gene count matrix. Pixel centroids were converted to micrometers using the mask pixel size, yielding a consistent h5ad representation for Cellpose, BIDCell, and the three QuPath settings.

#### 2.3 Transcript-purification methods

##### CellAdmix

CellAdmix received transcript coordinates, genes, segmented-cell assignments, and initial cell groups. Initial groups were derived from the unpurified matrix after retaining cells with at least five counts, total-count normalization, log1p transformation, PCA with up to 30 components, construction of a 10-nearest-neighbor graph, and Leiden clustering at resolution 0.5. The model used 10 NMF factors, up to 2,000 cells per class, up to 10,000 molecular-neighborhood records, and three initializations. The conditional random field used 10 neighbors, a same-label ratio of 5, and at most 40 iterations. Graph construction sampled at most 400 transcripts for dense cells, and factor learning used at most 300 transcripts per cell. A bridge test with 20 neighbors and 200 cells per class identified incompatible factor–cell-type combinations at P < 0.1; transcripts assigned to these combinations were removed and counts were re-aggregated.

##### Split

The segmentation matrix, cell coordinates, and identifiers were exported to R. Reference profiles were generated as subclass-level trimmed means from the Allen mouse-brain reference, using 30 Poisson-resampled pseudo-cells per subclass. RCTD was run with UMI_min = 10, counts_MIN = 5, UMI_min_sigma = 20, CELL_MIN_INSTANCE = 5, and doublet mode. Split rctd_based_purify was then applied with singlet purification, and the resulting gene-by-cell counts were converted back to cell-by-gene H5AD format.

##### Ovrlpy

Ovrlpy assessed transcript overlap using vertical signal integrity (VSI). Transcripts were assigned through the vendor cell_id or a segmentation mask. For CosMx, z indices were converted using 0.8 µm per layer and records with *z* < 0 were removed. Analysis used eight workers. If multiple VSI values were returned for one cell, their mean was used. Within each data variant, cells below the 10th percentile of VSI were removed; retained cells preserved their original expression matrix.

#### 2.4 Simulated perturbations

##### Segmentation error

Under-segmentation and over-segmentation were simulated together. A nearest-neighbor index was built from cell centroids, and approximately 10% of cells were involved in pairwise merging, corresponding to *n* × 0.10/2 = 0.05*n* nearest-neighbor cell pairs; counts were summed and coordinates were averaged. Next, 10% of surviving cells were split. For each gene, counts were distributed between two daughter cells as *Binomial* (*n*, 0.5), and daughter coordinates received Gaussian displacement with a standard deviation of three coordinate units.

##### Transcript-assignment noise

Transcript noise combined misassignment, dropout, and ambient signal. For each cell and gene, floor (10% of the count) was transferred to the spatial nearest neighbor. Remaining counts underwent binomial thinning with retention probability 0.9. Ambient counts were sampled as *Poisson*(*genemean* × 0.05) and added only to cell–gene positions that were already non-zero. PYTHONHASHSEED was fixed to ensure stable name-derived random seeds across processes.

#### 2.5 Evaluation of image-preprocessing quality

Upstream image-processing workflows were evaluated independently of the count-matrix preprocessing benchmark using three biologically motivated metrics: marker-purity F1 score, negative-marker fraction, and neighborhood contamination. Metrics were calculated across all cells within the predefined spatial region rather than within a selected cell type. Nine broad brain cell classes were considered: excitatory neuron, inhibitory neuron, astrocyte, oligodendrocyte, oligodendrocyte precursor cell, microglia/macrophage, endothelial cell, pericyte/vascular smooth muscle cell, and ependymal cell. Only marker genes represented in the corresponding platform panel were used.

For cell i and class t, the class score was defined as the mean log-transformed count across the available positive markers *M*_*t*_:

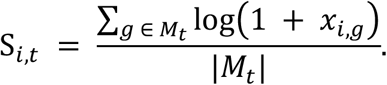

The highest-scoring class was assigned to each cell, and cells with zero counts across all available markers were left unassigned. Markers of the assigned class were treated as positive markers, whereas markers from the other eight classes formed the negative-marker set. Purity precision, positive-marker recall, negative-marker fraction, and marker-purity F1 score were calculated as follows:

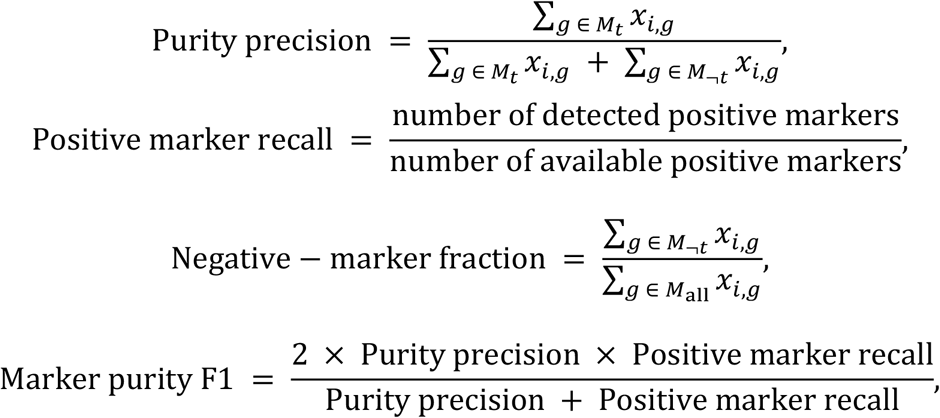

where *M*_*t*_ denotes the positive-marker set for the assigned cell type *t*, and *M*_¬ *t*_ = (⋃_*k*≠ *t*_ *M*_*k*_) \*M*_*t*_ denotes markers associated with other cell types.

Marker purity F1 measures whether transcripts assigned to a cell are consistent with its expected marker profile, whereas the negative marker fraction quantifies biologically incompatible marker expression and therefore reflects potential transcript misassignment.

For neighborhood contamination, cells with finite coordinates and assigned marker-defined classes were retained. A KD-tree returned 13 nearest entries per cell, including the query cell itself, thereby providing at most 12 external neighbors. The median nearest-neighbor distance, *d*_NN_, was used to normalize platform-specific spatial scales. For each directed heterotypic pair from source cell *i* to neighboring cell *j*, positive markers of the neighbor class were treated as neighborhood-negative markers for the source cell. Contamination was defined as:

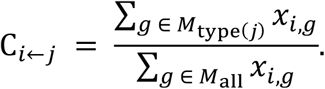

Directed pairs were stratified according to *d*/*d*_NN_ into [0, 1), [1, 2), [2, 4), and [4, ∞). The number of heterotypic pairs and the mean and median contamination were reported for each stratum. Elevated contamination at the shortest distances indicates stronger cross-boundary signal leakage or transcript misassignment. For comparative visualization and construction of the image-processing quality summary, metrics for which lower values indicate better performance (negative-marker fraction and neighborhood contamination) were direction-aligned before aggregation, so that larger values consistently indicated better quality.

For comparative visualization and construction of the image-processing quality summary, metrics were direction-aligned so that larger values consistently indicated better performance. Specifically, the direction-aligned negative-marker and neighborhood-contamination scores were defined as

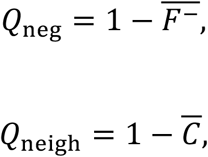

where 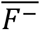 and 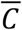 denote the corresponding aggregated negative-marker fraction and neighborhood-contamination values. Marker-based expression purity was summarized as

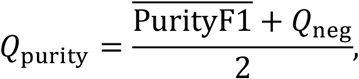

and the overall image-processing quality score was calculated as

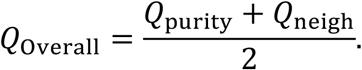

Thus, the two marker-based purity metrics jointly contributed one half of the overall score, whereas neighborhood contamination contributed the other half.

#### 2.6 Robustness of count-matrix preprocessing rankings

For each image-processing variant, all 371 count-matrix preprocessing workflows were independently applied and evaluated using the neighborhood consistency (NC), simulation fidelity (SF), global structure recovery (GSR), and overall-ranking framework described in Section 3. The performance of the 371 workflows under each upstream condition was represented as a preprocessing-ranking profile. Pairwise dissimilarity between profiles was calculated as one minus the Spearman rank correlation. Principal-coordinate analysis was used to visualize the resulting dissimilarities.

The unique contributions of platform, segmentation method, and post-segmentation state to variation among ranking profiles were quantified using a multivariable distance-based model. Partial R^2^ for each factor was calculated by comparing the full model with a reduced model excluding that factor while retaining the other predictors. This analysis distinguished changes in the relative preference among count-matrix preprocessing workflows from changes in the biological quality of the upstream image-processing output.

### 3. Benchmark Metrics

We evaluated the quality of the preprocessed data using three metrics across two aspects: Local and Global. The local aspect included two metrics, NC and SF, used to assess the quality of the *k*-NN graph. The global aspect evaluated recovery ability by leveraging scRNA-seq data.

#### *k*-NN graph identification

To infer the *k*-NN graph from the preprocessed data, we first performed principal component analysis (PCA) for dimensionality reduction. We then calculated the Euclidean distance between cells and identified the 50 nearest neighbors for each cell. This results in a 0/1 binary adjacency matrix, where a value of 1 at entry (*i, j*) indicates that cell *j* is among the 50 neighbors of cell *i*. This entire process was implemented using the Scanpy Python package^60^, with the choice of these parameters following the settings established in the Ref^2^.

#### *k*-NN overlap

The similarity between a pair of *k*-NN graphs, represented by two adjacency matrices *N*_1_ and *N*_2_ ∈ ℕ^*n*×*n*^ (*n* is the number of cells), is measured using the *k*-NN overlap. This overlap is calculated as the sum of element-wise products of the two matrices, normalized by the total number of cells:

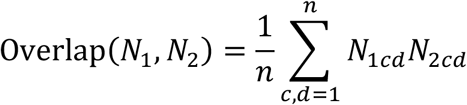

For a *k* = 50 graph, the resulting overlap metric ranges from 0 (no similarity) to 50 (perfect identity between the two graphs).

##### Neighborhood consistency (NC)

NC was adapted from the consistency benchmark introduced by Ahlmann-Eltze and Huber^2^ and was used to quantify the robustness of the inferred cell-state representation to perturbation of the input feature set. For each evaluation, genes were randomly permuted and divided into two non-overlapping subsets. The same count-matrix preprocessing workflow was independently applied to the two subsets, after which PCA and *k*-NN graphs, *N*_1_ and *N*_2_, were constructed separately. NC was then calculated as

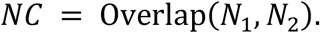

The complete random gene-splitting procedure was repeated five times, and the mean *k*-NN overlap across the five repetitions was used as the final NC score.

NC should be interpreted as a measure of representation robustness rather than direct biological accuracy. Different subsets of genes may contain distinct and non-redundant biological information, particularly in targeted ST panels; therefore, disagreement between two neighborhood graphs does not necessarily imply that either representation is biologically incorrect. Conversely, high NC alone does not establish biological correctness. NC was therefore used as a relative robustness criterion and was complemented by SF and GSR, for which explicitly defined reference structures are available.

##### Simulation fidelity (SF)

SF evaluates whether a preprocessing workflow can recover known continuous cell-state relationships from noisy count observations. The Random Walk and Linear Walk simulations were adapted from the transformation benchmarking framework of Ahlmann-Eltze and Huber^2^. We selected these two models because they represent complementary forms of continuous latent organization: Random Walk generates irregular and stochastic branching trajectories, whereas Linear Walk generates smoother, approximately linear trajectories. Evaluating both simulations reduces dependence of SF on a single assumed latent geometry.

To adapt these simulations to individual ST datasets, the simulation parameters were estimated directly from the corresponding real raw-count matrix. Let

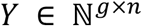

denote the reference count matrix, where *g* is the number of genes and *n* is the number of cells/spots. For cell *c*, the library-size factor was calculated as

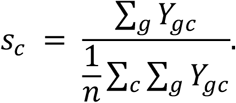

The count matrix was then library-size normalized and log transformed:

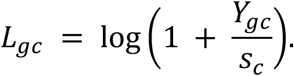

For each gene g, its expression variance was estimated as

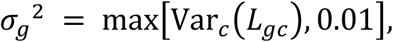

and the negative-binomial overdispersion was estimated by the method of moments:

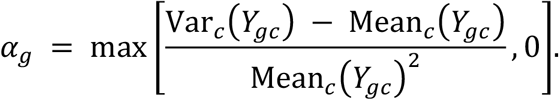

Two latent expression structures were then generated. In the Random Walk simulation, each new cellular state was generated by adding a Gaussian expression offset to its parent state, with periodic branching initiated from randomly selected earlier states:

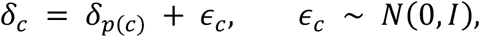

where *p*(*c*) denotes the parent of cell *c*. This procedure generates an irregular but locally continuous branching trajectory.

In the Linear Walk simulation, cellular states followed piecewise linear trajectories between simulated start and end states. For relative position t along a branch,

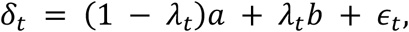

where *a* and *b* denote the start and end states, respectively, *λ*_*t*_ ∈ [0,1] specifies the relative position along the trajectory, and a small Gaussian perturbation was added to introduce local variation. This produces a smoother latent structure than the Random Walk simulation.

The resulting latent offsets were centered and rescaled gene by gene so that their variances matched the corresponding values estimated from the real ST dataset:

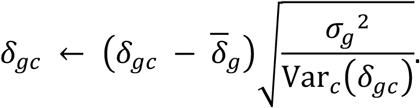

These rescaled latent expression states were treated as the ground-truth cell-state structure for SF.

To generate noisy UMI observations, the baseline log-expression level of each gene was defined as

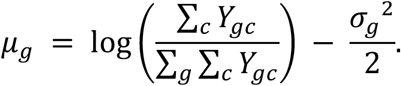

Using the original library size

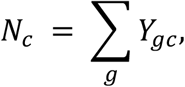

the expected simulated expression was

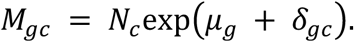

Synthetic UMI counts *Y*_*gc*_ ^sim^ were then sampled from a negative-binomial distribution:

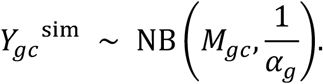

Thus, the simulated count matrix represents noisy count observations generated from an explicitly known continuous latent cell-state structure.

Each count-matrix preprocessing workflow was independently applied to the same simulated count matrix. PCA and k-NN graphs were then constructed separately from the ground-truth latent expression matrix and from the preprocessed simulated matrix. For simulation model m, SF was calculated as

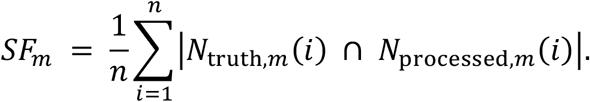

Higher SF indicates more faithful recovery of the known local cell-state relationships.

The final SF score used in the benchmark was calculated as the average of the Random Walk and Linear Walk scores:

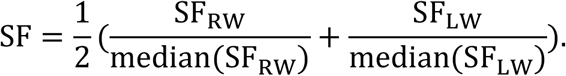

Using both stochastic branching and smooth continuous simulations therefore provides a more robust assessment of preprocessing fidelity than relying on a single latent geometry. The simulation framework and rationale correspond to the description provided in the revised analysis.

##### Global structure recovery (GSR)

GSR evaluates whether a preprocessing workflow preserves sufficient global biological structure to recover known cell-type organization from an iST-like count matrix. The recovery strategy is conceptually related to that used in the Xenium benchmarking study by Marco Salas et al.^6^, in which annotated scRNA-seq datasets were transformed toward Xenium-like measurement characteristics and preprocessing quality was evaluated by recovery of the original cell-type structure. We extended this principle to the different iST datasets included in our benchmark.

For each iST reference dataset, we first obtained a scRNA-seq dataset from CELLxGENE Census with matching species and tissue context. Where donor information was available, cells from a single donor were retained to minimize donor-associated variation, and cell types represented by fewer than 11 cells were excluded. The original scRNA-seq cell-type annotations were preserved as the external ground truth for GSR.

To construct a targeted panel, marker genes were identified using scanpy.tl.rank_genes_groups. For marker ranking, a temporary copy of the scRNA-seq dataset was total-count normalized, log-transformed, and used to identify up to 50 highly ranked genes for each retained cell type. Marker genes were then selected across cell types in a balanced rank-wise manner to prevent highly represented cell types from dominating the simulated panel. If the selected markers did not fill the required panel, additional genes were randomly sampled from the remaining genes until the number of genes matched the corresponding iST reference panel size.

We next modified the scRNA-seq count matrix to reproduce major count-matrix characteristics of the corresponding iST dataset. Mis-segmentation/transcript-assignment contamination was simulated by transferring a randomly sampled fraction of transcript counts from source cells to target cells. Additional low-level count noise was introduced through random count perturbations. When the scRNA-seq dataset contained more cells than the corresponding iST dataset, cells were subsampled while approximately preserving the original cell-type proportions, thereby matching the target cell number where possible.

Sequencing depth was subsequently matched using binomial thinning rather than directly rescaling normalized values. Let *D*_*SC*_ denote the median transcript count per retained simulated cell and *D*_*ST*_ the median transcript count per cell in the corresponding iST reference. The thinning probability was

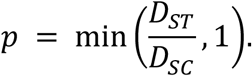

Each count was independently sampled as

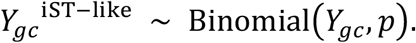

The resulting matrix therefore preserved the known cell-type identities of the source scRNA-seq dataset while approximating several major measurement characteristics of the corresponding iST dataset, including gene-panel size, cell number, transcript depth, and transcript-assignment-associated technical noise. Importantly, this procedure generated an iST-like expression matrix rather than a physical spatial simulation; spatial coordinates, tissue morphology, and spatial graphs were not simulated.

Each count-matrix preprocessing workflow was then applied independently to the same iST-like count matrix. PCA was performed on the resulting representation, followed by construction of a k-NN graph and Leiden clustering. To obtain a comparable clustering granularity across preprocessing workflows, the Leiden resolution was iteratively adjusted until the inferred number of clusters was within ±2 of the number of ground-truth cell types. The known number of cell types was used only to select clustering granularity; the identities of the ground-truth labels were not used during clustering.

The inferred Leiden clusters were compared with the preserved source cell-type annotations using the adjusted Rand index (ARI):

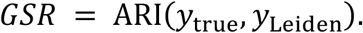

Higher GSR indicates better recovery of the known global cell-type organization. Thus, unlike NC, which evaluates robustness to input-feature perturbation, GSR evaluates recovery against an externally defined biological partition.

#### The overall score

We compute an overall ranking score to comprehensively assess the global performance of each preprocessing. This is achieved by first ranking all preprocessings based on their performance across each individual metric. These individual metric rankings are then averaged to produce a composite score. The overall score is calculated as follows:

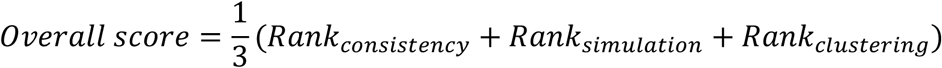

### 4. Influence of hyperparameter settings and refinement operations

We analyzed the influence of each hyperparameter setting and refinement operation through two aspects: *k*-NN graph overlaps and the overall ranking score.

#### Quantifying Influence by *k*-NN graph overlap

As previously stated, for any two preprocessing strategies, *P*_*a*_ and *P*_*b*_, and their corresponding *k*-NN graphs, *N*_*a*_ and *N*_*b*_, the *k*-NN overlap *O*(*P*_*a*_, *P*_*b*_) can be calculated as:

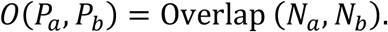

Let *E* be a specific hyperparameter setting or refinement operation set (e.g., Z-score, HVG, overdispersion, or size factors). Let *S* be a set of preprocessing strategies where all choices are identical except for the specific parameter *e ϵ E*. The minimum *k*-NN overlap, *I*_*k*NN_ (*E, S*), can be calculated as:

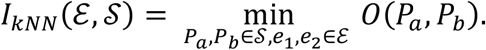

#### Quantifying Influence by overall ranking score

For any two preprocessing strategies, *P*_*a*_ and *P*_*b*_, their overall ranking scores are denoted as *Score*(*P*_*a*_) and *Score*(*P*_*b*_) . Let *E* be a specific hyperparameter setting or refinement operation set. Let *S* be a set of preprocessing strategies where all choices are identical except for the specific parameter *e ϵ E*. The maximum change in the overall score, *I*_*score*_ (*E, S*), can be calculated as:

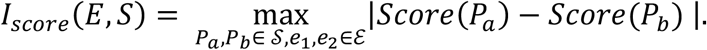

#### PCC

To assess the linear relationship between two variables, *X* and *Y* ∈ ℝ^1×*n*^, we used the PCC, denoted by *r*. The PCC quantifies the strength and direction of the linear association between two continuous variables and is defined as:

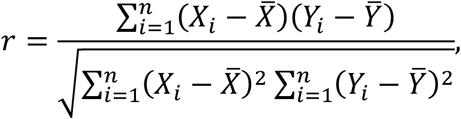

where 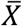 and 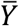 are the means of *X* and *Y*, respectively. We employed the ‘scipy.stats.pearsonr’ function^41^ to calculate the PCC (*r*).

### 5. Influence of data structures

#### Definition of data structure

We considered three data structure features: cell number, panel size, and GELPU. Let *Y* ∈ ℕ^*g*×*c*^ denote the input raw-count matrix obtained from the publicly released dataset, where *c* represents the number of cells, *g* denotes the number of genes, and *Y*_*ij*_ is the count of gene *i* in cell *j*. GELPU was calculated directly from this publicly provided raw-count matrix before any additional filtering of low-expression genes or low-quality cells/spots in our preprocessing pipeline. The definitions of GELPU:

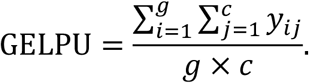

#### Different data structure simulation

To investigate the influence of individual data structure features (cell number, panel size, and GELPU), we generated 17 simulated datasets from the original three mouse brain datasets (10x Visium, Xenium v1, and Xenium 5K). The simulation strategies employed were random cell sampling, random gene selection, matched gene selection, and downsampling.

Random cell sampling. For a given data *X* and a target data *Y*, we performed random cell sampling from *X* to generate a simulated data. This sampling was done such that the resulting data’s cell number matched that of data *Y*.

Random gene selection. For a given data *X* and a target data *Y*, we performed random gene selection from *X* to generate a simulated data. This selection was done such that the resulting data’s panel size (gene number) matched that of data *Y*.

Matched gene selection. For a given data *X* and a target data *Y*, we performed random gene selection from *X* to generate a simulated data. This selection involved choosing genes from *X* that overlapped with *Y* gene panel such that the resulting data’s panel matched that of data *Y*.

Downsampling. For a given data *X* and a target data *Y*, we performed downsampling on the data *X* to generate a simulated data. This process was executed to proportionally reduce the total transcript counts such that the resulting data’s GELPU matched that of data *Y*. This function is implemented in the scuttle R package^42^.

### 6. Variance partitioning of workflow-ranking profiles

#### Workflow-ranking response matrix

Variance partitioning was performed on the complete 45-dataset benchmark and the 371 canonical count-matrix preprocessing workflows. For each dataset, NC, the Random Walk branch of SF, the Linear Walk branch of SF, and, for imaging-based spatial transcriptomics (iST), GSR were divided by their corresponding within-dataset medians. The two normalized SF branches were combined as

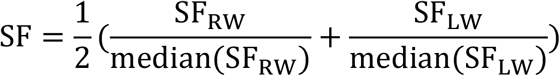

NC and SF were ranked separately for sequencing-based spatial transcriptomics (sST), whereas NC, SF, and GSR were ranked separately for iST. Component ranks used minimum-rank tie handling with ascending numerical ranks; because larger stored quality scores receive larger component ranks, larger component-rank sums indicate better performance. The sum of the applicable component ranks was ranked again with the same tie rule to form a 371-workflow profile for each dataset.

#### Dataset dissimilarity and ordination

Pairwise dissimilarity between datasets was defined as one minus the Spearman correlation between their 371-workflow rank profiles. Because this distance matrix was non-Euclidean, principal-coordinate analysis (PCoA) was performed after a Lingoes correction. The initial Gower matrix was eigendecomposed; when its minimum eigenvalue was negative, twice its absolute value was added to all off-diagonal squared distances, the diagonal was reset to zero, and square roots were taken before the corrected Gower matrix was eigendecomposed. Only positive PCoA coordinates were retained as the multivariate response.

#### Explanatory variables and metadata processing

The predictors were ST platform, grouped tissue, grouped species, gene panel size, GELPU, and cell number. Tissue was grouped as Brain, Kidney, mouse olfactory bulb (MOB), or Other; species was grouped as Human, Mouse, or Other. GELPU was calculated from the raw count matrix as

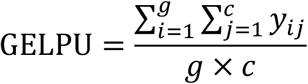

where *Y*_*ij*_ is the count of gene *i* in cell/spot *j*. GELPU and cell number were log10-transformed and standardized using the sample standard deviation (ddof=1). Data 12 stored log1p values in X and was inverted using expm1 before count summation. Only CodeClass = Endogenous features were retained for Data 45 in this analysis, yielding 950 genes. Tissue and species for Data 42–45 were manually curated because these fields were absent from data/metadata.tsv.

#### Distance-based models

Categorical predictors were encoded using treatment-style dummy variables with the first level dropped. Ordinary R^2^ values were obtained from six separate single-predictor multivariate regressions and therefore represent marginal associations that include shared variation. Conditional effects were estimated in a full model containing all six predictors. For each term, partial R^2^ was defined as

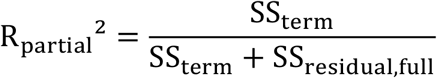

where SSterm is the additional sum of squares obtained by adding the term to the corresponding reduced model and SSresidual,full is the residual sum of squares from the full model. Raw/semi-partial and adjusted estimates were retained together with conditional partial R^2^.

### 7. Ranking robustness and sensitivity analyses

#### Parameter sensitivity

The default evaluation graph was constructed using (*k*=50), 20 principal components, and Euclidean distance. To assess sensitivity to graph construction, one parameter was varied at a time while the other two were kept at their default values. The tested (*k*) values were 5, 10, 20, 30, 50, 75, 100, 150, and 200; the numbers of principal components were 5, 10, 20, 30, 50, 75, 100, 150, and 200; and the distance metrics were Euclidean, cosine, and correlation. This resulted in 19 unique parameter settings. For each setting, metric branches were normalized by their within-dataset medians, the two SF branches were averaged, and workflows were ranked using NC and SF for sST or NC, SF, and GSR for iST.

#### Data dropout

For each data, 10% of cells/spots or 10% of genes were randomly removed using seed 42, and workflow rankings were compared with those obtained from the complete dataset using the default graph parameters. A single random subsample was generated for each dropout condition. Workflows were ordered according to their default combined ranks.

#### One-hundred-repeat NC stability

For each repeat, the gene set was randomly permuted and divided equally into two subsets. Within each subset, genes were centered, 20 principal components were obtained using randomized SVD, and self-excluded FAISS (*k)*-nearest-neighbor graphs were constructed with (*k*=50). NC was calculated as the mean number of shared neighbors between the two graphs across all cells or spots.

### 8. Downstream task evaluation

To assess the biological utility of the different preprocessing strategies, we evaluated their performance on four critical downstream tasks: cell type identification, spatial domain identification, marker-recovery validation, SVG-ranking stability validation.

#### 8.1 Cell type identification

##### Clustering

Consistent with the *k*-NN graph settings established previously, we first performed Principal Component Analysis (sc.pp.pca) to reduce the dimensionality to 20 components for each preprocessed data, followed by constructing a *k*-NN graph (sc.pp.neighbors) using 50 neighbors. Finally, we applied the leiden (sc.tl.leiden) algorithm with its default resolution to perform clustering.

##### Five-fold cross-validation

For each preprocessed dataset, we trained a Support Vector Machine (SVM) classifier on each preprocessed data using the ground truth cell type labels. The dataset was partitioned into five subsets; in each iteration, the model was trained on four subsets and evaluated on the held-out subset. Macro-averaged precision, macro-averaged recall, and global accuracy were calculated independently for each held-out fold, and the final reported value for each metric was the mean across the five folds.

##### Metrics

For multiclass cell type classification with C classes, precision and recall were first calculated separately for each class using a one-versus-rest scheme. For class c, all observations belonging to class c were treated as positive and observations belonging to all other classes as negative. Class-specific precision and recall were defined as

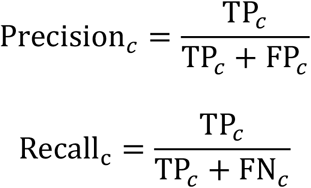

where TP_*c*_, FP_*c*_, and FN_*c*_ denote the numbers of true positives, false positives, and false negatives, respectively, for class *c*. Macro-averaged precision and recall were then calculated as the unweighted means across all *C* classes:

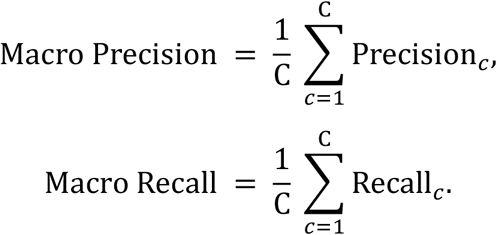

Accuracy was calculated globally rather than averaged across classes and was defined as the proportion of correctly classified observations among all observations:

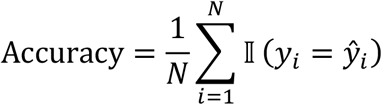

where is *N* the total number of observations, *y*_*i*_ is the ground-truth label, *ŷ*_*i*_ is the predicted label, and I(·) is the indicator function.

##### Statistical significance testing

We conducted pairwise statistical tests on the resulting scores to assess the statistical significance of performance differences between key preprocessing configurations. Given the unknown underlying score distributions, we employed the non-parametric two-sided Mann-Whitney U test. The Mann-Whitney U test (also known as the Wilcoxon Rank-Sum test) is a non-parametric statistical test used to determine whether there is a significant difference in the location of the distribution (median) between two independent samples. Since it does not rely on assumptions about the data’s distribution (such as normality), it is highly robust. The resulting statistical significance was then visualized directly on the plots using the star format (e.g., * for p < 0.05), clearly highlighting the comparisons that exhibited a statistically significant difference in performance.

#### 8.2 Spatial domain identification

##### Metrics

We compared the predicted spatial cluster labels against the ground truth spatial labels using three information-theoretic metrics: HOM, NMI, COM. These metrics are independent of the absolute values of the labels, only considering the agreement between cluster assignments. All scores range from 0 to 1, where 1 indicates a perfect match.

HOM measures whether each cluster contains only members belonging to a single class (i.e., pure clusters)^43^. A score of 1.0 is achieved if all clusters contain data points from only one true class. It is calculated as:

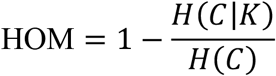

where *H*(*C*) is the entropy of the true class labels *C. H*(*C*|*K*) is the conditional entropy of the true class labels given the cluster assignments (*K*).

COM measures whether all data points belonging to a single true class are assigned to the same cluster^43^. A score of 1.0 is achieved if all members of a given true class are grouped into the same cluster. It is calculated as:

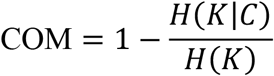

where *H*(*K*) is the entropy of the cluster assignments (*K*). *H*(*K*|*C*) is the conditional entropy of the cluster assignments given the true class labels (*C*).

NMI quantifies the agreement between the ground-truth class labels and inferred cluster assignments based on their mutual information. Let *U* denote the ground-truth labels and *V* denote the inferred cluster labels. NMI was calculated using arithmetic-mean normalization as:

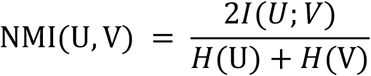

where *I*(U; V) is the mutual information between U and V, and *H*(U) and *H*(V) denote their respective entropies. NMI ranges from 0 to 1, with larger values indicating greater agreement between the inferred clusters and ground-truth labels.

##### Spatial domain identification methods

We evaluated the different preprocessings data by testing the performance of three advanced spatial clustering algorithms: SEDR, GraphST, and SpaGCN.

##### SEDR

SEDR integrates gene-expression and spatial information by combining a deep autoencoder with a graph-based latent representation. The resulting embedding was used for spatial domain clustering. In the parameter-sensitivity analysis, the number of spatial neighbors was set to 6 or 12, and the autoencoder/graph-convolution hidden dimensions were set to 64–16 or 128–32. The default setting used 12 spatial neighbors and hidden dimensions of 128–32.

##### GraphST

GraphST is highly effective for spatial domain identification because it learns informative and discriminative spot representations by enforcing local spatial consistency^31^. By using a contrastive learning mechanism within a GNN framework, GraphST minimizes the embedding distance between spatially adjacent spots. This process allows the algorithm to robustly delineate fine-grained tissue structures and accurately identify spatial domains, even across multiple tissue slices in a joint (multi-sample) analysis. We evaluated models trained for 100 or 300 epochs, with 300 epochs used as the default setting.

##### SpaGCN

SpaGCN is specifically designed to identify spatial domains characterized by coherent gene expression and histological patterns^30^. It achieves this by constructing a spatial graph and employing GCNs to aggregate gene expression from neighboring spots. This localized aggregation suppresses noise and highlights expression differences across regions, enabling the accurate and computationally efficient detection of spatial domains. Subsequently, these identified domains are used to guide differential expression analysis, revealing genes that define the detected spatial organization. The parameter (p), which controls the contribution of histological information to the spatial graph, was evaluated at 0.2, 0.5, and 0.7, with (p=0.5) used as the default setting.

We input the preprocessed data into each of the three algorithm frameworks. For each run, we supplied the ground truth number of spatial domains and executed the algorithms according to their default parameters.

#### 8.3 Marker-recovery validation

Marker-recovery analysis was used to assess whether preprocessing preserved multigene expression patterns informative of genes excluded from the input features. For each data, top-20 highly variable genes were withheld from the feature matrix and treated as prediction targets. Expression of each held-out marker was binarized according to the predefined analysis criteria. The remaining genes were reduced to 50 principal components, which were used to predict each marker independently with k-nearest-neighbor, logistic-regression, and multilayer-perceptron classifiers.

Cells or spots were randomly divided into equally sized training and test sets using seed 42. The same split was used for all preprocessing workflows within each dataset to enable paired comparison. Prediction performance was evaluated for each marker using AUROC, average precision, balanced accuracy, and F1 score, and then macro-averaged across the 20 markers.

For each platform, the top 30 and bottom 30 count-matrix preprocessing workflows were selected independently of the marker-recovery results. Workflow ranking was based on NC and SF for sST data and on NC, SF, and GSR for iST data. Workflows containing HVG selection were included. Scores were summarized by dataset, classifier, workflow group, and evaluation metric, and top-minus-bottom differences were calculated within each dataset while retaining platform- and classifier-specific comparisons. The same workflow occurring in both sST and iST analyses was not treated as an independent biological replicate.

#### 8.4 SVG-ranking stability validation

SVG-ranking stability was evaluated by comparing spatial-gene rankings obtained from the full dataset with those obtained after random subsampling. Workflows containing non-spatial HVG selection were excluded to avoid restricting the gene set before SVG detection. This left 187 of the 371 preprocessing workflows for evaluation. Within each platform, the top 30 and bottom 30 eligible workflows were selected according to NC and SF for sST data and NC, SF, and GSR for iST data. For each workflow, SVG analysis was performed on the complete dataset and on random subsamples containing 80% of the cells or spots. SVGs were ranked independently using Moran’s I and SPARK-X, with option=“mixture” used for SPARK-X.

Agreement among the highest-ranked genes was additionally evaluated using Jaccard similarity on the union of the two top-gene sets. Top-gene sets were defined as the highest-ranked 5%, 10%, or 20% of genes, with a minimum of 20 and a maximum of 500 genes. For the full-data set *A* and subsampled set *B*, Jaccard similarity was calculated as

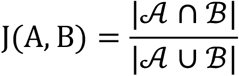

## Data availability

Data 1, Data 2, and Data 4 to Data 8 were obtained from: https://www.10xgenomics.com/data/.

Data 3 was obtained from: https://mender-tutorial.readthedocs.io/en/latest/Visium_MOB.html.

Data 9 to Data 15 were obtained from: https://gene.ai.tencent.com/SpatialOmics/.

Data 16 to Data 19 were obtained from: https://db.cngb.org/stomics/.

Data 20 and Data 21 were obtained from Ref^21^.

Data 22 to Data 24 were obtained from: https://gene.ai.tencent.com/SpatialOmics/.

Data 25 and Data 26 were obtained from Ref^45^.

Data 27 was obtained from Ref^46^. Data 28 was obtained from Ref^25^.

Data 29 was obtained from: https://info.vizgen.com/mouse-liver-data.

Data 30 to Data 32, Data 34, and Data 35 were obtained from: https://www.10xgenomics.com/data/.

Data 33 was obtained from Ref^47^.

Data 36 to Data 41 were obtained from: https://www.10xgenomics.com/data/.

Data 42, Data 43 were obtained from: https://www.10xgenomics.com/data/.

Data 44, Date 45 were obtained from: https://brukerspatialbiology.com/products/cosmx-spatial-molecular-imager/ffpe-dataset/.

scRNA-seq Data were obtained from: https://cellxgene.cziscience.com/data/.

For several very large-scale datasets, benchmarking was performed on a selected spatial region rather than the entire dataset because of computational constraints.

## Computational resources

Intel(R) Xeon(R) Gold 5318Y CPUs (2.10 GHz, 36 MB L3 cache, 48 cores in total), 503 GB of system memory, and an NVIDIA RTX A6000 graphics processing unit (GPU) with 48 GB of GPU memory (CUDA version 12.2).

## Code availability

All code used to reproduce the analyses and generate the figures in this study is publicly available at https://github.com/YSTLab/ST-data-preprocessing-benchmark.git. The repository includes the st-preprocess Python toolkit implementing all 371 count-matrix preprocessing workflows evaluated in this study, together with platform- and data structure-based recommendation functions. Detailed installation instructions, tutorials, example commands, input/output specifications, and both Python and command-line interfaces are provided to facilitate reproducible application of the preprocessing framework.

## Author contributions

Z.Y. and Y.D. conceived and designed the benchmarking strategy. The manuscript was written by Z.Y. and Y.D. The primary data analysis rationale and figure generation were performed by Y.D. and Z.Y. Z.Y. provided the main guidance on figure layout and arrangement. S.L. contributed specifically by offering perspectives and insights for result interpretation, assisting with figure plotting, and completing the drawing of schematic diagrams for most main figures, as well as the splicing and arrangement of some result figures. Z.W. and S.L. contributed to the manuscript revision and polishing. Data collection was jointly conducted by Y.D. and H.S. Q.Z. and D.Z. participated in data collection and provided guidance on figure generation. All authors have read and approved the final version of the manuscript.

### Acknowledgements

The authors acknowledge the support by the Computational Biology Program (25JS2850200 (Z.Y.)) of Science and Technology Commission of Shanghai Municipality (STCSM), National Nature Science Foundation of China (62303119 (Z.Y.), 32470706 (Z.Y.)), Shanghai Science and Technology Development Funds (23YF1403000 (Z.Y.)), Fund of Fudan University and Cao’ejiang Basic Research (24FCA10 (Z.Y.)).

## Inclusion & Ethics

Not relevant.

