## Supplemental Figure for "Platform-specific count-matrix preprocessing workflows for spatial transcriptomics data analysis"

**Figure S1**

| Data ID | Spatial type | Platform | Tissue | Spots/Cells | Panel size | GELPU |
|---|---|---|---|---|---|---|
| Data 1 | Sequencing-based | 10x Visium | MCH | 2,438 | 19,465 | 0.837 |
| Data 2 | | | MKD | 3,055 | 32,285 | 0.917 |
| Data 3 | | | MOB | 918 | 31,053 | 0.339 |
| Data 4 | | | Human prostate | 2,543 | 17,943 | 0.286 |
| Data 5 | | | Human ovarian cancer | 3,455 | 17,943 | 1.542 |
| Data 6 | | | Human intestine cancer | 2,660 | 17,943 | 1.351 |
| Data 7 | | | Human heart | 4,247 | 36,601 | 0.239 |
| Data 8 | | | Human breast cancer | 2,518 | 17,943 | 0.742 |
| Data 9 | | Slide-seq V2 | MCH | 33,611 | 22,683 | 0.018 |
| Data 10 | | | MKD | 29,011 | 20,262 | 0.008 |
| Data 11 | | | MOB | 21,724 | 21,220 | 0.022 |
| Data 12 | | | Human kidney | 18,471 | 25,018 | 0.009 |
| Data 13 | | | Mouse testis | 31,659 | 24,450 | 0.032 |
| Data 14 | | | Human brain | 37,363 | 28,763 | 0.023 |
| Data 15 | | | Mouse brain | 35,895 | 19,758 | 0.005 |
| Data 16 | | Stereo-seq | MCH | 38,476 | 26,177 | 0.136 |
| Data 17 | | | Arabidopsis leaves | 721 | 18,257 | 0.080 |
| Data 18 | | | Mouse Liver | 35,667 | 23,498 | 0.292 |
| Data 19 | | | Ambystoma mexicanum brain | 9,440 | 27,600 | 0.119 |
| Data 20 | | Array-seq | MCH | 57,258 | 32,285 | 0.029 |
| Data 21 | | | MOB | 13,784 | 32,285 | 0.110 |
| Data 22 | Imaging-based | STARmap | MCH | 29,623 | 1,022 | 0.200 |
| Data 23 | | | MOB | 15,791 | 1,022 | 0.143 |
| Data 24 | | seqFISH+ | MKD | 220,753 | 1,298 | 0.154 |
| Data 25 | | MERFISH | MCH | 13,510 | 1,122 | 0.132 |
| Data 26 | | | MOB | 12,429 | 1,122 | 0.346 |
| Data 27 | | | MKD | 212,090 | 307 | 0.482 |
| Data 28 | | | Human embryo brain | 562,306 | 300 | 0.196 |
| Data 29 | | | Mouse liver | 395,215 | 347 | 2.449 |
| Data 30 | | Xenium v1 | MCH | 59,935 | 347 | 0.591 |
| Data 31 | | | Mouse embryo | 1,355,849 | 379 | 0.370 |
| Data 32 | | | Mouse brain | 162,033 | 248 | 1.199 |
| Data 33 | | | Human lung | 12,915 | 343 | 0.482 |
| Data 34 | | | Human colorectal cancer | 307,762 | 422 | 0.178 |
| Data 35 | | | Human brain cancer | 816,769 | 480 | 0.363 |
| Data 36 | | Xenium 5K | Human lung cancer | 278,328 | 5,001 | 0.088 |
| Data 37 | | | Human ovarian cancer | 407,124 | 5,101 | 0.051 |
| Data 38 | | | Human lymph node | 708,983 | 4,624 | 0.058 |
| Data 39 | | | Human skin cancer | 112,551 | 5,006 | 0.093 |
| Data 40 | | | Human prostate cancer | 193,000 | 5,006 | 0.041 |
| Data 41 | | | MCH | 63,173 | 5,006 | 0.274 |
| Data 42 | Sequencing-based | Visium HD | MCH | 393,543 | 19,070 | 0.012 |
| Data 43 | | | MKD | 502,009 | 19,070 | 0.026 |
| Data 44 | Imaging-based | CosMx | Huamn liver | 793,318 | 1,000 | 0.834 |
| Data 45 | | | MCH | 38,996 | 1,147 | 1.518 |

**Figure S2**

| Stage | Category | Operation |
|---|---|---|
| Stage 1: Hyperparameter Settings | Size factors | 10 100 1000 Mean Volume |
| | Overdispersion | 0 0.05 Est. |
| Stage 2: Transformations | Delta method | log(y/s + 1) acosh(2αy/s + 1) Scran log(y/s + 1/(4α)) TMM Deseq2 |
| | GLM residuals | Analytic pearson SCTransform Pearson Random quantile Pearson (no clip) |
| | Lat Expr | Sanity Normaliser |
| Stage 3: Refinement operations | Clip | None 10 sqrt_n sqrt_(30/n) |
| | HVG | Yes No |
| | Z score | Yes No |

Figure S3

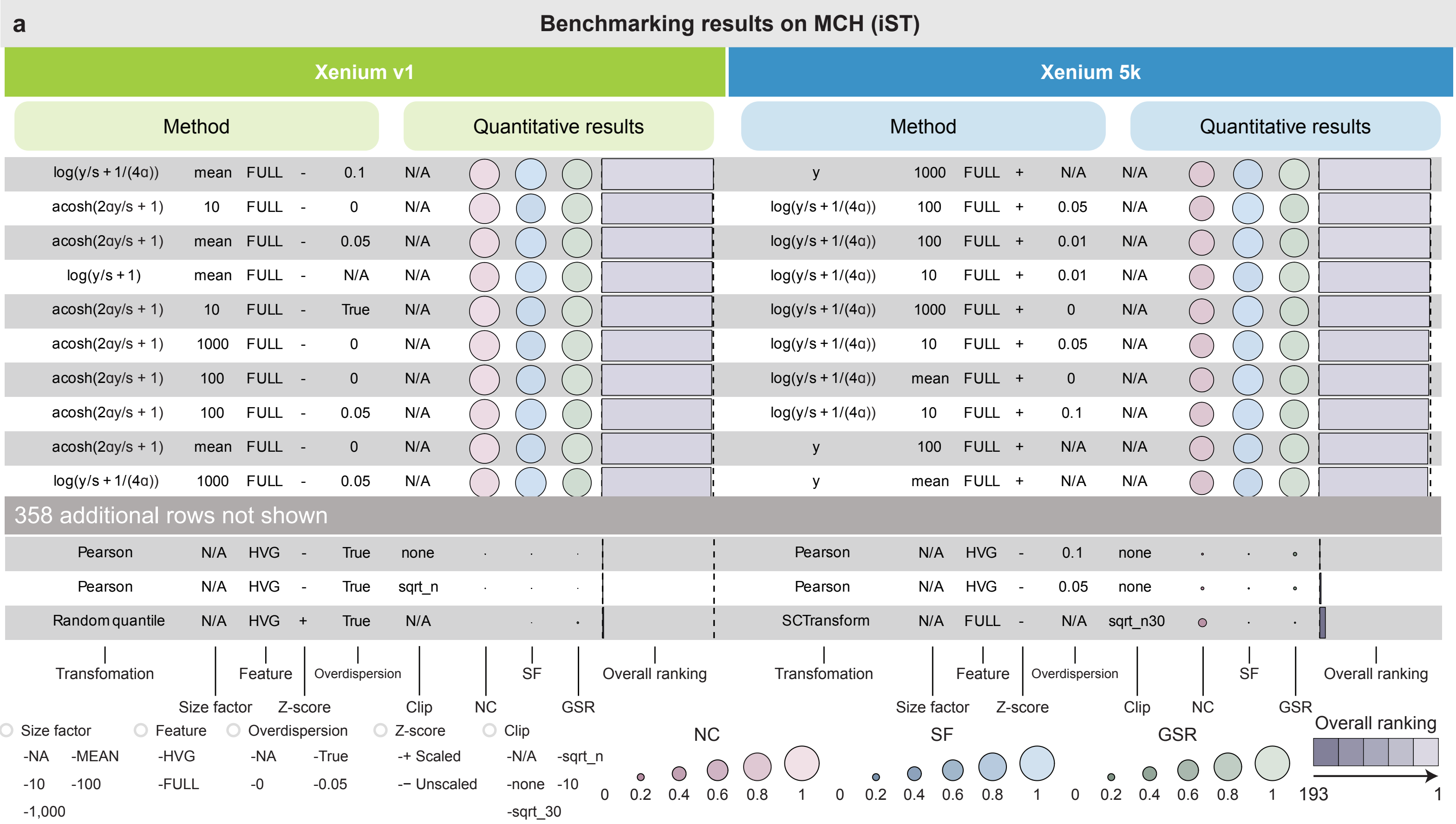
a
Benchmarking results on MCH (iST)
Xenium v1
Method
Quantitative results
log(y/s + 1/(4α)) mean FULL - 0.1 N/A
acosh(2αy/s + 1) 10 FULL - 0 N/A
acosh(2αy/s + 1) mean FULL - 0.05 N/A
log(y/s + 1) mean FULL - N/A N/A
acosh(2αy/s + 1) 10 FULL - True N/A
acosh(2αy/s + 1) 1000 FULL - 0 N/A
acosh(2αy/s + 1) 100 FULL - 0 N/A
acosh(2αy/s + 1) 100 FULL - 0.05 N/A
acosh(2αy/s + 1) mean FULL - 0 N/A
log(y/s + 1/(4α)) 1000 FULL - 0.05 N/A
Pearson N/A HVG - True none
Pearson N/A HVG - True sqrt_n
Random quantile N/A HVG + True N/A
Transfomation
Size factor
Feature
Z-score
Overdispersion
Clip
NC
SF
GSR
Overall ranking
Xenium 5k
Method
Quantitative results
y 1000 FULL + N/A N/A
log(y/s + 1/(4α)) 100 FULL + 0.05 N/A
log(y/s + 1/(4α)) 100 FULL + 0.01 N/A
log(y/s + 1/(4α)) 10 FULL + 0.01 N/A
log(y/s + 1/(4α)) 1000 FULL + 0 N/A
log(y/s + 1/(4α)) 10 FULL + 0.05 N/A
log(y/s + 1/(4α)) mean FULL + 0 N/A
log(y/s + 1/(4α)) 10 FULL + 0.1 N/A
y 100 FULL + N/A N/A
y mean FULL + N/A N/A
Pearson N/A HVG - 0.1 none
Pearson N/A HVG - 0.05 none
SCTransform N/A FULL - N/A sqrt_n30
Transfomation
Size factor
Feature
Z-score
Overdispersion
Clip
NC
SF
GSR
Overall ranking
358 additional rows not shown
Size factor
-NA -MEAN
-10 -100
-1,000
Feature
-HVG
-FULL
Overdispersion
-NA -True
-0 -0.05
Z-score
-+ Scaled
-− Unscaled
Clip
-N/A -sqrt_n
-none -10
-sqrt_30
NC
0 0.2 0.4 0.6 0.8 1
SF
0 0.2 0.4 0.6 0.8 1
GSR
0 0.2 0.4 0.6 0.8 1
Overall ranking
193 1

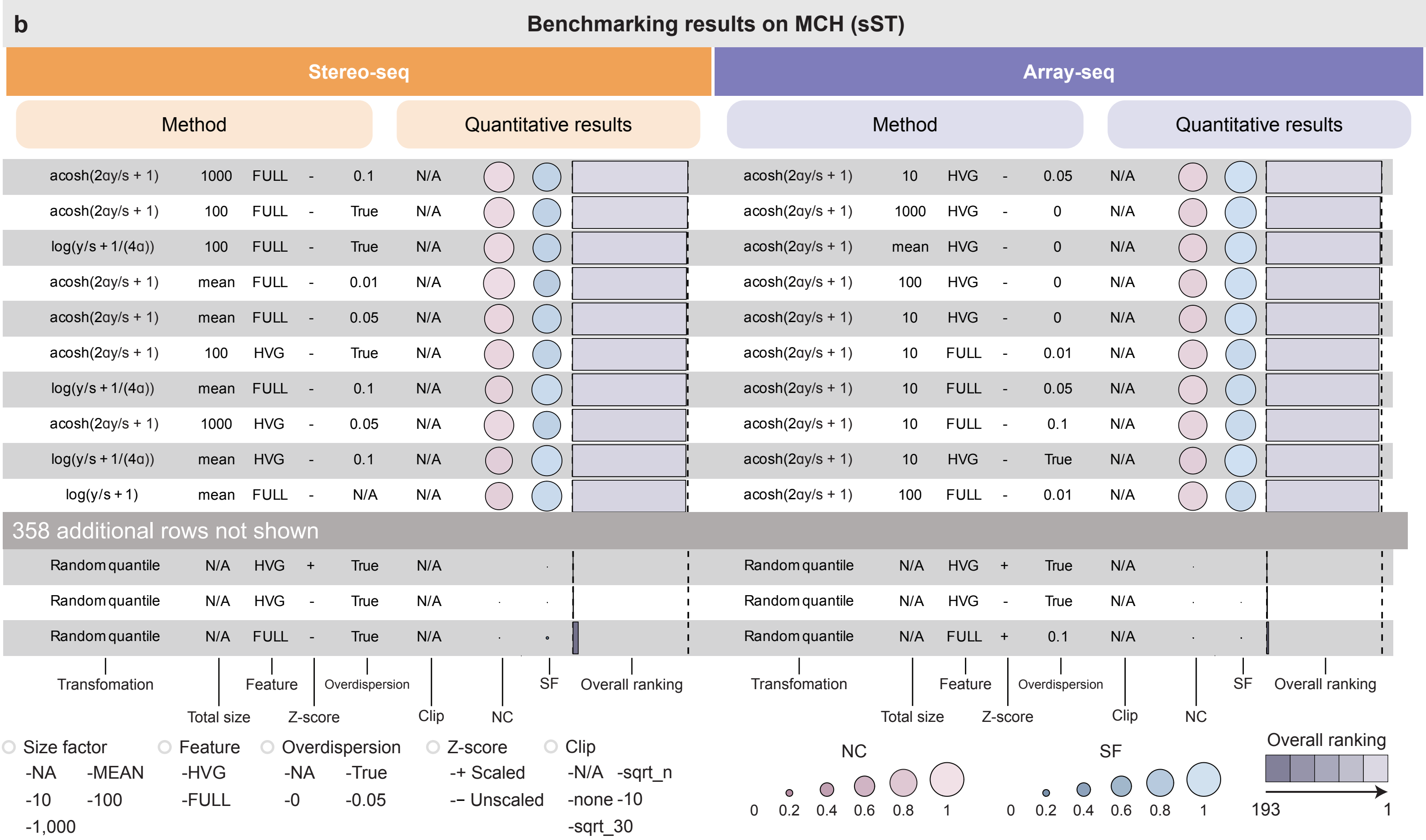
b
Benchmarking results on MCH (sST)
Stereo-seq
Method
Quantitative results
acosh(2αy/s + 1) 1000 FULL - 0.1 N/A
acosh(2αy/s + 1) 100 FULL - True N/A
log(y/s + 1/(4α)) 100 FULL - True N/A
acosh(2αy/s + 1) mean FULL - 0.01 N/A
acosh(2αy/s + 1) mean FULL - 0.05 N/A
acosh(2αy/s + 1) 100 HVG - True N/A
log(y/s + 1/(4α)) mean FULL - 0.1 N/A
acosh(2αy/s + 1) 1000 HVG - 0.05 N/A
log(y/s + 1/(4α)) mean HVG - 0.1 N/A
log(y/s + 1) mean FULL - N/A N/A
Random quantile N/A HVG + True N/A
Random quantile N/A HVG - True N/A
Random quantile N/A FULL - True N/A
Transfomation
Total size
Feature
Z-score
Overdispersion
Clip
NC
SF
Overall ranking
Array-seq
Method
Quantitative results
acosh(2αy/s + 1) 10 HVG - 0.05 N/A
acosh(2αy/s + 1) 1000 HVG - 0 N/A
acosh(2αy/s + 1) mean HVG - 0 N/A
acosh(2αy/s + 1) 100 HVG - 0 N/A
acosh(2αy/s + 1) 10 HVG - 0 N/A
acosh(2αy/s + 1) 10 FULL - 0.01 N/A
acosh(2αy/s + 1) 10 FULL - 0.05 N/A
acosh(2αy/s + 1) 10 FULL - 0.1 N/A
acosh(2αy/s + 1) 10 HVG - True N/A
acosh(2αy/s + 1) 100 FULL - 0.01 N/A
Random quantile N/A HVG + True N/A
Random quantile N/A HVG - True N/A
Random quantile N/A FULL + 0.1 N/A
Transfomation
Total size
Feature
Z-score
Overdispersion
Clip
NC
SF
Overall ranking
358 additional rows not shown
Size factor
-NA -MEAN
-10 -100
-1,000
Feature
-HVG
-FULL
Overdispersion
-NA -True
-0 -0.05
Z-score
-+ Scaled
-− Unscaled
Clip
-N/A -sqrt_n
-none -10
-sqrt_30
NC
0 0.2 0.4 0.6 0.8 1
SF
0 0.2 0.4 0.6 0.8 1
Overall ranking
193 1

### Figure S4

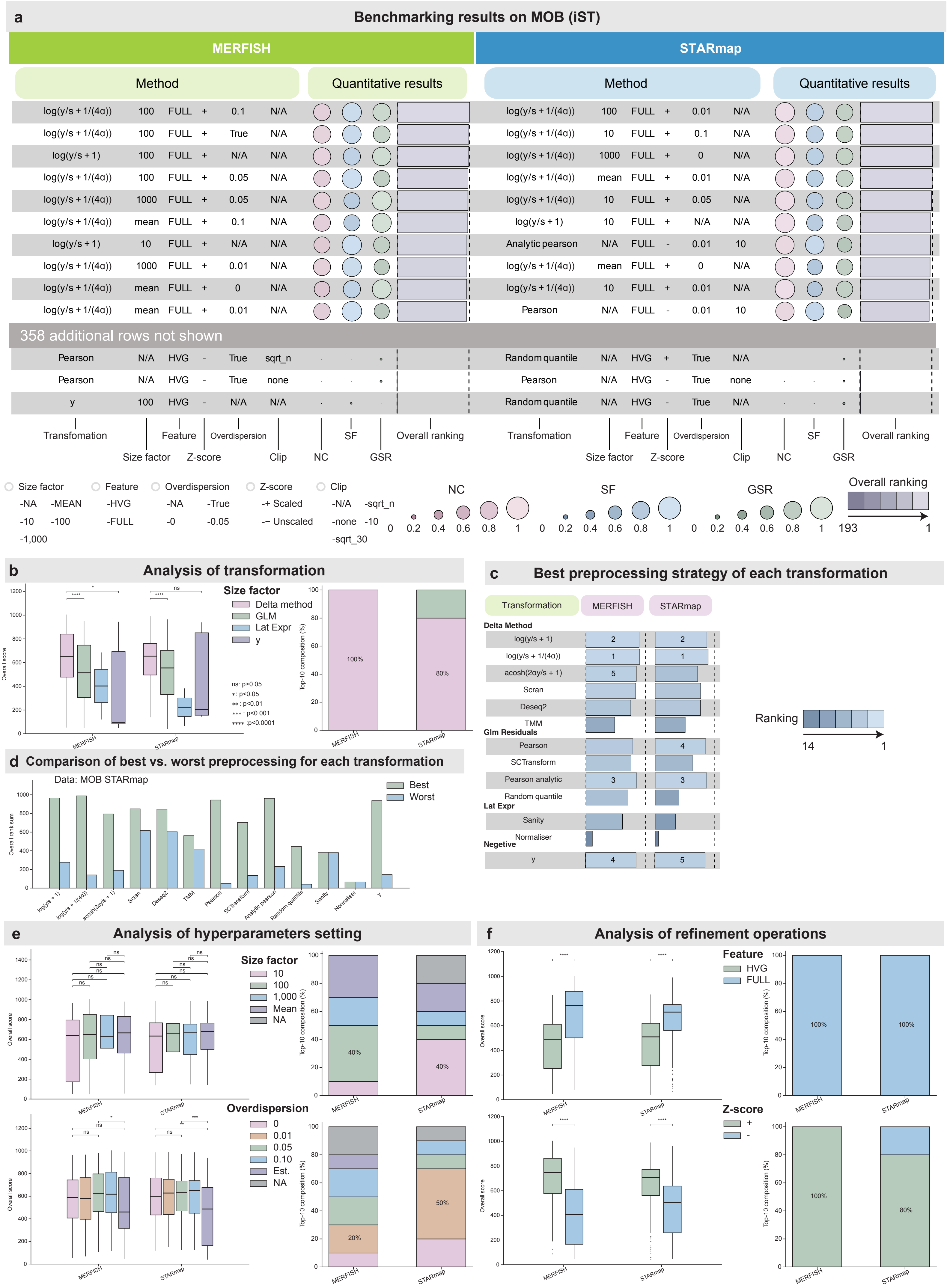

a
Benchmarking results on MOB (iST)
MERFISH
STARmap
Method
Quantitative results
log(y/s + 1/(4α)) 100 FULL + 0.1 N/A
log(y/s + 1/(4α)) 100 FULL + True N/A
log(y/s + 1) 100 FULL + N/A N/A
log(y/s + 1/(4α)) 100 FULL + 0.05 N/A
log(y/s + 1/(4α)) 1000 FULL + 0.05 N/A
log(y/s + 1/(4α)) mean FULL + 0.1 N/A
log(y/s + 1) 10 FULL + N/A N/A
log(y/s + 1/(4α)) 1000 FULL + 0.01 N/A
log(y/s + 1/(4α)) mean FULL + 0 N/A
log(y/s + 1/(4α)) mean FULL + 0.01 N/A
log(y/s + 1/(4α)) 100 FULL + 0.01 N/A
log(y/s + 1/(4α)) 10 FULL + 0.1 N/A
log(y/s + 1/(4α)) 1000 FULL + 0 N/A
log(y/s + 1/(4α)) mean FULL + 0.01 N/A
log(y/s + 1/(4α)) 10 FULL + 0.05 N/A
log(y/s + 1) 10 FULL + N/A N/A
Analytic pearson N/A FULL - 0.01 10
log(y/s + 1/(4α)) mean FULL + 0 N/A
log(y/s + 1/(4α)) 10 FULL + 0.01 N/A
Pearson N/A FULL - 0.01 10
358 additional rows not shown
Pearson N/A HVG - True sqrt_n
Pearson N/A HVG - True none
y 100 HVG - N/A N/A
Random quantile N/A HVG + True N/A
Pearson N/A HVG - True none
Random quantile N/A HVG - True N/A
Transformation
Size factor
Feature
Z-score
Overdispersion
Clip
NC
SF
GSR
Overall ranking
Size factor -NA -MEAN -10 -100 -1,000
Feature -HVG -FULL
Overdispersion -NA -True -0 -0.05
Z-score -+ Scaled -– Unscaled
Clip -N/A -sqrt_n -none -10 -sqrt_30
0 0.2 0.4 0.6 0.8 1
193 1
b
Analysis of transformation
Size factor
Delta method
GLM
Lat Expr
y
ns: p>0.05
*: p<0.05
**: p<0.01
***: p<0.001
****: p<0.0001
Overall score
Top-10 composition (%)
100%
80%
MERFISH
STARmap
c
Best preprocessing strategy of each transformation
Transformation
MERFISH
STARmap
Delta Method
log(y/s + 1) 2 2
log(y/s + 1/(4α)) 1 1
acosh(2αy/s + 1) 5
Scran
Deseq2
TMM
Glm Residuals
Pearson 4
SCTransform
Pearson analytic 3 3
Random quantile
Lat Expr
Sanity
Normaliser
Negetive
y 4 5
Ranking
14 1
d
Comparison of best vs. worst preprocessing for each transformation
Data: MOB STARmap
Best
Worst
Overall rank sum
log(y/s + 1)
log(y/s + 1/(4α))
acosh(2αy/s + 1)
Scran
Deseq2
TMM
Pearson
SCTransform
Analytic pearson
Random quantile
Sanity
Normaliser
y
e
Analysis of hyperparameters setting
Size factor
10
100
1,000
Mean
NA
40%
40%
Overdispersion
0
0.01
0.05
0.10
Est.
NA
20%
50%
f
Analysis of refinement operations
Feature
HVG
FULL
100%
100%
Z-score
+
-
100%
80%

Figure S5

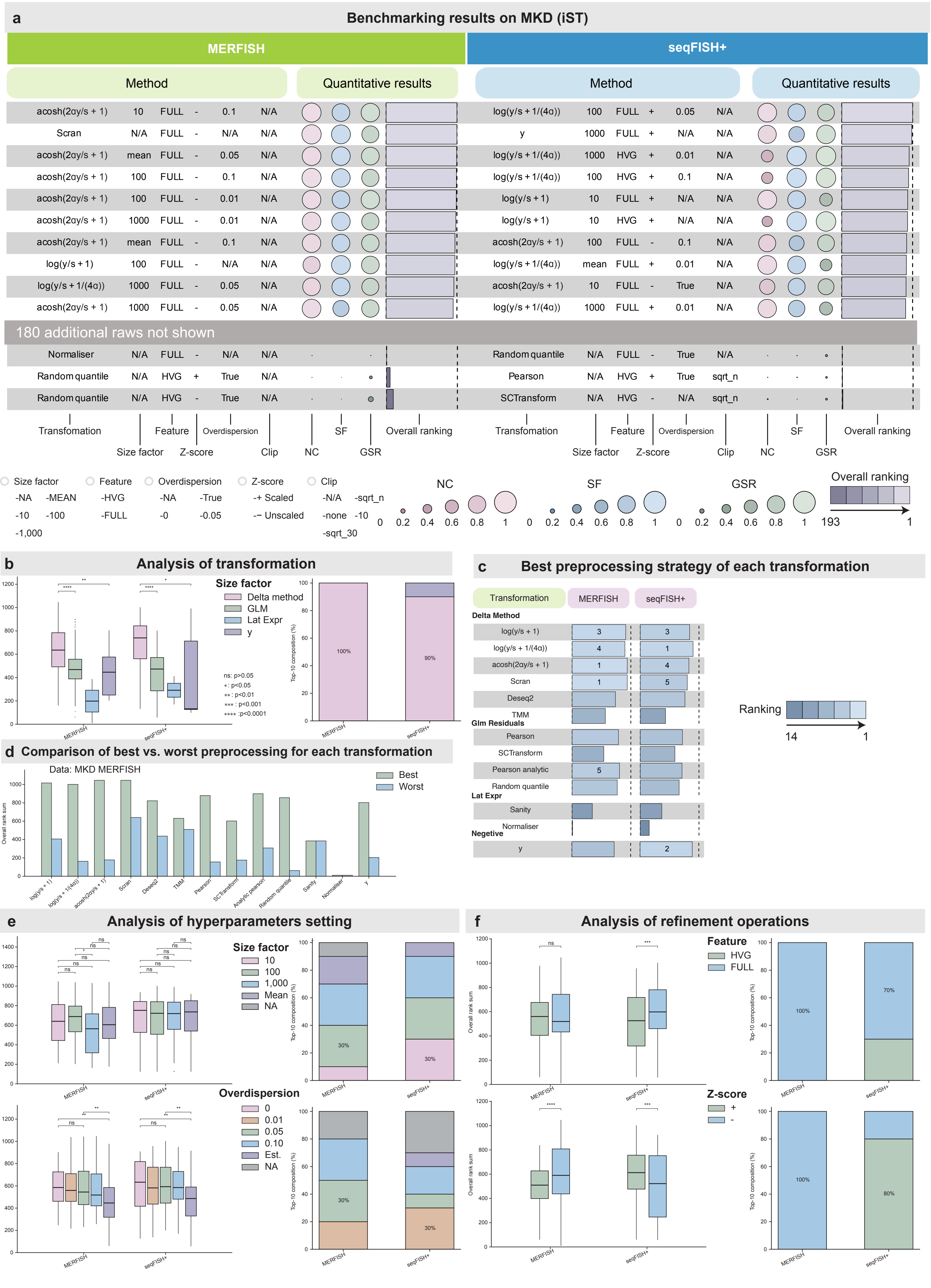
a
Benchmarking results on MKD (iST)
MERFISH
seqFISH+
Method
Quantitative results
acosh(2αy/s + 1) 10 FULL - 0.1 N/A
Scran N/A FULL - N/A N/A
acosh(2αy/s + 1) mean FULL - 0.05 N/A
acosh(2αy/s + 1) 100 FULL - 0.1 N/A
acosh(2αy/s + 1) 100 FULL - 0.01 N/A
acosh(2αy/s + 1) 1000 FULL - 0.01 N/A
acosh(2αy/s + 1) mean FULL - 0.1 N/A
log(y/s + 1) 100 FULL - N/A N/A
log(y/s + 1/(4α)) 1000 FULL - 0.05 N/A
acosh(2αy/s + 1) 1000 FULL - 0.05 N/A
log(y/s + 1/(4α)) 100 FULL + 0.05 N/A
y 1000 FULL + N/A N/A
log(y/s + 1/(4α)) 1000 HVG + 0.01 N/A
log(y/s + 1/(4α)) 100 HVG + 0.1 N/A
log(y/s + 1) 10 FULL + N/A N/A
log(y/s + 1) 10 HVG + N/A N/A
acosh(2αy/s + 1) 100 FULL - 0.1 N/A
log(y/s + 1/(4α)) mean FULL + 0.01 N/A
acosh(2αy/s + 1) 10 FULL - True N/A
log(y/s + 1/(4α)) 1000 FULL + 0.01 N/A
180 additional raws not shown
Normaliser N/A FULL - N/A N/A
Random quantile N/A HVG + True N/A
Random quantile N/A HVG - True N/A
Random quantile N/A FULL - True N/A
Pearson N/A HVG + True sqrt_n
SCTransform N/A HVG - N/A sqrt_n
Transfomation
Size factor
Feature
Z-score
Overdispersion
Clip
NC
SF
GSR
Overall ranking
Size factor -NA -MEAN -10 -100 -1,000
Feature -HVG -FULL
Overdispersion -NA -True -0 -0.05
Z-score -+ Scaled -– Unscaled
Clip -N/A -sqrt_n -none -10 -sqrt_30
NC 0 0.2 0.4 0.6 0.8 1
SF 0 0.2 0.4 0.6 0.8 1
GSR 0 0.2 0.4 0.6 0.8 1
Overall ranking 193 1
b
Analysis of transformation
Size factor
Delta method
GLM
Lat Expr
y
ns: p>0.05
*: p<0.05
**: p<0.01
***: p<0.001
****: p<0.0001
Overall rank sum
Top-10 composition (%)
100%
90%
MERFISH
seqFISH+
c
Best preprocessing strategy of each transformation
Transformation
MERFISH
seqFISH+
Delta Method
log(y/s + 1) 3 3
log(y/s + 1/(4α)) 4 1
acosh(2αy/s + 1) 1 4
Scran 1 5
Deseq2
TMM
Glm Residuals
Pearson
SCTransform
Pearson analytic 5
Random quantile
Lat Expr
Sanity
Normaliser
Negetive
y 2
Ranking
14 1
d
Comparison of best vs. worst preprocessing for each transformation
Data: MKD MERFISH
Best
Worst
Overall rank sum
log(y/s + 1)
log(y/s + 1/(4α))
acosh(2αy/s + 1)
Scran
Deseq2
TMM
Pearson
SCTransform
Analytic pearson
Random quantile
Sanity
Normaliser
y
e
Analysis of hyperparameters setting
Size factor
10
100
1,000
Mean
NA
30%
Overdispersion
0
0.01
0.05
0.10
Est.
NA
f
Analysis of refinement operations
Feature
HVG
FULL
100%
70%
Z-score
+
-
100%
80%

### Figure S6

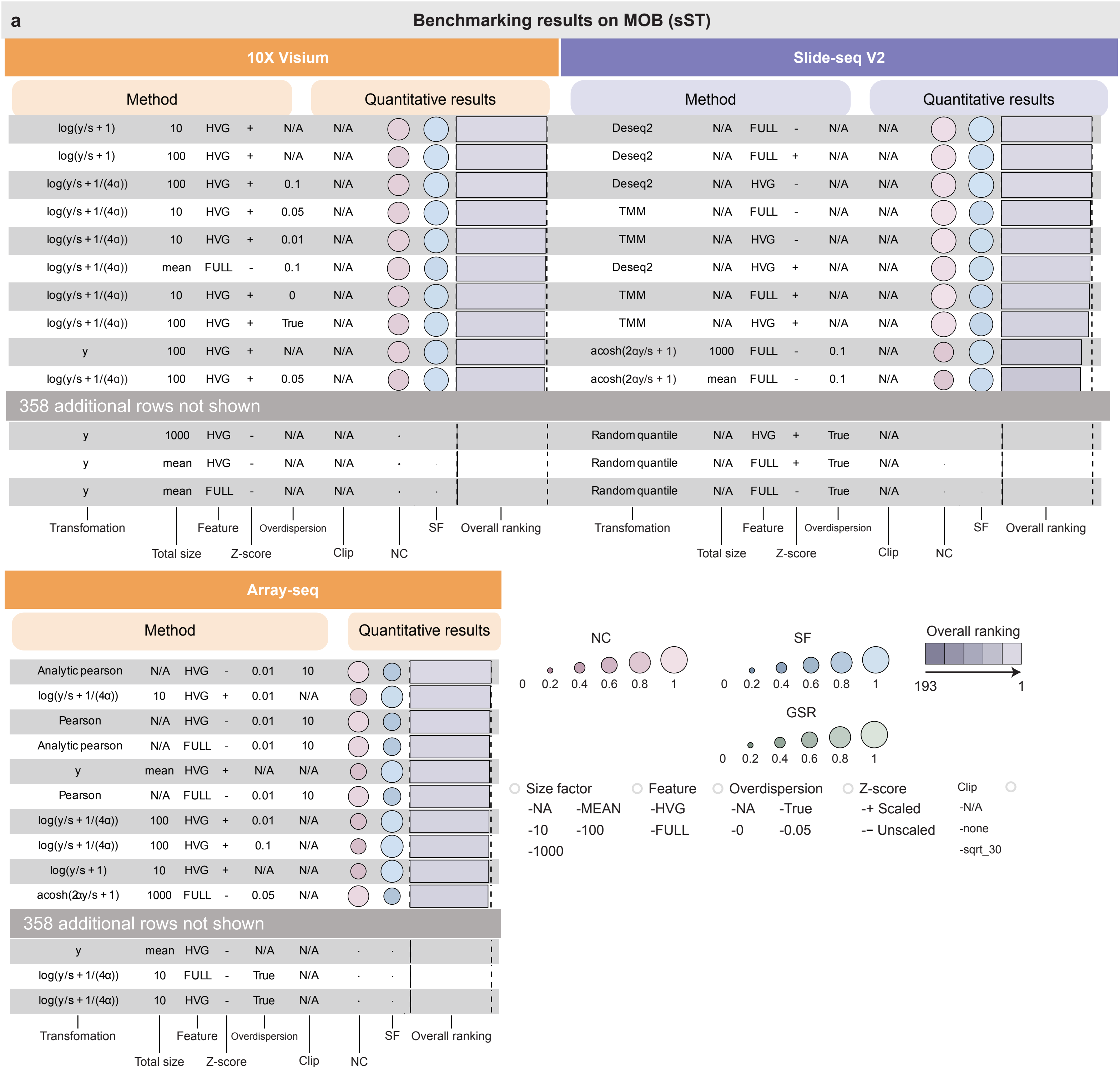
a
Benchmarking results on MOB (sST)
10X Visium
Method
Quantitative results
log(y/s + 1) 10 HVG + N/A N/A
log(y/s + 1) 100 HVG + N/A N/A
log(y/s + 1/(4α)) 100 HVG + 0.1 N/A
log(y/s + 1/(4α)) 10 HVG + 0.05 N/A
log(y/s + 1/(4α)) 10 HVG + 0.01 N/A
log(y/s + 1/(4α)) mean FULL - 0.1 N/A
log(y/s + 1/(4α)) 10 HVG + 0 N/A
log(y/s + 1/(4α)) 100 HVG + True N/A
y 100 HVG + N/A N/A
log(y/s + 1/(4α)) 100 HVG + 0.05 N/A
358 additional rows not shown
y 1000 HVG - N/A N/A
y mean HVG - N/A N/A
y mean FULL - N/A N/A
Transfomation
Total size
Feature
Z-score
Overdispersion
Clip
NC
SF
Overall ranking
Slide-seq V2
Method
Quantitative results
Deseq2 N/A FULL - N/A N/A
Deseq2 N/A FULL + N/A N/A
Deseq2 N/A HVG - N/A N/A
TMM N/A FULL - N/A N/A
TMM N/A HVG - N/A N/A
Deseq2 N/A HVG + N/A N/A
TMM N/A FULL + N/A N/A
TMM N/A HVG + N/A N/A
acosh(2αy/s + 1) 1000 FULL - 0.1 N/A
acosh(2αy/s + 1) mean FULL - 0.1 N/A
358 additional rows not shown
Random quantile N/A HVG + True N/A
Random quantile N/A FULL + True N/A
Random quantile N/A FULL - True N/A
Transfomation
Total size
Feature
Z-score
Overdispersion
Clip
NC
SF
Overall ranking
Array-seq
Method
Quantitative results
Analytic pearson N/A HVG - 0.01 10
log(y/s + 1/(4α)) 10 HVG + 0.01 N/A
Pearson N/A HVG - 0.01 10
Analytic pearson N/A FULL - 0.01 10
y mean HVG + N/A N/A
Pearson N/A FULL - 0.01 10
log(y/s + 1/(4α)) 100 HVG + 0.01 N/A
log(y/s + 1/(4α)) 100 HVG + 0.1 N/A
log(y/s + 1) 10 HVG + N/A N/A
acosh(2αy/s + 1) 1000 FULL - 0.05 N/A
358 additional rows not shown
y mean HVG - N/A N/A
log(y/s + 1/(4α)) 10 FULL - True N/A
log(y/s + 1/(4α)) 10 HVG - True N/A
Transfomation
Total size
Feature
Z-score
Overdispersion
Clip
NC
SF
Overall ranking
NC
0 0.2 0.4 0.6 0.8 1
SF
0 0.2 0.4 0.6 0.8 1
Overall ranking
193 1
GSR
0 0.2 0.4 0.6 0.8 1
Size factor
-NA -MEAN
-10 -100
-1000
Feature
-HVG
-FULL
Overdispersion
-NA -True
-0 -0.05
Z-score
-+ Scaled
-– Unscaled
Clip
-N/A
-none
-sqrt_30

Figure S7

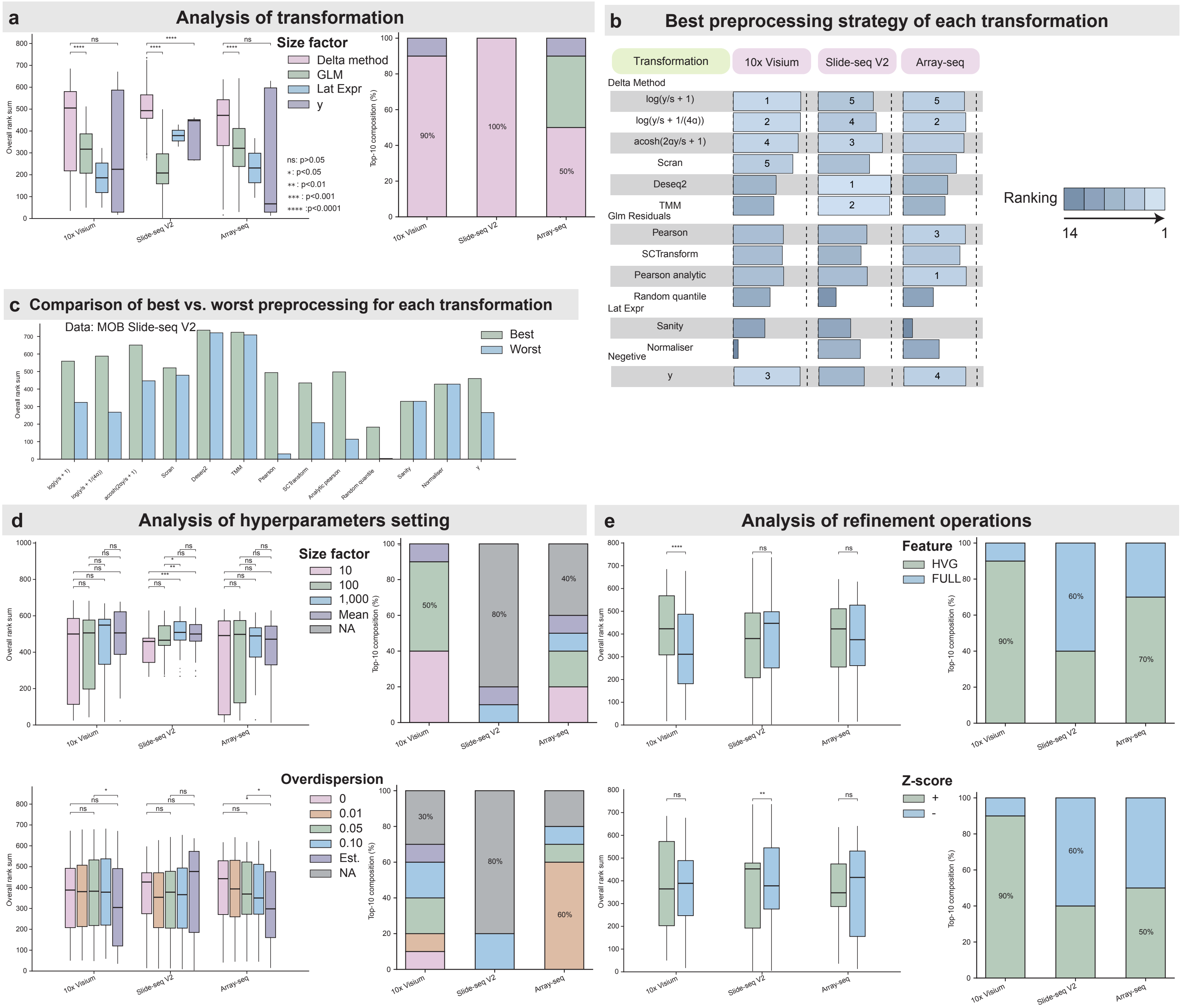

a
Analysis of transformation
Size factor
Delta method
GLM
Lat Expr
y
ns: p>0.05
*: p<0.05
**: p<0.01
***: p<0.001
****: p<0.0001
Overall rank sum
Top-10 composition (%)
10x Visium
Slide-seq V2
Array-seq
90%
100%
50%
b
Best preprocessing strategy of each transformation
Transformation
10x Visium
Slide-seq V2
Array-seq
Delta Method
log(y/s + 1)
log(y/s + 1/(4α))
acosh(2αy/s + 1)
Scran
Deseq2
TMM
Glm Residuals
Pearson
SCTransform
Pearson analytic
Random quantile
Lat Expr
Sanity
Normaliser
Negetive
y
Ranking
14
1
c
Comparison of best vs. worst preprocessing for each transformation
Data: MOB Slide-seq V2
Best
Worst
Overall rank sum
d
Analysis of hyperparameters setting
Size factor
10
100
1,000
Mean
NA
Overdispersion
0
0.01
0.05
0.10
Est.
NA
50%
80%
40%
30%
60%
e
Analysis of refinement operations
Feature
HVG
FULL
Z-score
+
-
90%
60%
70%
50%

#### Figure S8

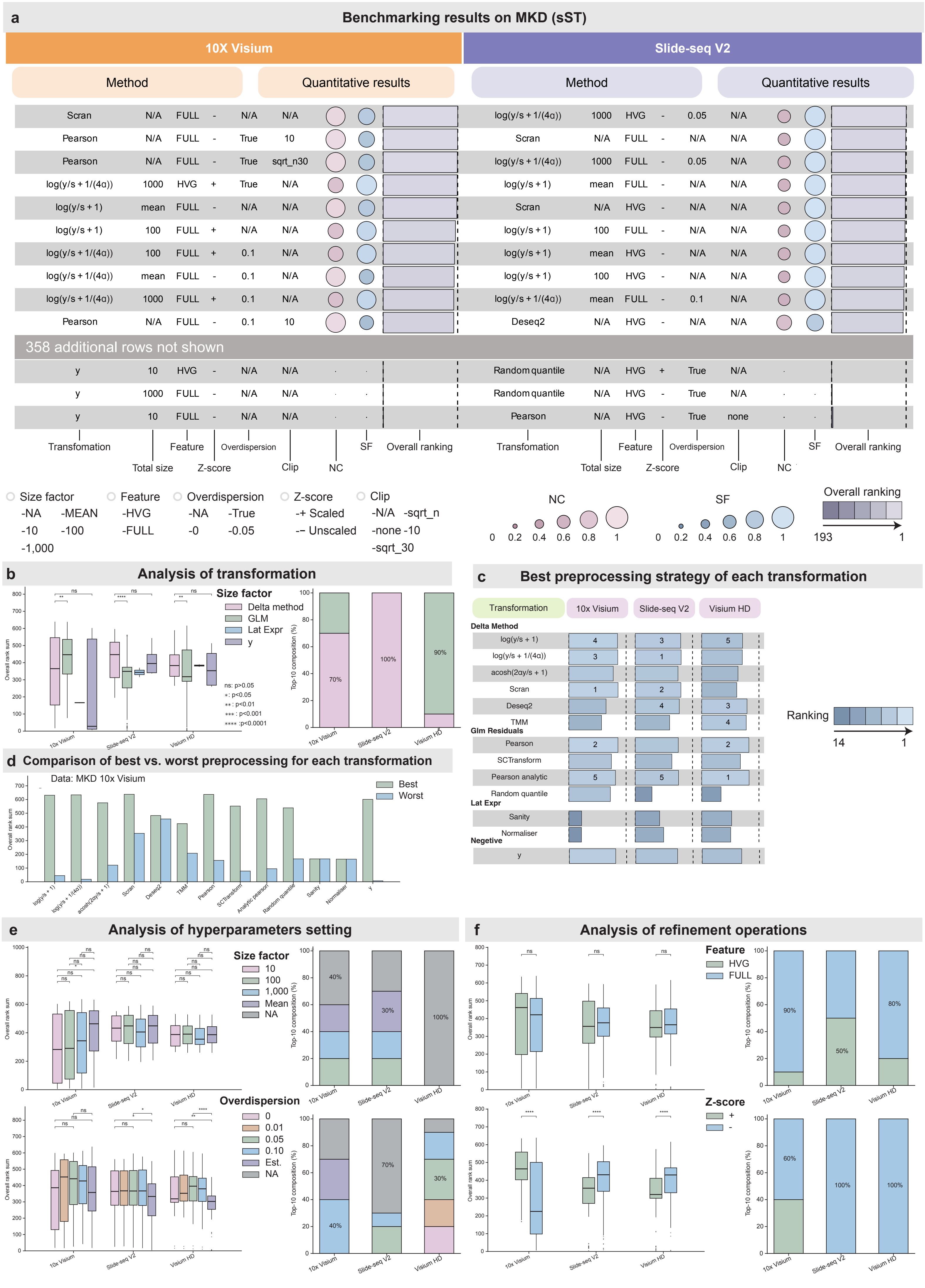
a
Benchmarking results on MKD (sST)
10X Visium
Slide-seq V2
Method
Quantitative results
Scran N/A FULL - N/A N/A
Pearson N/A FULL - True 10
Pearson N/A FULL - True sqrt_n30
log(y/s + 1/(4α)) 1000 HVG + True N/A
log(y/s + 1) mean FULL - N/A N/A
log(y/s + 1) 100 FULL + N/A N/A
log(y/s + 1/(4α)) 100 FULL + 0.1 N/A
log(y/s + 1/(4α)) mean FULL - 0.1 N/A
log(y/s + 1/(4α)) 1000 FULL + 0.1 N/A
Pearson N/A FULL - 0.1 10
log(y/s + 1/(4α)) 1000 HVG - 0.05 N/A
Scran N/A FULL - N/A N/A
log(y/s + 1/(4α)) 1000 FULL - 0.05 N/A
log(y/s + 1) mean FULL - N/A N/A
Scran N/A HVG - N/A N/A
log(y/s + 1) 100 FULL - N/A N/A
log(y/s + 1) mean HVG - N/A N/A
log(y/s + 1) 100 HVG - N/A N/A
log(y/s + 1/(4α)) mean FULL - 0.1 N/A
Deseq2 N/A HVG - N/A N/A
358 additional rows not shown
y 10 HVG - N/A N/A
y 1000 FULL - N/A N/A
y 10 FULL - N/A N/A
Random quantile N/A HVG + True N/A
Random quantile N/A HVG - True N/A
Pearson N/A HVG - True none
Transfomation
Total size
Feature
Z-score
Overdispersion
Clip
NC
SF
Overall ranking
Size factor -NA -MEAN -10 -100 -1,000
Feature -HVG -FULL
Overdispersion -NA -True -0 -0.05
Z-score -+ Scaled -– Unscaled
Clip -N/A -sqrt_n -none -10 -sqrt_30
NC 0 0.2 0.4 0.6 0.8 1
SF 0 0.2 0.4 0.6 0.8 1
Overall ranking 193 1
b
Analysis of transformation
Size factor
Delta method
GLM
Lat Expr
y
ns: p>0.05
*: p<0.05
**: p<0.01
***: p<0.001
****: p<0.0001
Overall rank sum
Top-10 composition (%)
70%
100%
90%
10x Visium
Slide-seq V2
Visium HD
c
Best preprocessing strategy of each transformation
Transformation
10x Visium
Slide-seq V2
Visium HD
Delta Method
log(y/s + 1) 4 3 5
log(y/s + 1/(4α)) 3 1
acosh(2αy/s + 1)
Scran 1 2
Deseq2 4 3
TMM 4
Glm Residuals
Pearson 2 2
SCTransform
Pearson analytic 5 5 1
Random quantile
Lat Expr
Sanity
Normaliser
Negetive
y
Ranking
14 1
d
Comparison of best vs. worst preprocessing for each transformation
Data: MKD 10x Visium
Best
Worst
Overall rank sum
log(y/s + 1)
log(y/s + 1/(4α))
acosh(2αy/s + 1)
Scran
Deseq2
TMM
Pearson
SCTransform
Analytic pearson
Random quantile
Sanity
Normaliser
y
e
Analysis of hyperparameters setting
Size factor
10
100
1,000
Mean
NA
40%
30%
100%
Overdispersion
0
0.01
0.05
0.10
Est.
NA
40%
70%
30%
Overall rank sum
Top-10 composition (%)
10x Visium
Slide-seq V2
Visium HD
f
Analysis of refinement operations
Feature
HVG
FULL
90%
50%
80%
Z-score
+
-
60%
100%
100%
Overall rank sum
Top-10 composition (%)
10x Visium
Slide-seq V2
Visium HD

#### Figure S9

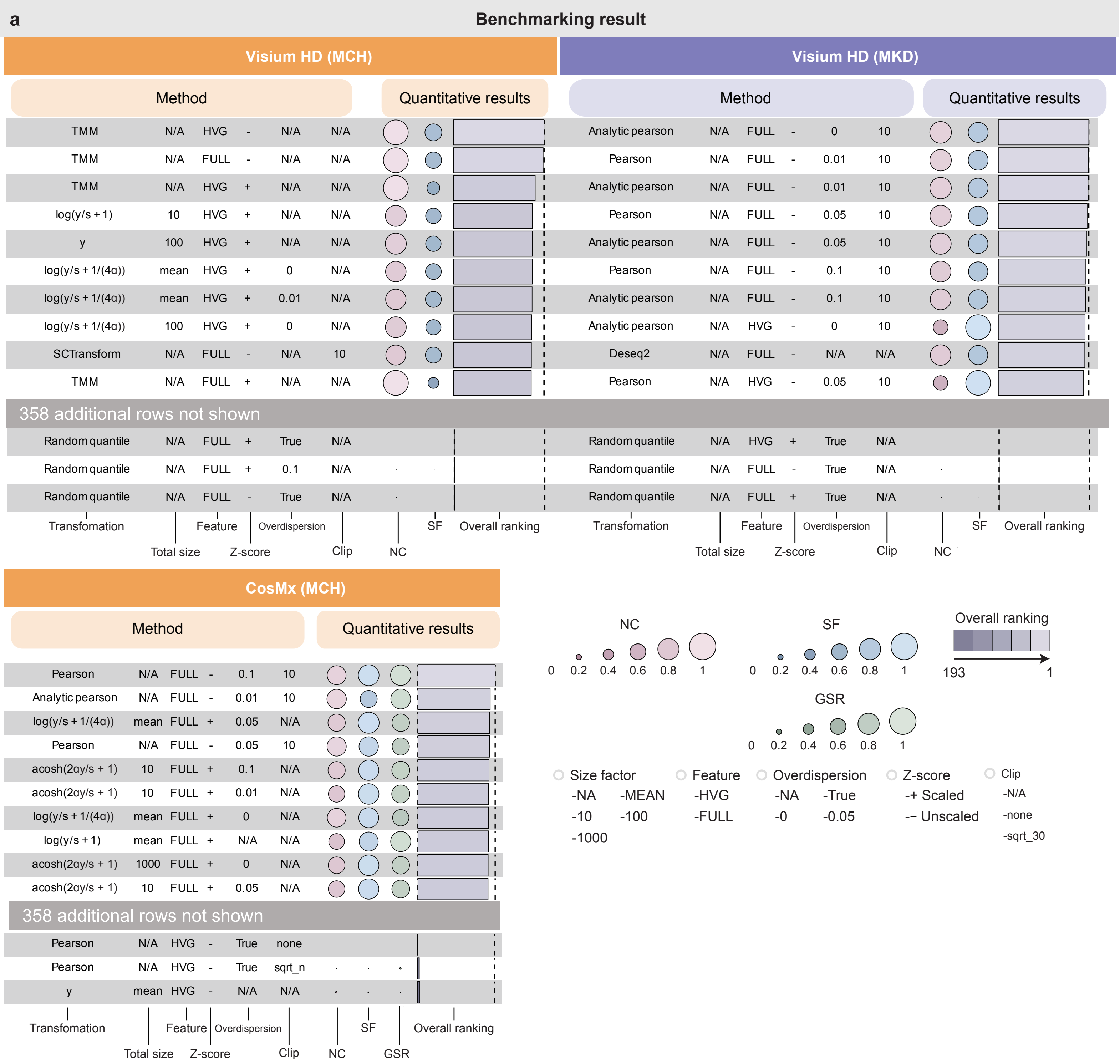
a
Benchmarking result
Visium HD (MCH)
Method
Quantitative results
TMM N/A HVG - N/A N/A
TMM N/A FULL - N/A N/A
TMM N/A HVG + N/A N/A
log(y/s + 1) 10 HVG + N/A N/A
y 100 HVG + N/A N/A
log(y/s + 1/(4α)) mean HVG + 0 N/A
log(y/s + 1/(4α)) mean HVG + 0.01 N/A
log(y/s + 1/(4α)) 100 HVG + 0 N/A
SCTransform N/A FULL - N/A 10
TMM N/A FULL + N/A N/A
358 additional rows not shown
Random quantile N/A FULL + True N/A
Random quantile N/A FULL + 0.1 N/A
Random quantile N/A FULL - True N/A
Transfomation
Total size
Feature
Z-score
Overdispersion
Clip
NC
SF
Overall ranking
Visium HD (MKD)
Method
Quantitative results
Analytic pearson N/A FULL - 0 10
Pearson N/A FULL - 0.01 10
Analytic pearson N/A FULL - 0.01 10
Pearson N/A FULL - 0.05 10
Analytic pearson N/A FULL - 0.05 10
Pearson N/A FULL - 0.1 10
Analytic pearson N/A FULL - 0.1 10
Analytic pearson N/A HVG - 0 10
Deseq2 N/A FULL - N/A N/A
Pearson N/A HVG - 0.05 10
358 additional rows not shown
Random quantile N/A HVG + True N/A
Random quantile N/A FULL - True N/A
Random quantile N/A FULL + True N/A
Transfomation
Total size
Feature
Z-score
Overdispersion
Clip
NC
SF
Overall ranking
CosMx (MCH)
Method
Quantitative results
Pearson N/A FULL - 0.1 10
Analytic pearson N/A FULL - 0.01 10
log(y/s + 1/(4α)) mean FULL + 0.05 N/A
Pearson N/A FULL - 0.05 10
acosh(2αy/s + 1) 10 FULL + 0.1 N/A
acosh(2αy/s + 1) 10 FULL + 0.01 N/A
log(y/s + 1/(4α)) mean FULL + 0 N/A
log(y/s + 1) mean FULL + N/A N/A
acosh(2αy/s + 1) 1000 FULL + 0 N/A
acosh(2αy/s + 1) 10 FULL + 0.05 N/A
358 additional rows not shown
Pearson N/A HVG - True none
Pearson N/A HVG - True sqrt_n
y mean HVG - N/A N/A
Transfomation
Total size
Feature
Z-score
Overdispersion
Clip
NC
SF
GSR
Overall ranking
NC
0 0.2 0.4 0.6 0.8 1
SF
0 0.2 0.4 0.6 0.8 1
Overall ranking
193 1
GSR
0 0.2 0.4 0.6 0.8 1
Size factor
-NA -MEAN
-10 -100
-1000
Feature
-HVG
-FULL
Overdispersion
-NA -True
-0 -0.05
Z-score
-+ Scaled
-– Unscaled
Clip
-N/A
-none
-sqrt_30

#### Figure S10

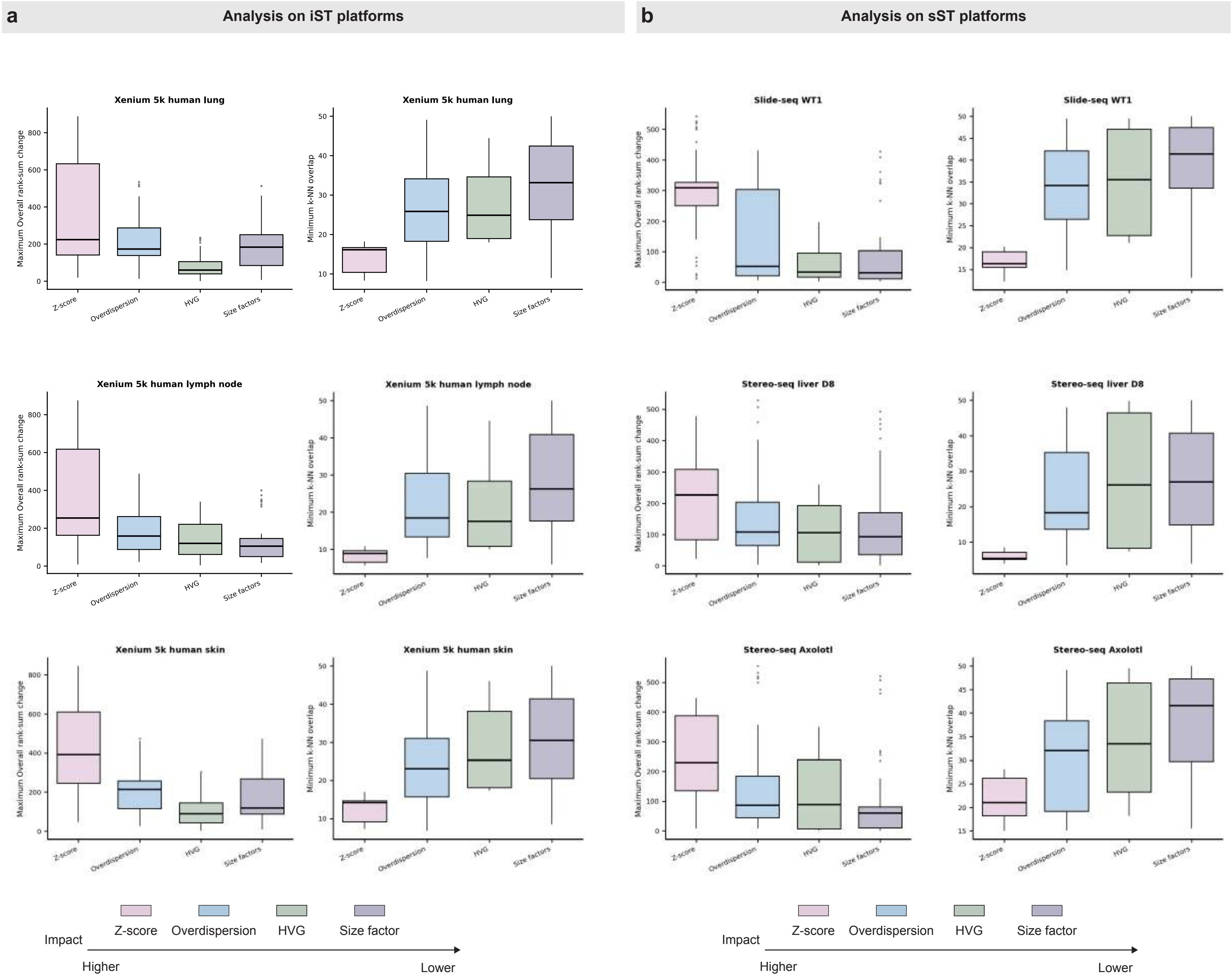

a
Analysis on iST platforms
b
Analysis on sST platforms
Xenium 5k human lung
Xenium 5k human lymph node
Xenium 5k human skin
Slide-seq WT1
Stereo-seq liver D8
Stereo-seq Axolotl
Maximum Overall rank-sum change
Minimum k-NN overlap
Z-score
Overdispersion
HVG
Size factors
Impact
Z-score
Overdispersion
HVG
Size factor
Higher
Lower

### Figure S11

#### a Schematic of ANCOVA

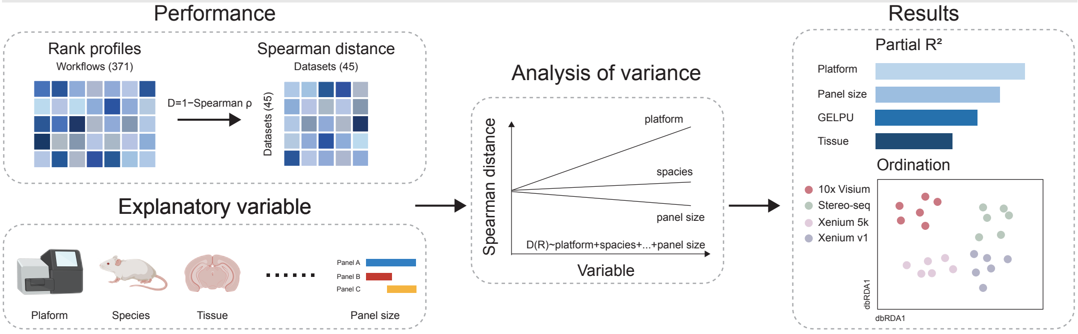


#### b Explanatory variable table

| | |
|---|---|
| Categorical | Platform,Tissue,Species |
| Continuous | Cell number,GELPU, Panel size |

#### c Ordination of preprocessing-ranking profiles

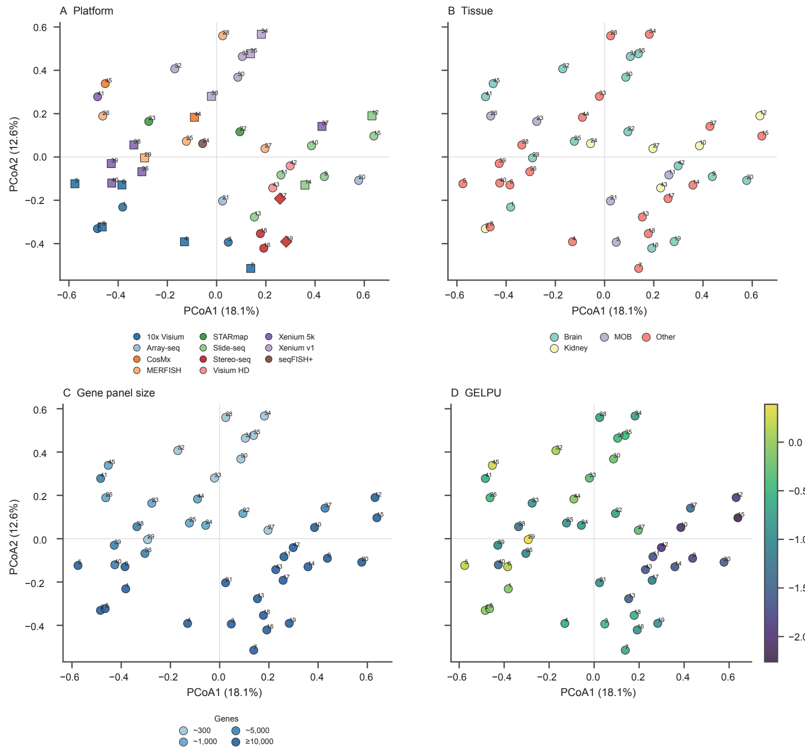


#### d Variance partitioning of preprocessing-ranking profiles

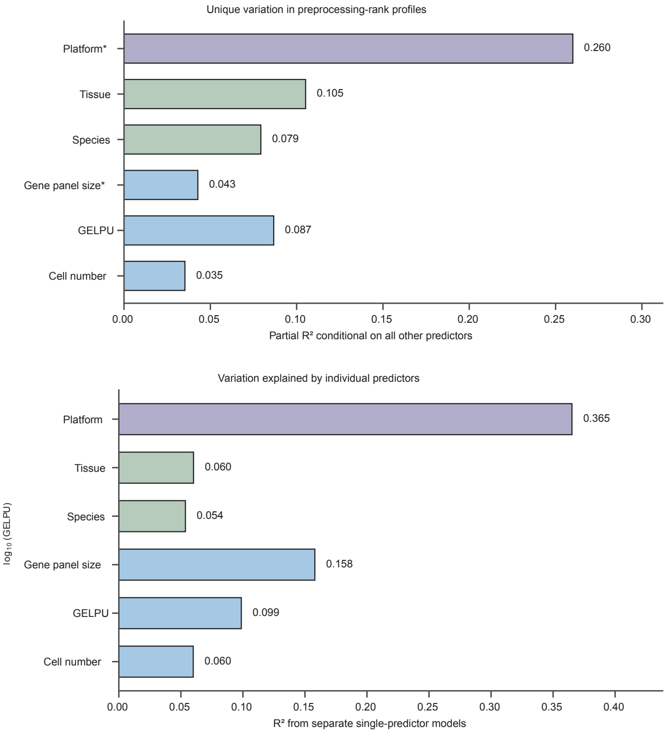

#### Figure S12

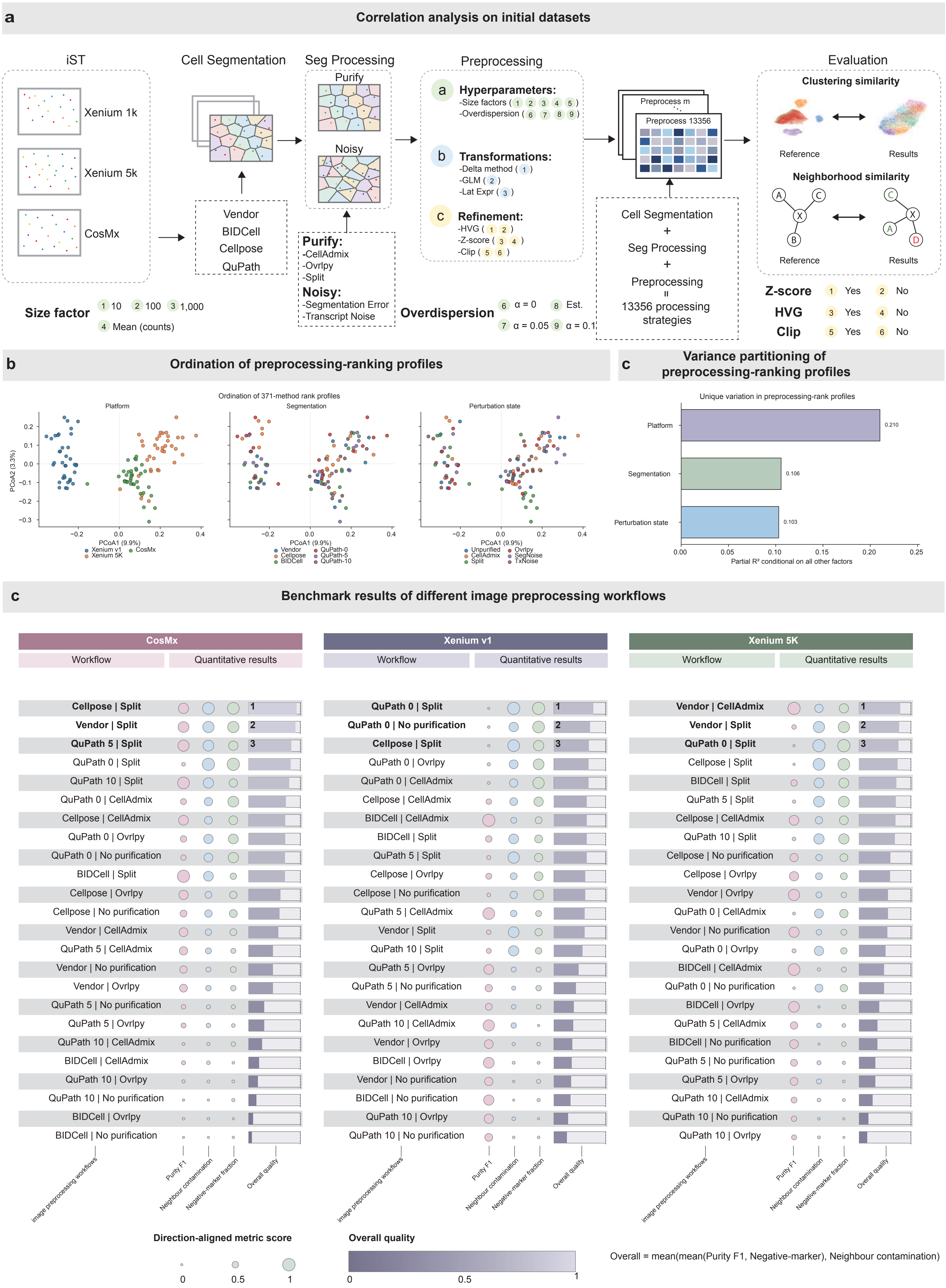

a
Correlation analysis on initial datasets
iST
Xenium 1k
Xenium 5k
CosMx
Cell Segmentation
Vendor
BIDCell
Cellpose
QuPath
Seg Processing
Purify
Noisy
Purify:
-CellAdmix
-Ovrlpy
-Split
Noisy:
-Segmentation Error
-Transcript Noise
Preprocessing
Hyperparameters:
-Size factors ( 1 2 3 4 5 )
-Overdispersion ( 6 7 8 9 )
Transformations:
-Delta method ( 1 )
-GLM ( 2 )
-Lat Expr ( 3 )
Refinement:
-HVG ( 1 2 )
-Z-score ( 3 4 )
-Clip ( 5 6 )
Preprocess m
Preprocess 13356
Cell Segmentation
+
Seg Processing
+
Preprocessing
=
13356 processing strategies
Evaluation
Clustering similarity
Reference
Results
Neighborhood similarity
Reference
Results
Size factor
1 10 2 100 3 1,000
4 Mean (counts)
Overdispersion
6 α = 0 8 Est.
7 α = 0.05 9 α = 0.1
Z-score 1 Yes 2 No
HVG 3 Yes 4 No
Clip 5 Yes 6 No
b
Ordination of preprocessing-ranking profiles
Ordination of 371-method rank profiles
Platform
Segmentation
Perturbation state
PCoA2 (3.3%)
PCoA1 (9.9%)
Xenium v1
Xenium 5K
CosMx
Vendor
Cellpose
BIDCell
QuPath-0
QuPath-5
QuPath-10
Unpurified
CellAdmix
Split
Ovrlpy
SegNoise
TxNoise
c
Variance partitioning of preprocessing-ranking profiles
Unique variation in preprocessing-rank profiles
Platform 0.210
Segmentation 0.106
Perturbation state 0.103
Partial R² conditional on all other factors
c
Benchmark results of different image preprocessing workflows
CosMx
Workflow
Quantitative results
Cellpose | Split
Vendor | Split
QuPath 5 | Split
QuPath 0 | Split
QuPath 10 | Split
QuPath 0 | CellAdmix
Cellpose | CellAdmix
QuPath 0 | Ovrlpy
QuPath 0 | No purification
BIDCell | Split
Cellpose | Ovrlpy
Cellpose | No purification
Vendor | CellAdmix
QuPath 5 | CellAdmix
Vendor | No purification
Vendor | Ovrlpy
QuPath 5 | No purification
QuPath 5 | Ovrlpy
QuPath 10 | CellAdmix
BIDCell | CellAdmix
QuPath 10 | Ovrlpy
QuPath 10 | No purification
BIDCell | Ovrlpy
BIDCell | No purification
Xenium v1
Workflow
Quantitative results
QuPath 0 | Split
QuPath 0 | No purification
Cellpose | Split
QuPath 0 | Ovrlpy
QuPath 0 | CellAdmix
Cellpose | CellAdmix
BIDCell | CellAdmix
BIDCell | Split
QuPath 5 | Split
Cellpose | Ovrlpy
Cellpose | No purification
QuPath 5 | CellAdmix
Vendor | Split
QuPath 10 | Split
QuPath 5 | Ovrlpy
QuPath 5 | No purification
Vendor | CellAdmix
QuPath 10 | CellAdmix
Vendor | Ovrlpy
BIDCell | Ovrlpy
Vendor | No purification
BIDCell | No purification
QuPath 10 | Ovrlpy
QuPath 10 | No purification
Xenium 5K
Workflow
Quantitative results
Vendor | CellAdmix
Vendor | Split
QuPath 0 | Split
Cellpose | Split
BIDCell | Split
QuPath 5 | Split
Cellpose | CellAdmix
QuPath 10 | Split
Cellpose | No purification
Cellpose | Ovrlpy
Vendor | Ovrlpy
QuPath 0 | CellAdmix
Vendor | No purification
QuPath 0 | Ovrlpy
BIDCell | CellAdmix
QuPath 0 | No purification
BIDCell | Ovrlpy
QuPath 5 | CellAdmix
BIDCell | No purification
QuPath 5 | No purification
QuPath 5 | Ovrlpy
QuPath 10 | CellAdmix
QuPath 10 | No purification
QuPath 10 | Ovrlpy
image preprocessing workflows
Purity F1
Neighbour contamination
Negative-marker fraction
Overall quality
Direction-aligned metric score
0 0.5 1
Overall quality
0 0.5 1
Overall = mean(mean(Purity F1, Negative-marker), Neighbour contamination)

Figure S13

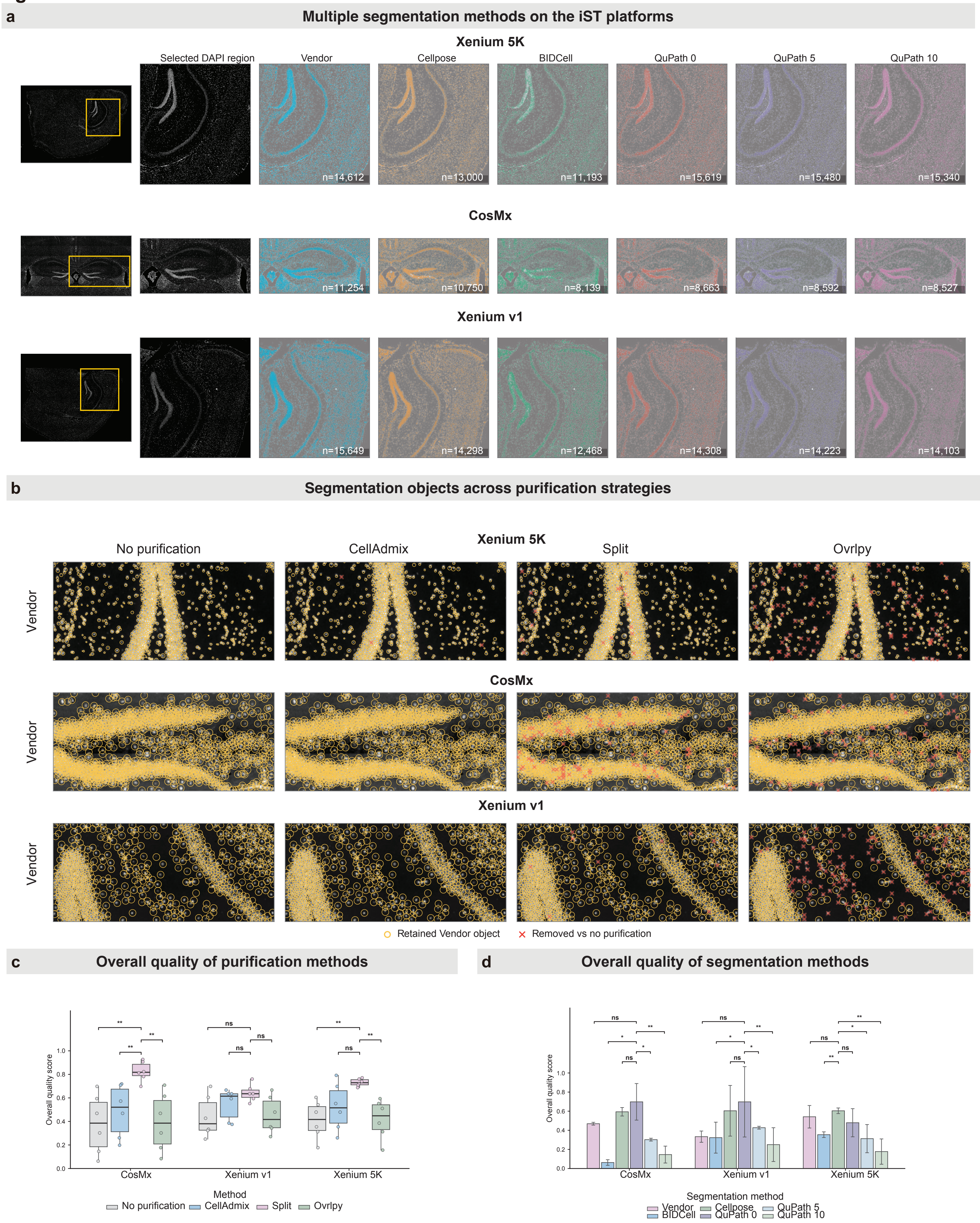
a
Multiple segmentation methods on the iST platforms
Xenium 5K
Selected DAPI region
Vendor
Cellpose
BIDCell
QuPath 0
QuPath 5
QuPath 10
n=14,612
n=13,000
n=11,193
n=15,619
n=15,480
n=15,340
CosMx
n=11,254
n=10,750
n=8,139
n=8,663
n=8,592
n=8,527
Xenium v1
n=15,649
n=14,298
n=12,468
n=14,308
n=14,223
n=14,103
b
Segmentation objects across purification strategies
Xenium 5K
No purification
CellAdmix
Split
Ovrlpy
Vendor
CosMx
Vendor
Xenium v1
Vendor
Retained Vendor object
Removed vs no purification
c
Overall quality of purification methods
Overall quality score
CosMx
Xenium v1
Xenium 5K
Method
No purification
CellAdmix
Split
Ovrlpy
d
Overall quality of segmentation methods
Overall quality score
CosMx
Xenium v1
Xenium 5K
Segmentation method
Vendor
Cellpose
QuPath 5
BIDCell
QuPath 0
QuPath 10

Figure S14

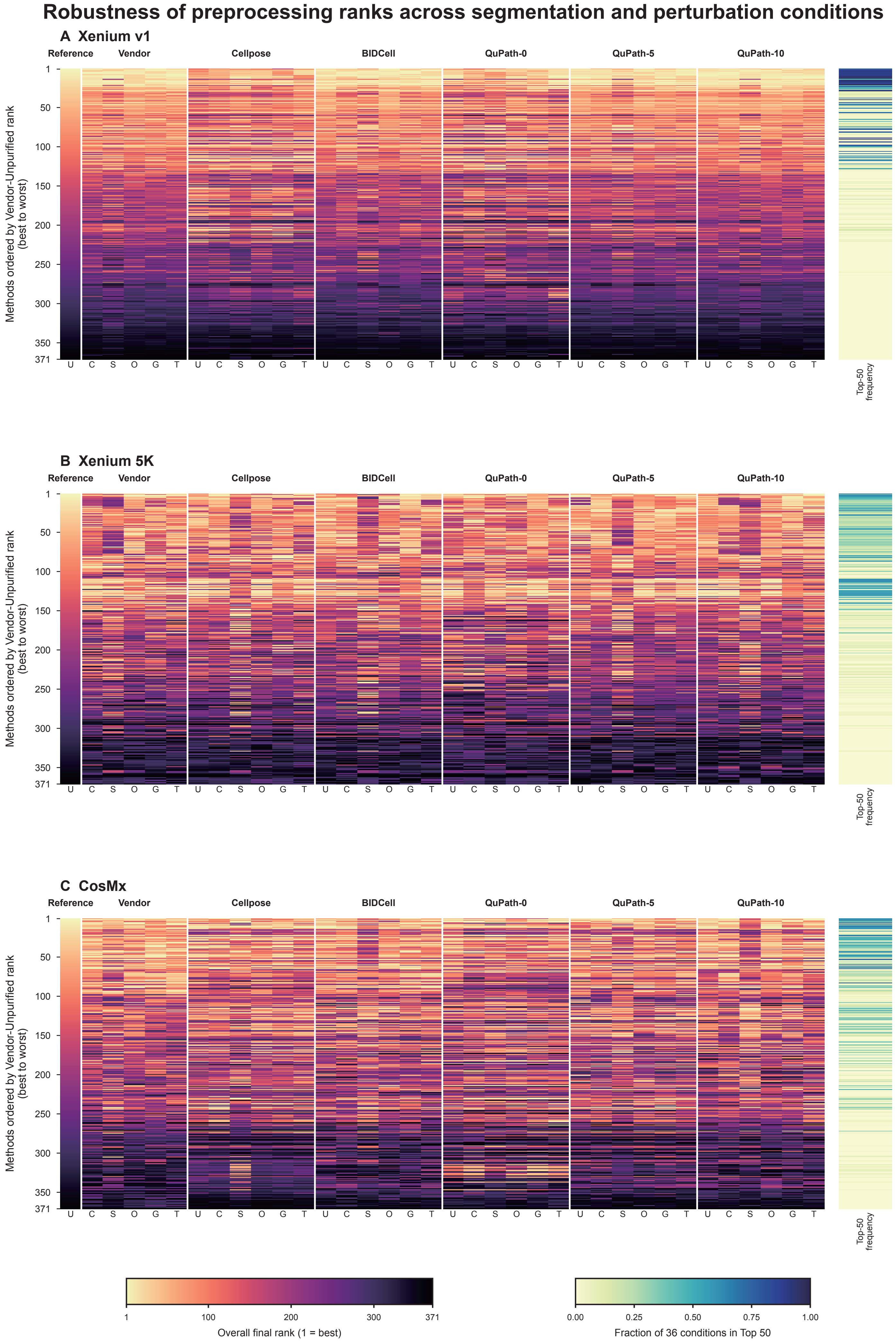


Columns: U=Unpurified, C=CellAdmix, S=Split, O=Ovrlpy, G=SegNoise, T=TxNoise. Rows are fixed by Vendor-Unpurified rank.

#### Figure S15

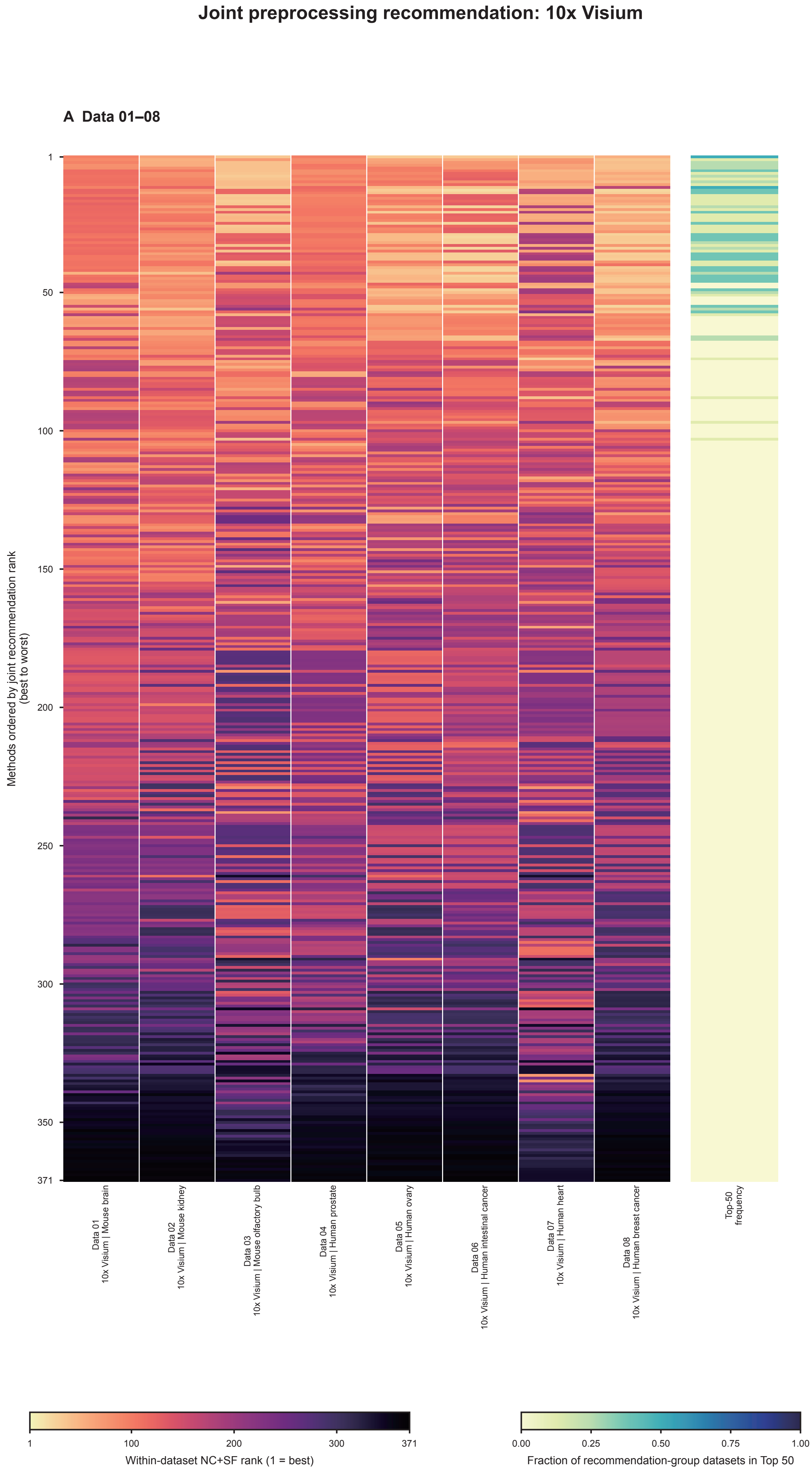


Rows follow the joint recommendation rank: aggregate of mean within-dataset component ranks across all displayed datasets.

Figure S16

#### Joint preprocessing recommendation: Array-seq + Stereo-seq

**A Data 16–21**

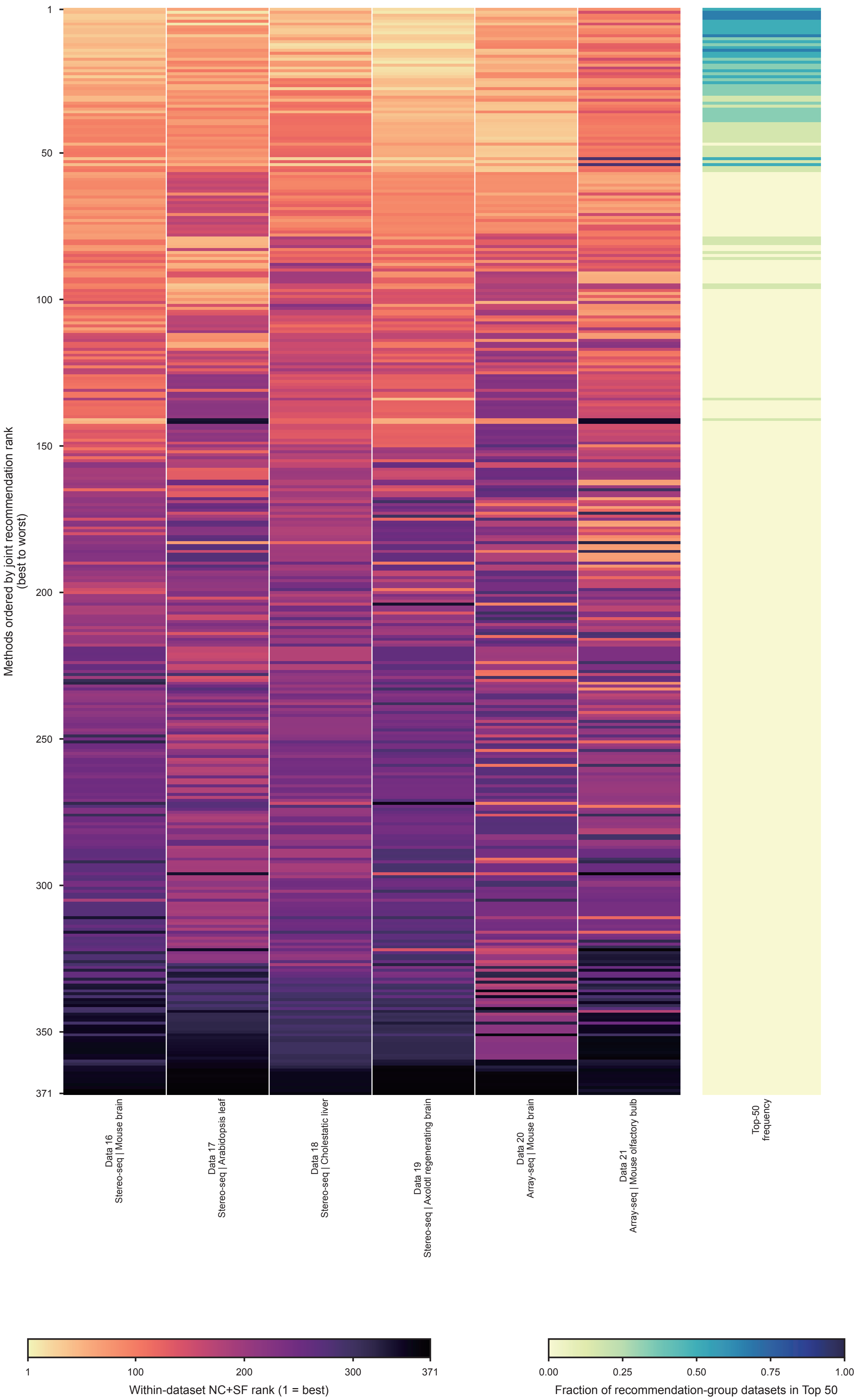


Rows follow the joint recommendation rank: aggregate of mean within-dataset component ranks across all displayed datasets.

#### Figure S17

##### Joint preprocessing recommendation: MERFISH 300

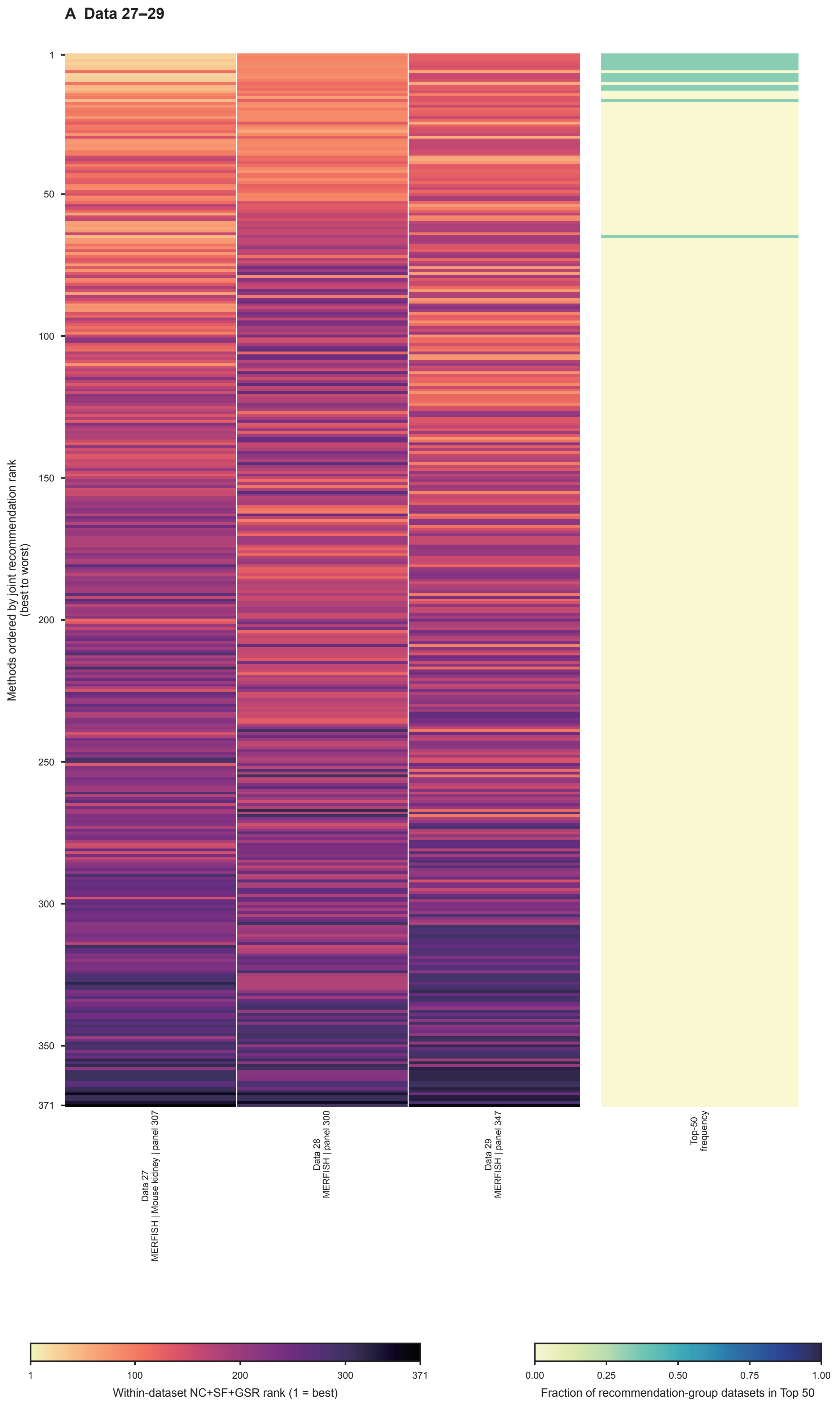


Rows follow the joint recommendation rank: aggregate of mean within-dataset component ranks across all displayed datasets.

#### Figure S18

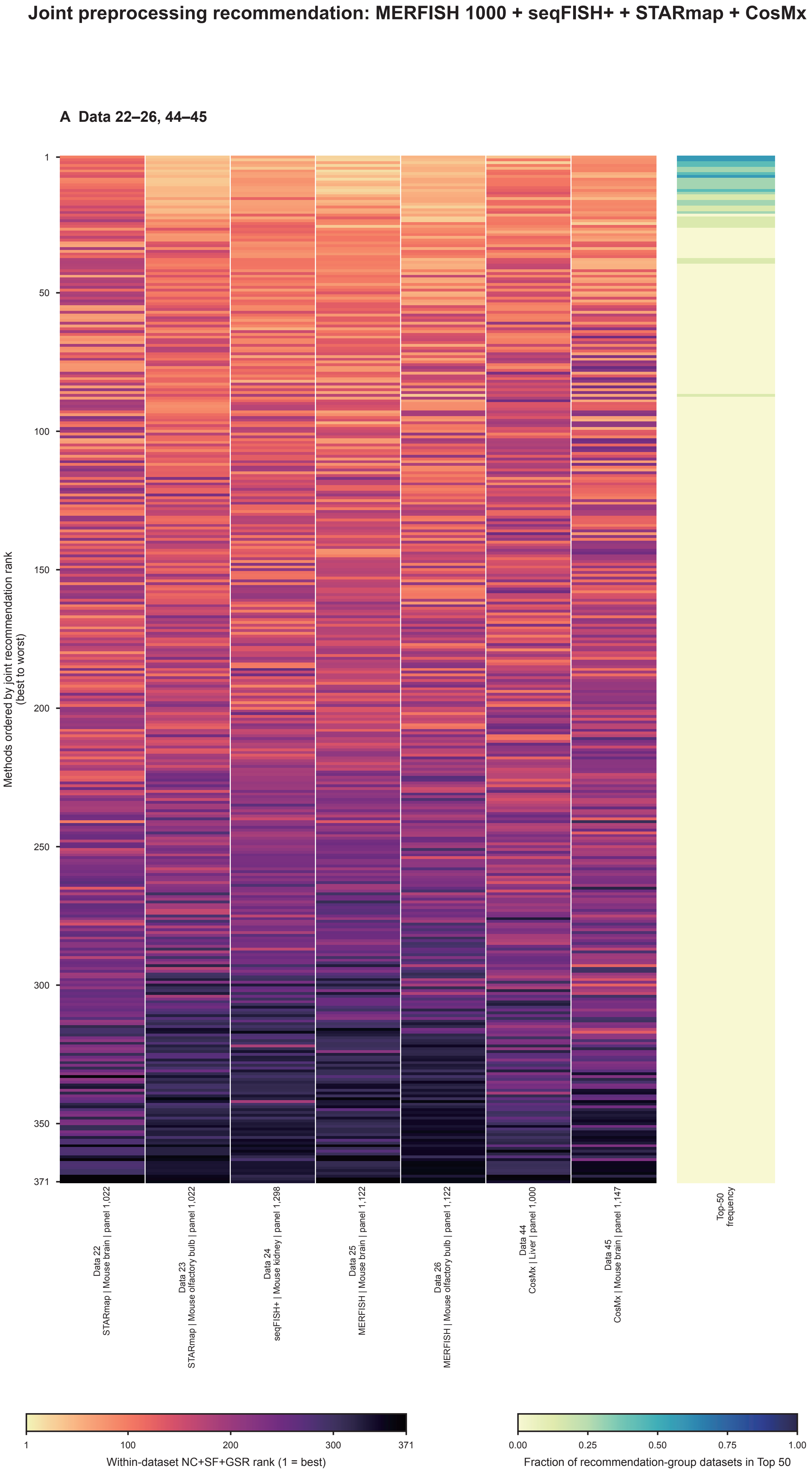


Rows follow the joint recommendation rank: aggregate of mean within-dataset component ranks across all displayed datasets.

Figure S19

#### Joint preprocessing recommendation: Slide-seq V2 + Visium HD

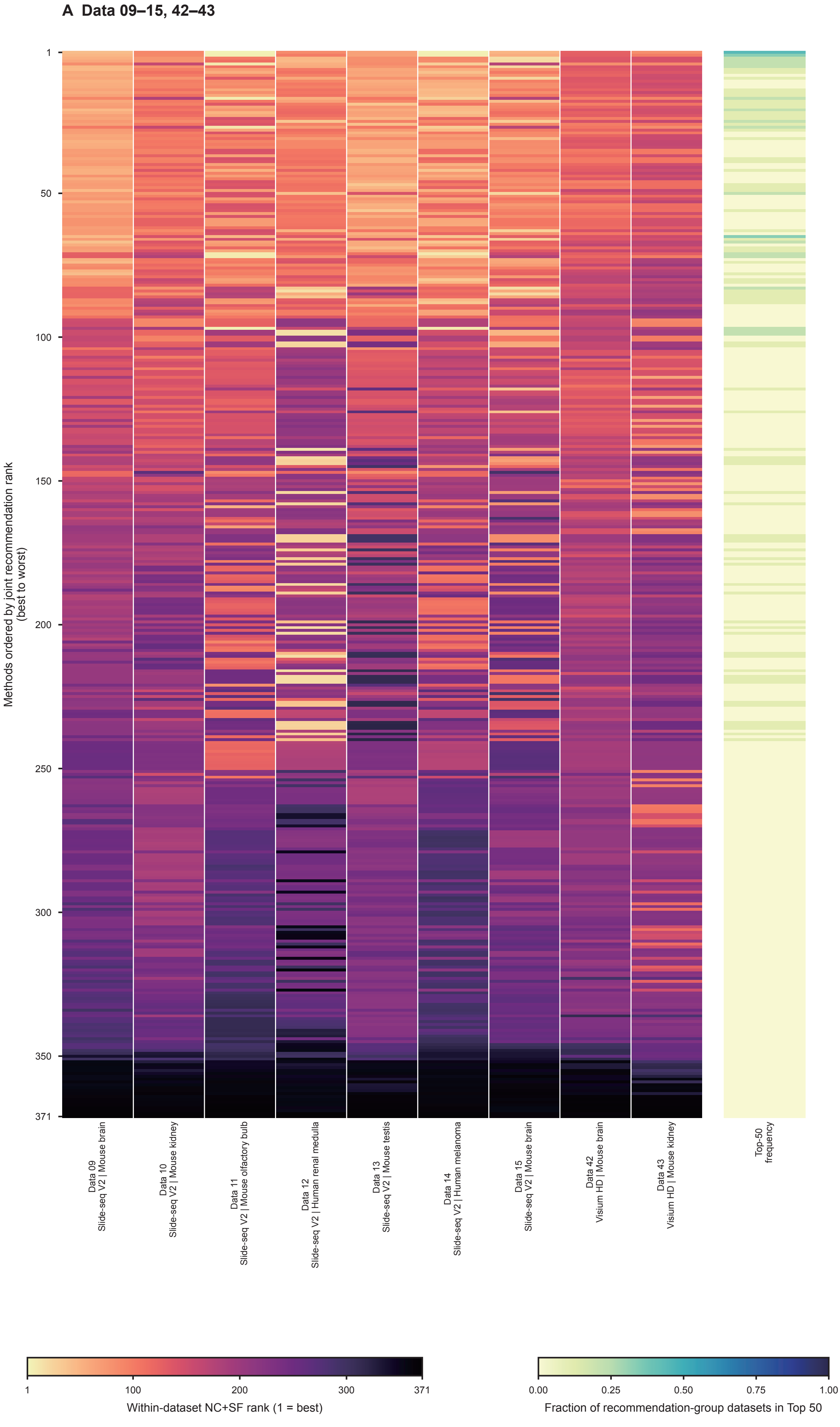


Rows follow the joint recommendation rank: aggregate of mean within-dataset component ranks across all displayed datasets.

### Figure S20

#### Joint preprocessing recommendation: Xenium 5k

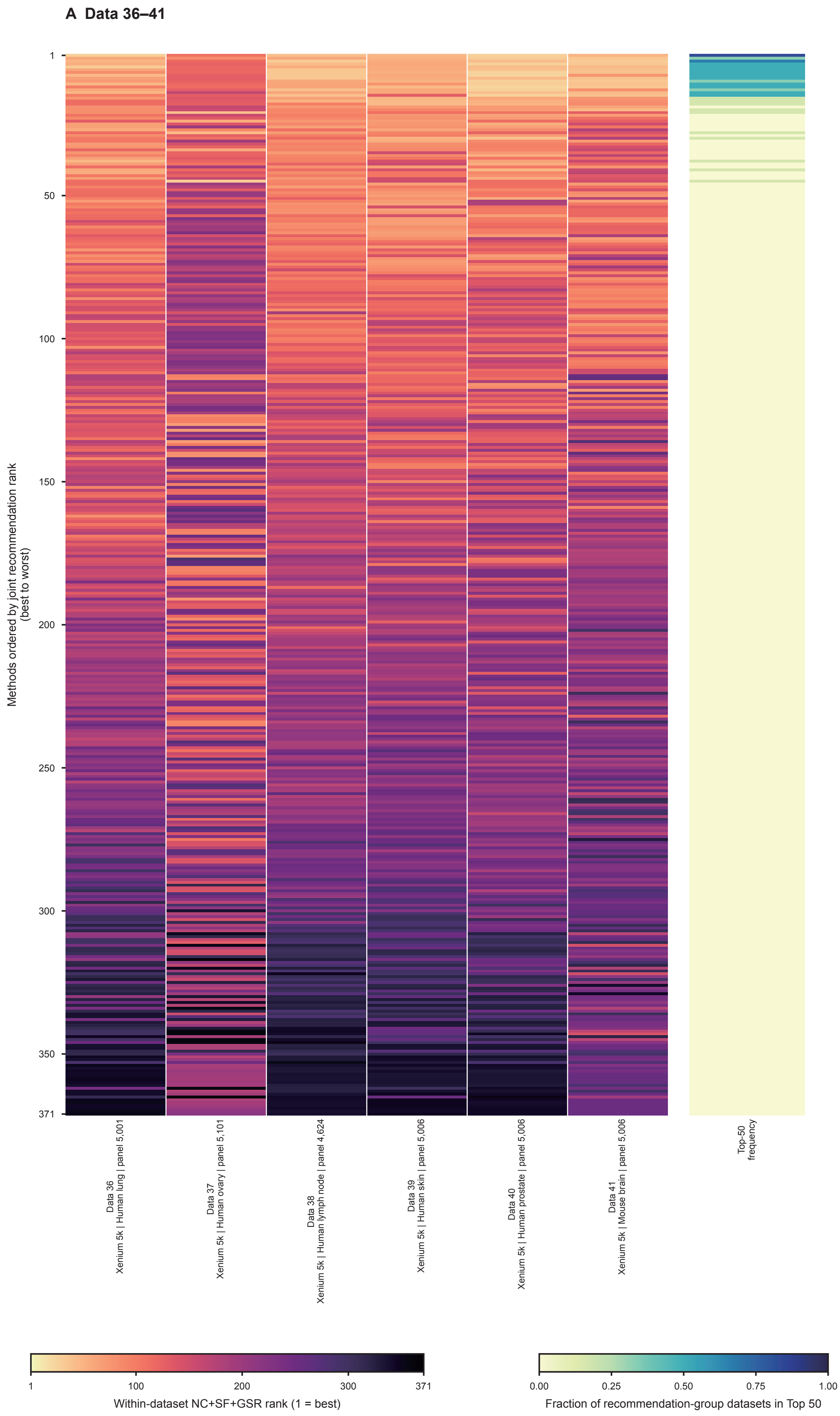


Rows follow the joint recommendation rank: aggregate of mean within-dataset component ranks across all displayed datasets.

#### Figure S21

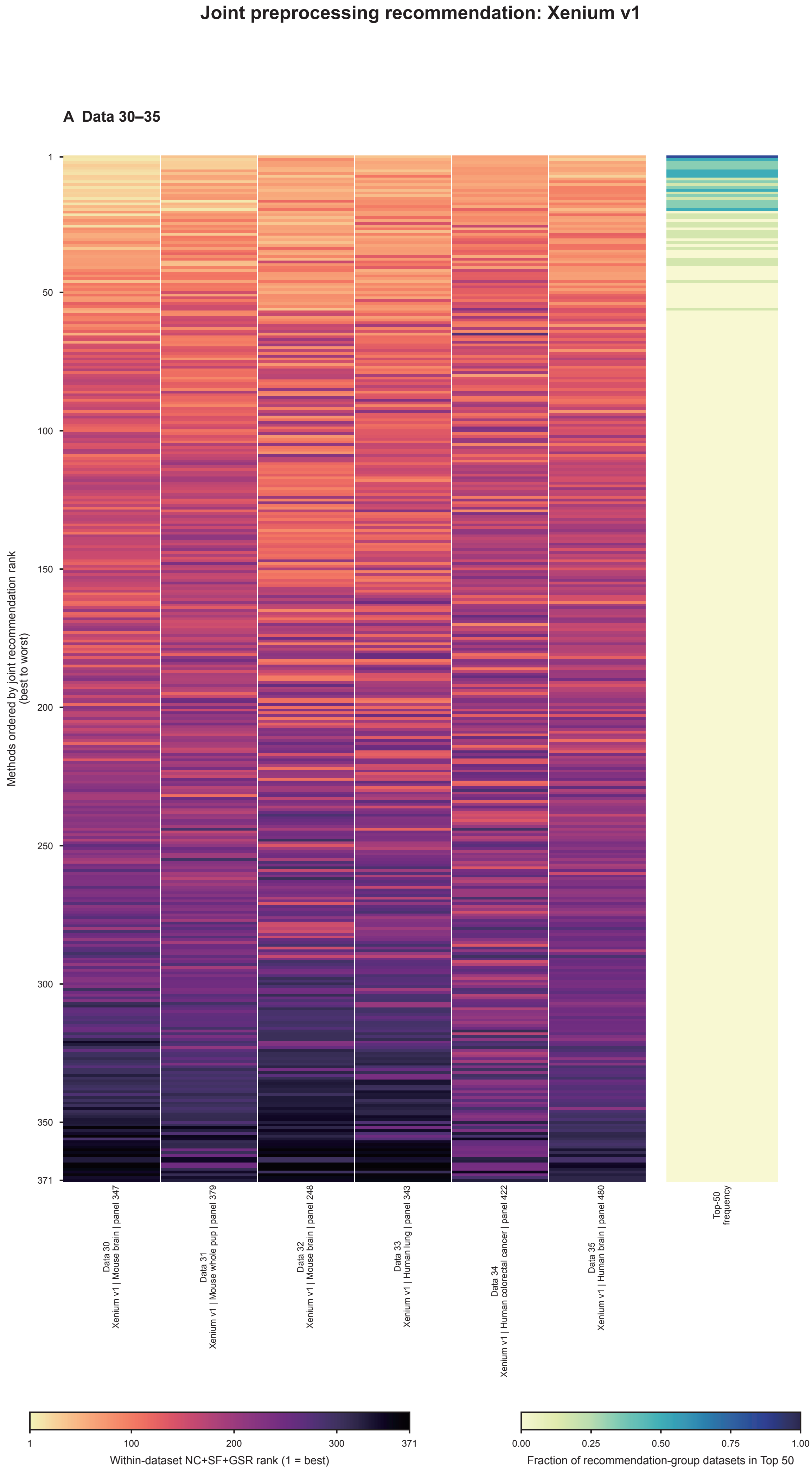


Rows follow the joint recommendation rank: aggregate of mean within-dataset component ranks across all displayed datasets.

### Figure S22

#### Clustered preprocessing-rank profiles across all 45 spatial datasets

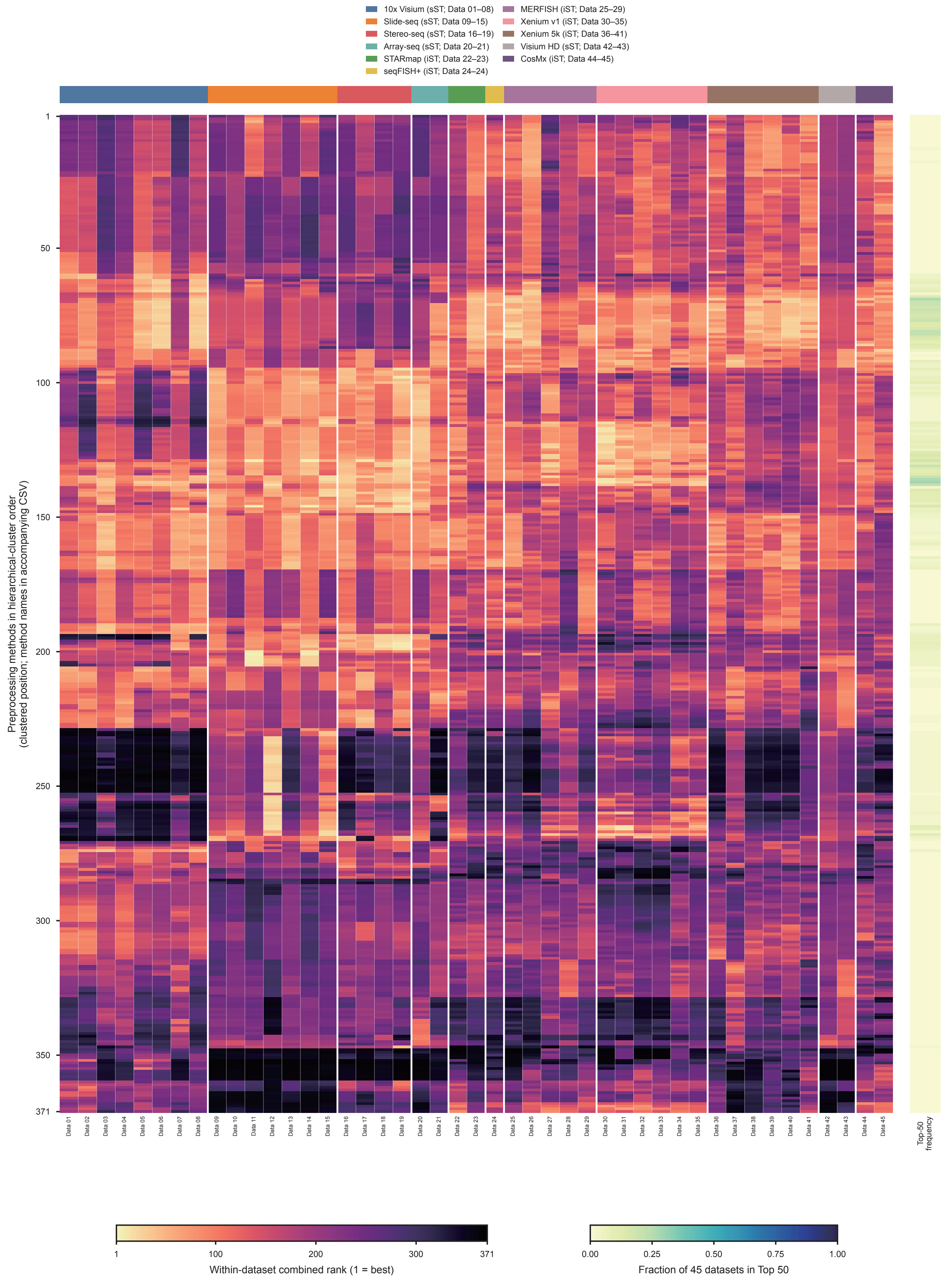


Rows: Ward hierarchical clustering of Euclidean distances between normalized 45-dataset rank profiles.
sST: NC+SF; iST: NC+SF+GSR.

Figure S23

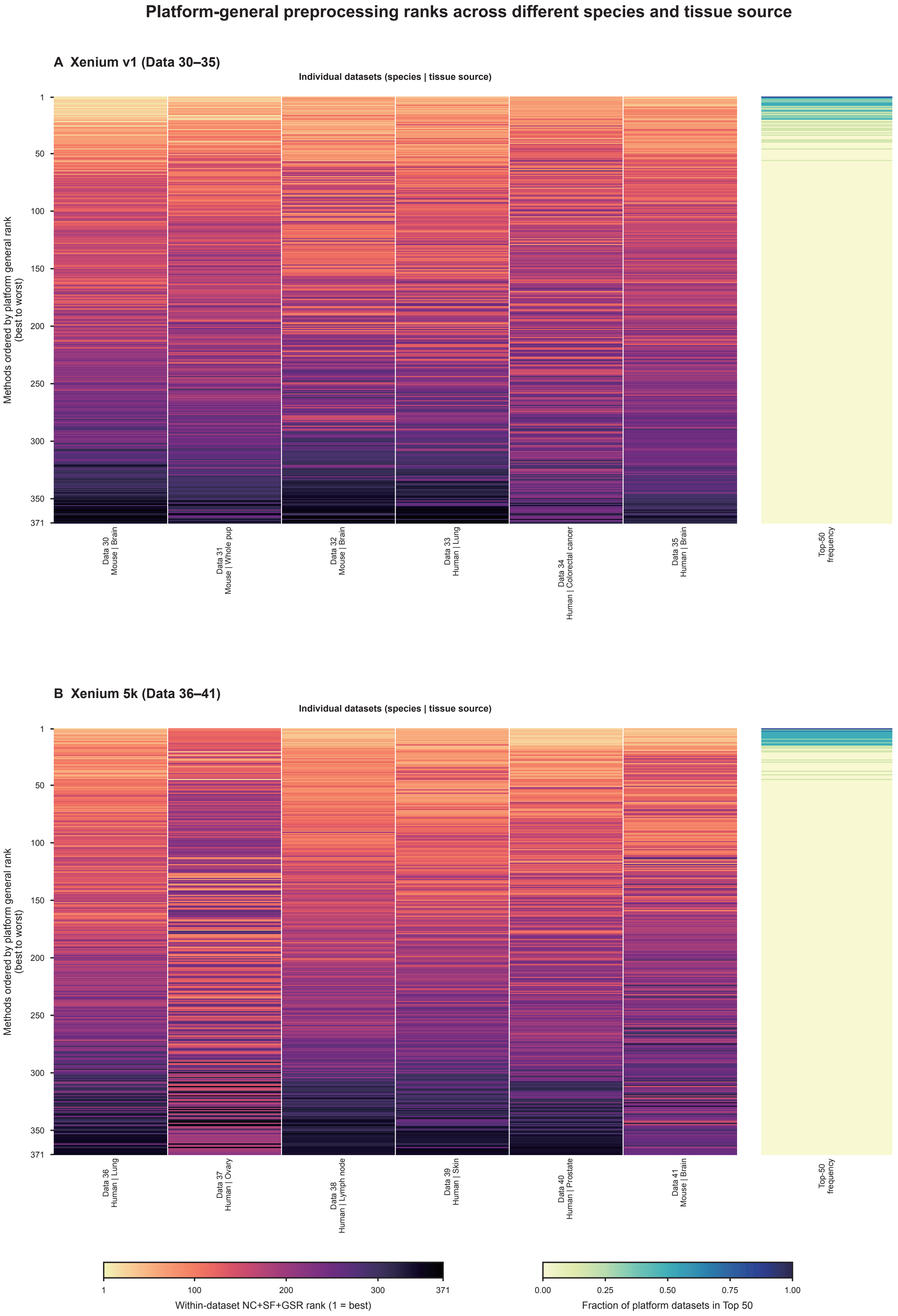


Rows follow each platform's general rank: aggregate of mean within-dataset component ranks across all displayed datasets.

### Figure S24

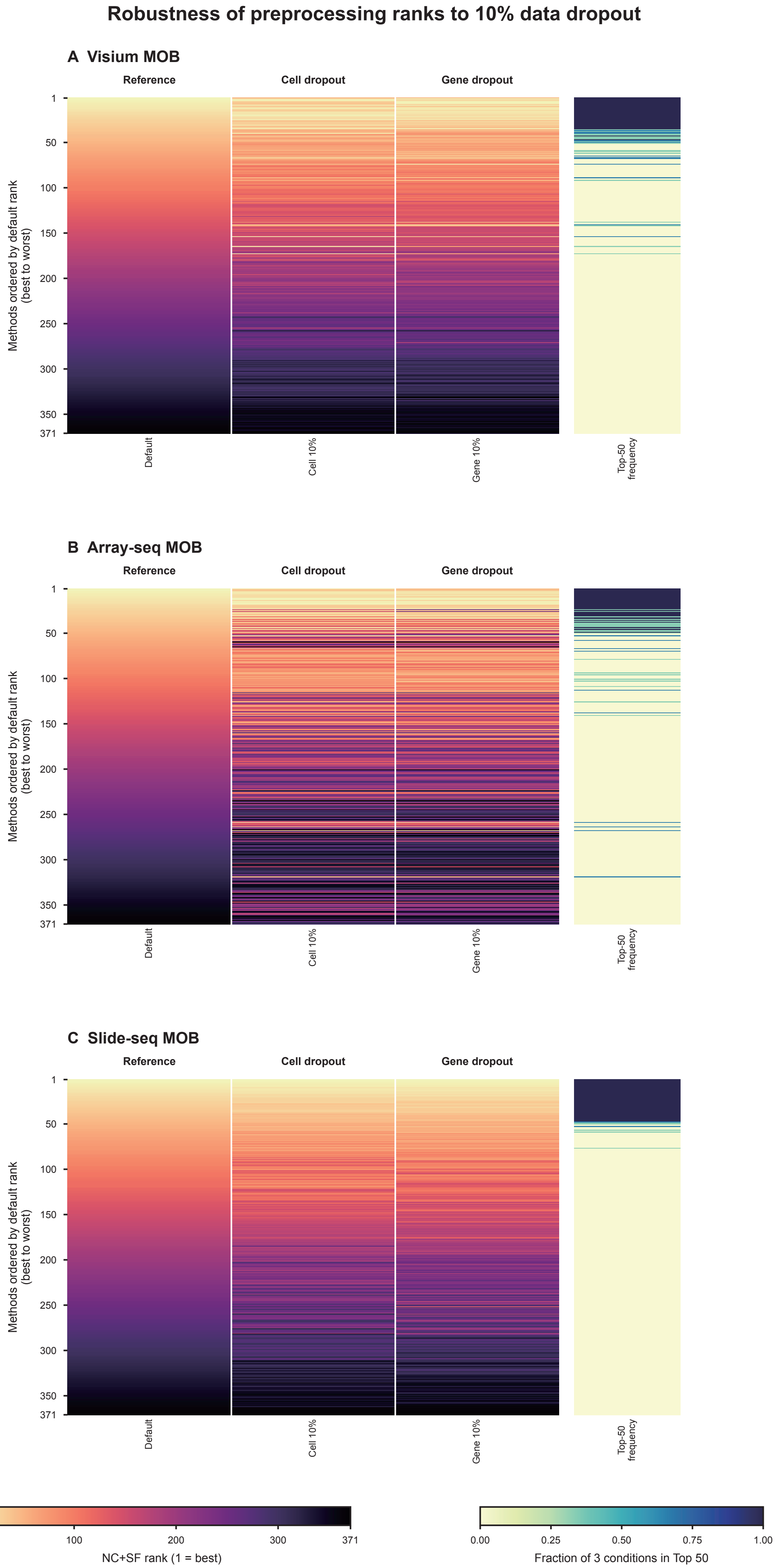


Default: k=50, PCs=20, Euclidean (L2). Perturbations remove 10% of cells or 10% of genes.

**Figure S25**

#### Robustness of imaging-data preprocessing ranks to 10% data dropout

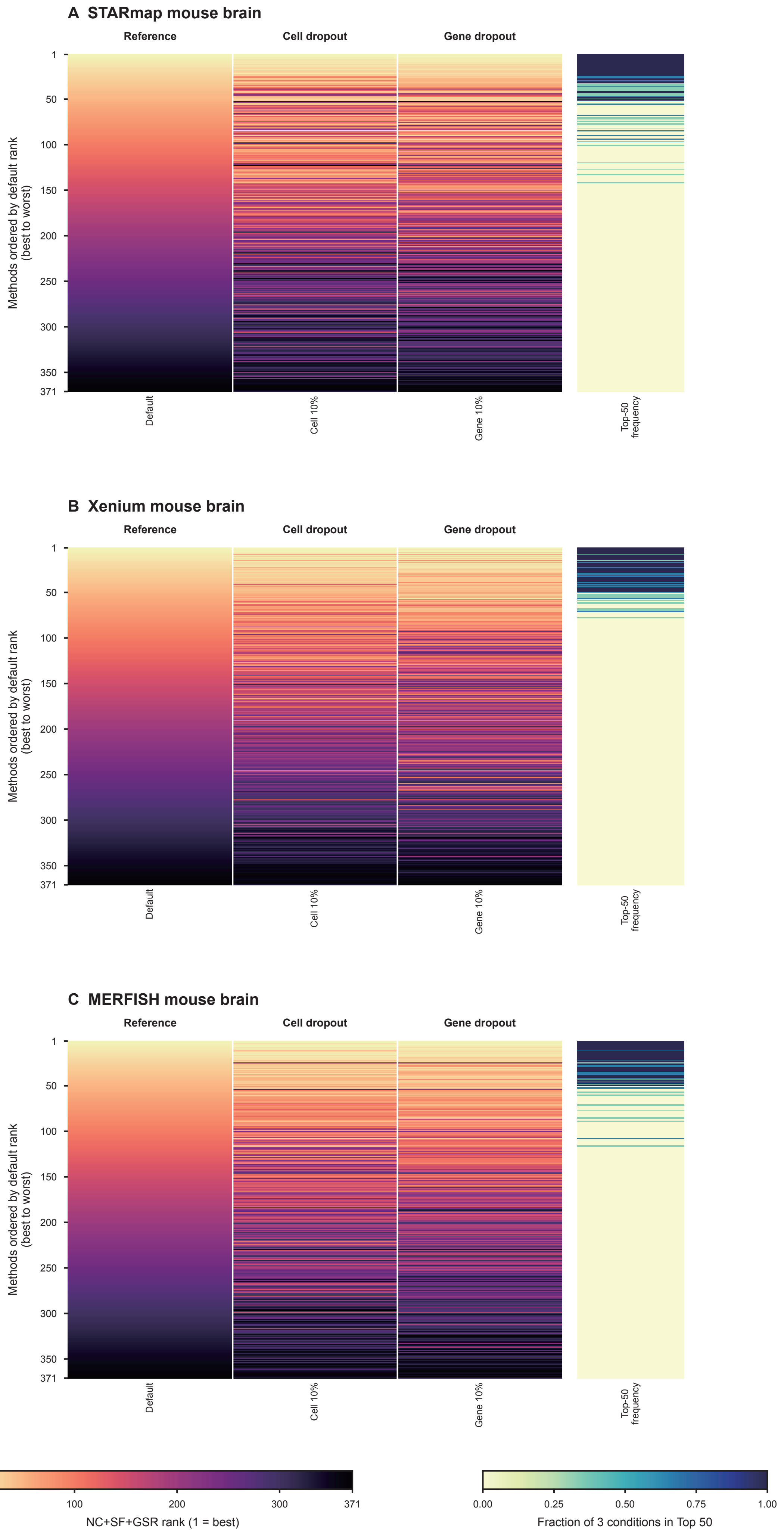


Default: k=50, PCs=20, Euclidean (L2). Perturbations remove 10% of cells or 10% of genes.

### Figure S26

#### Robustness of imaging-data preprocessing ranks across parameters settings

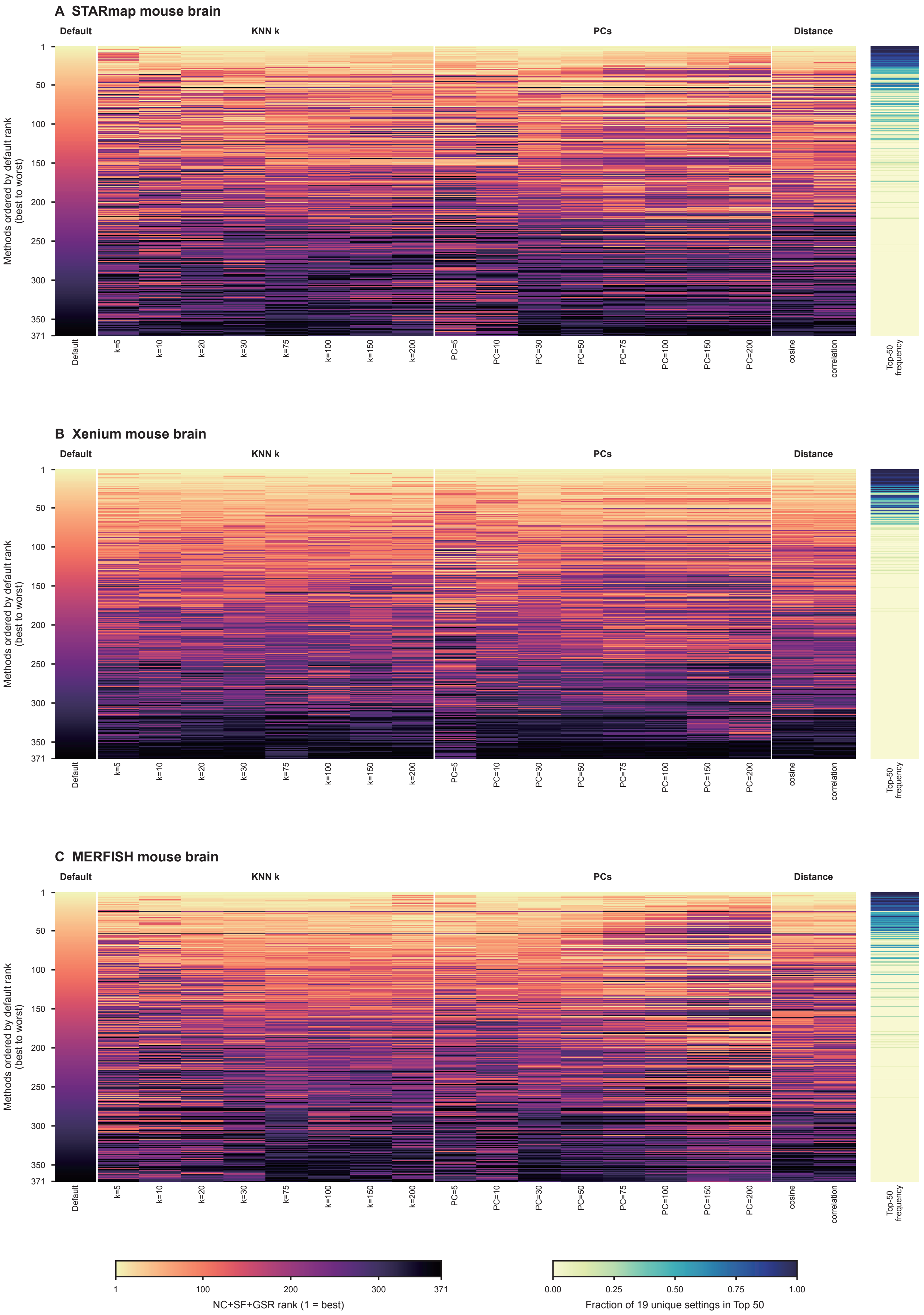


Default: k=50, PCs=20, Euclidean (L2). Top-50 frequency uses one default and 18 non-default perturbations.

Figure S27

#### Platform-general preprocessing ranks across sequencing-based spatial datasets

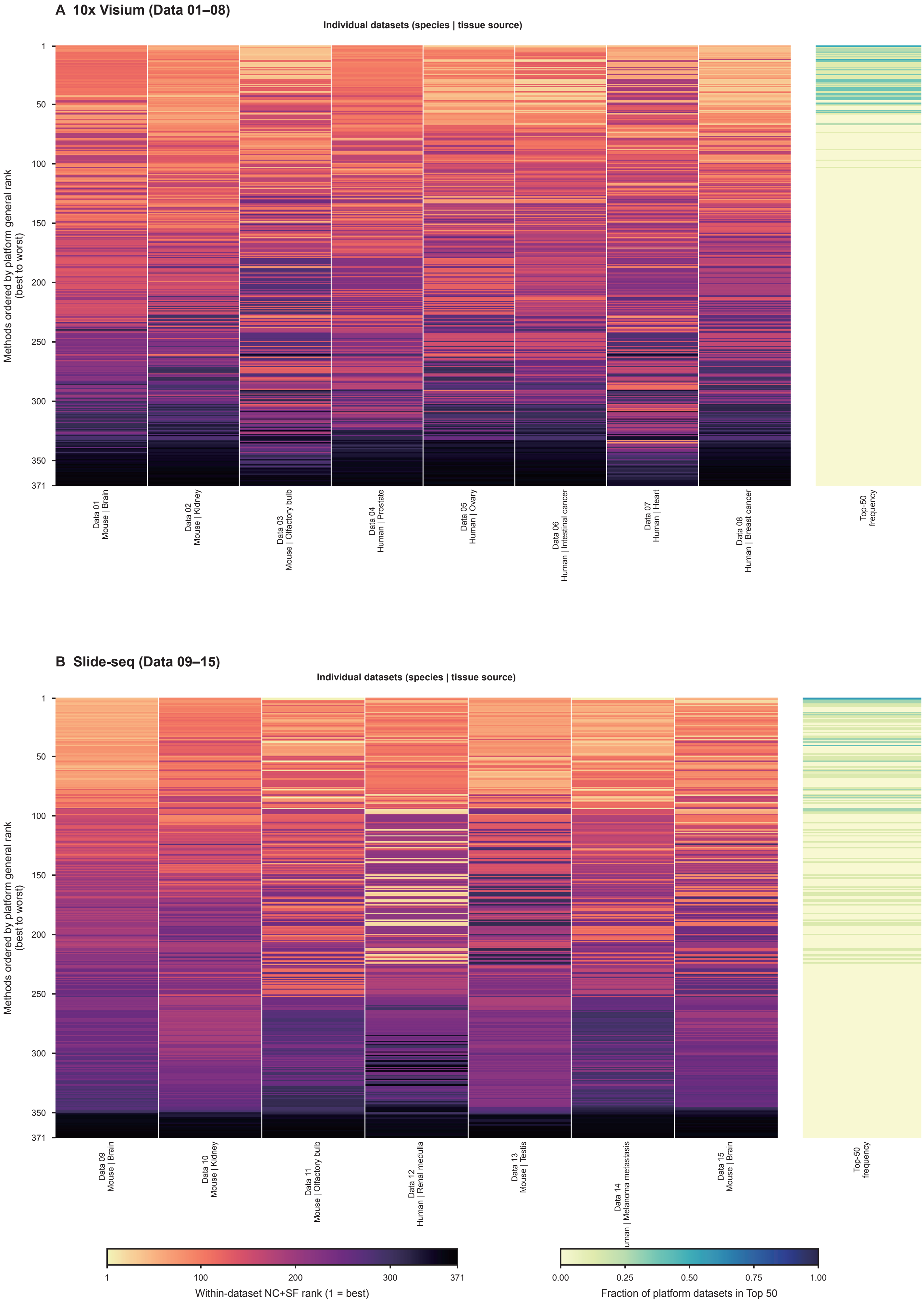


Rows follow each platform's general rank: aggregate of mean within-dataset component ranks across all displayed datasets.

#### Figure S28

### Cross-species stability of Stereo-seq preprocessing ranks

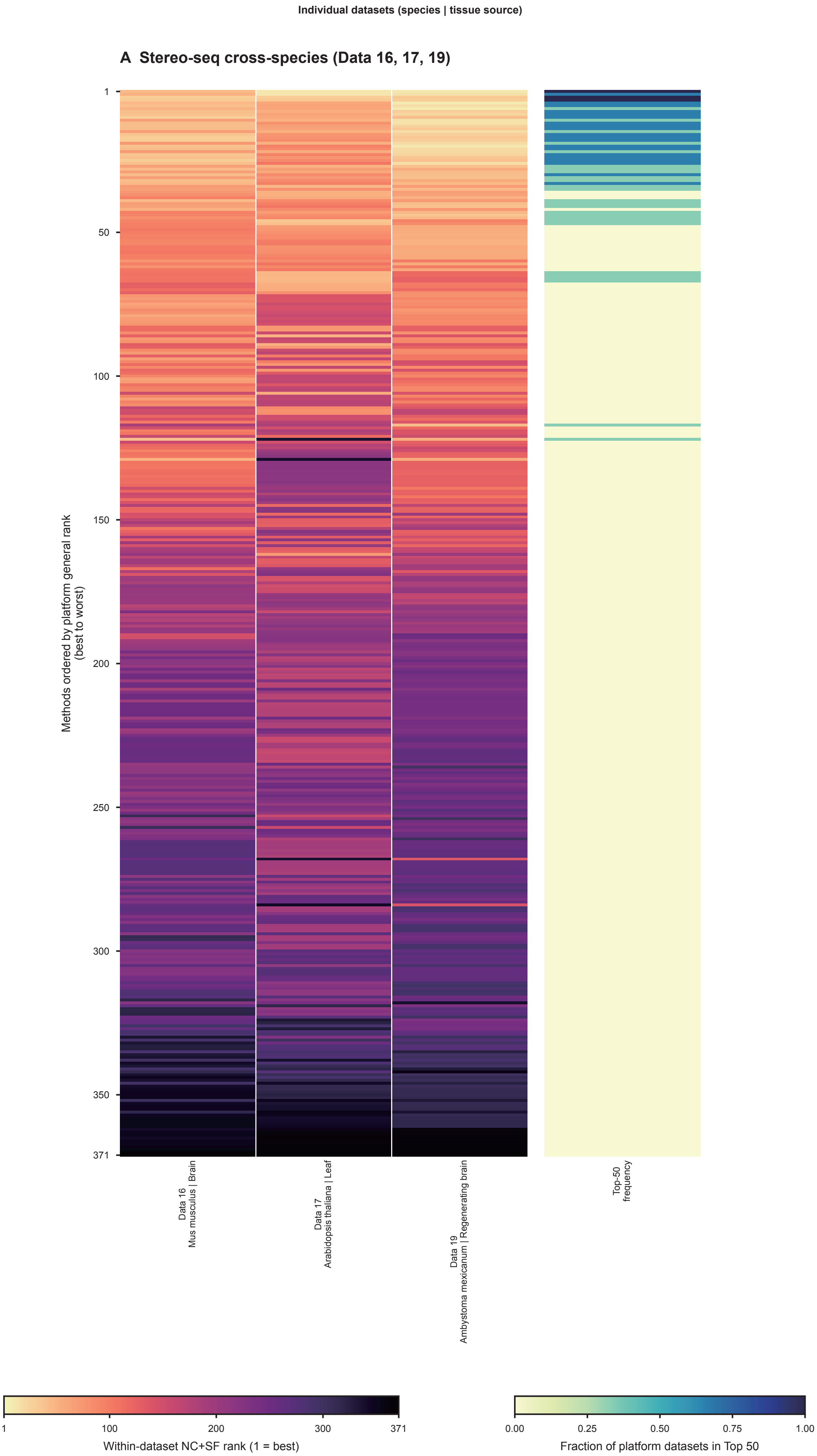


Rows follow each platform's general rank: aggregate of mean within-dataset component ranks across all displayed datasets.

### Figure S29

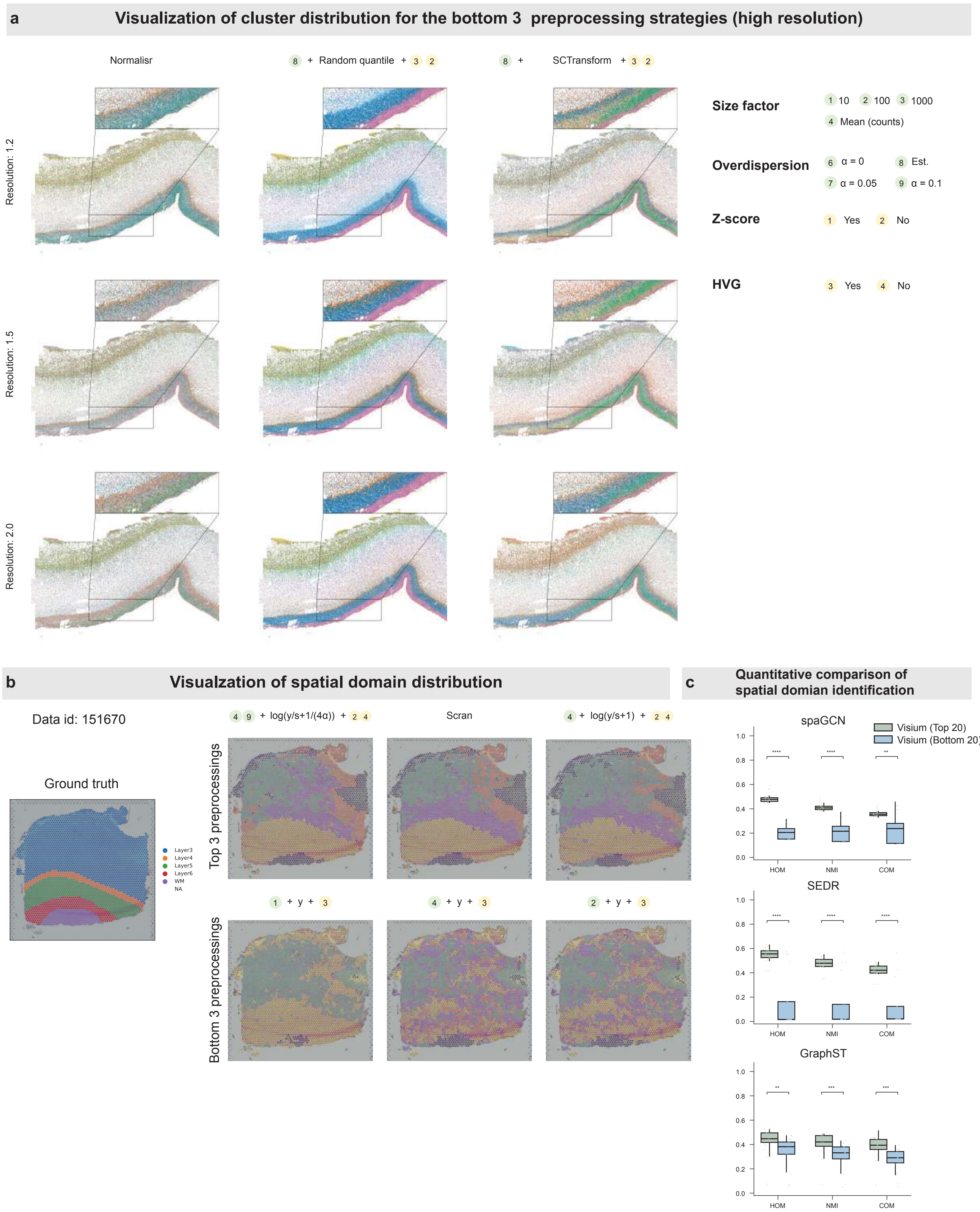
a
Visualization of cluster distribution for the bottom 3 preprocessing strategies (high resolution)
Normalisr
8 + Random quantile + 3 2
8 + SCTransform + 3 2
Resolution: 1.2
Resolution: 1.5
Resolution: 2.0
Size factor
1 10 2 100 3 1000
4 Mean (counts)
Overdispersion
6 α = 0
8 Est.
7 α = 0.05
9 α = 0.1
Z-score
1 Yes 2 No
HVG
3 Yes 4 No
b
Visualzation of spatial domain distribution
Data id: 151670
Ground truth
Layer3
Layer4
Layer5
Layer6
WM
NA
4 9 + log(y/s+1/(4α)) + 2 4
Scran
4 + log(y/s+1) + 2 4
Top 3 preprocessings
1 + y + 3
4 + y + 3
2 + y + 3
Bottom 3 preprocessings
c
Quantitative comparison of spatial domian identification
spaGCN
Visium (Top 20)
Visium (Bottom 20)
SEDR
GraphST
HOM
NMI
COM

### Figure S30

#### a Quantitative comparison of spatially variable gene detection

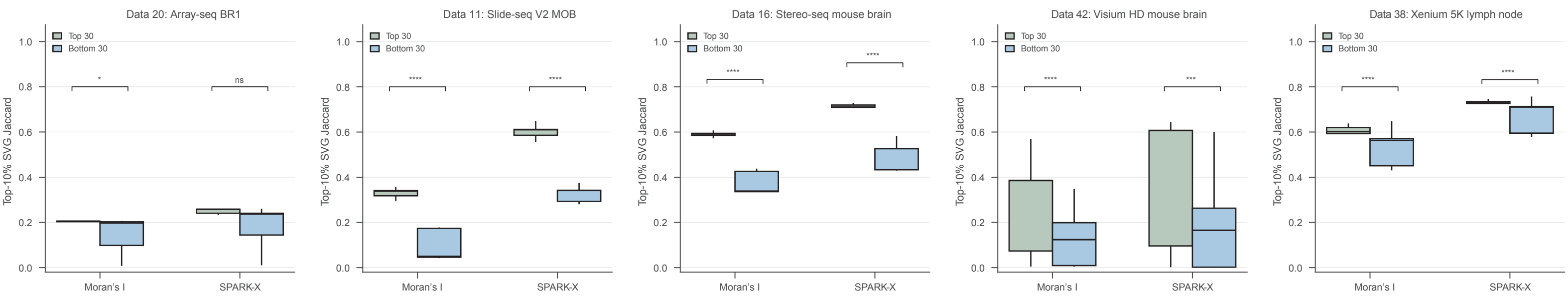


#### b Quantitative comparison of marker recovery

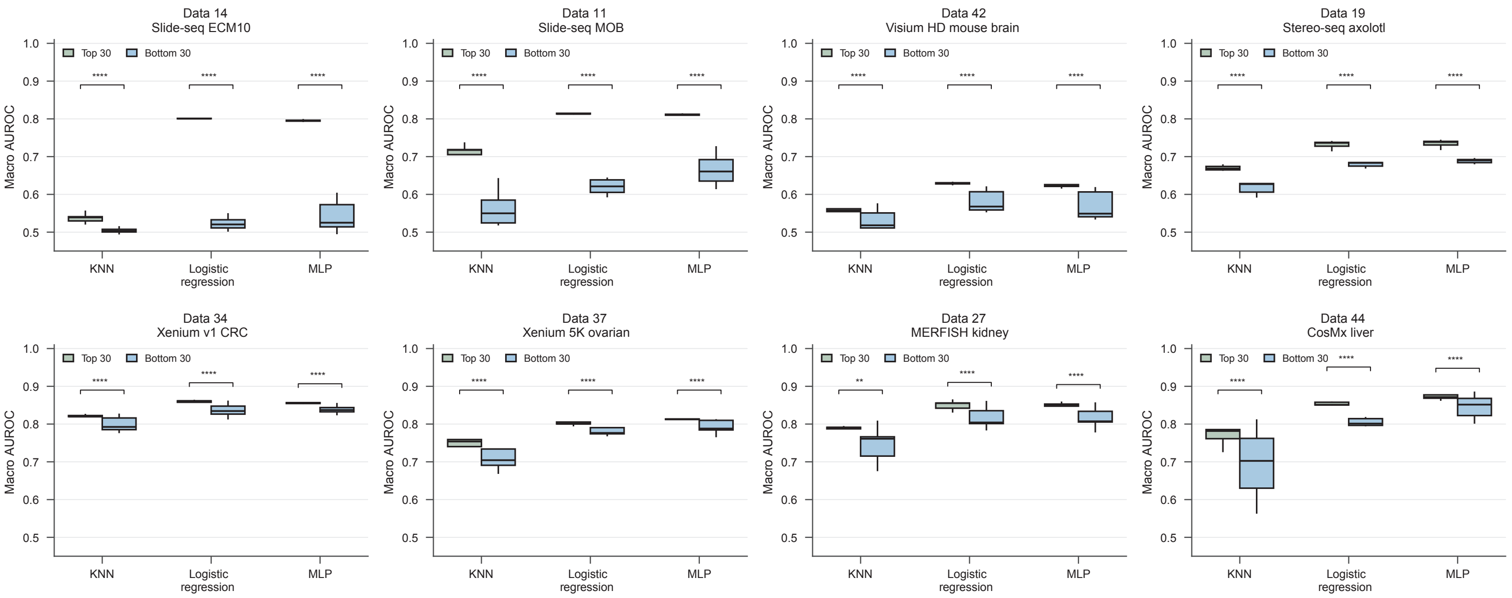

### Figure S31

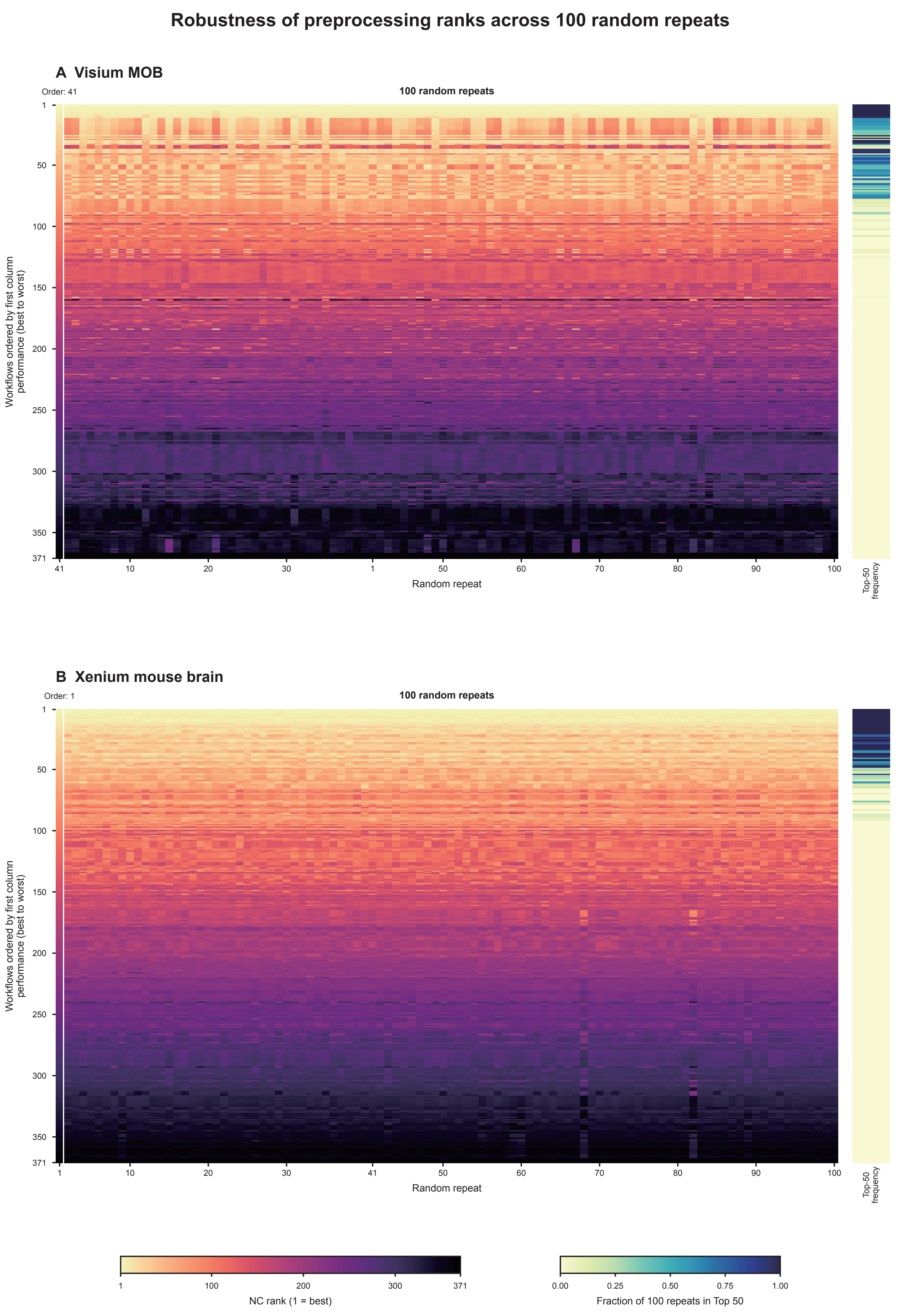


Each repeat is ranked independently by NC (lower is better); row order is fixed by the first displayed column.

### Figure S32

#### Robustness of preprocessing ranks across neighborhood settings

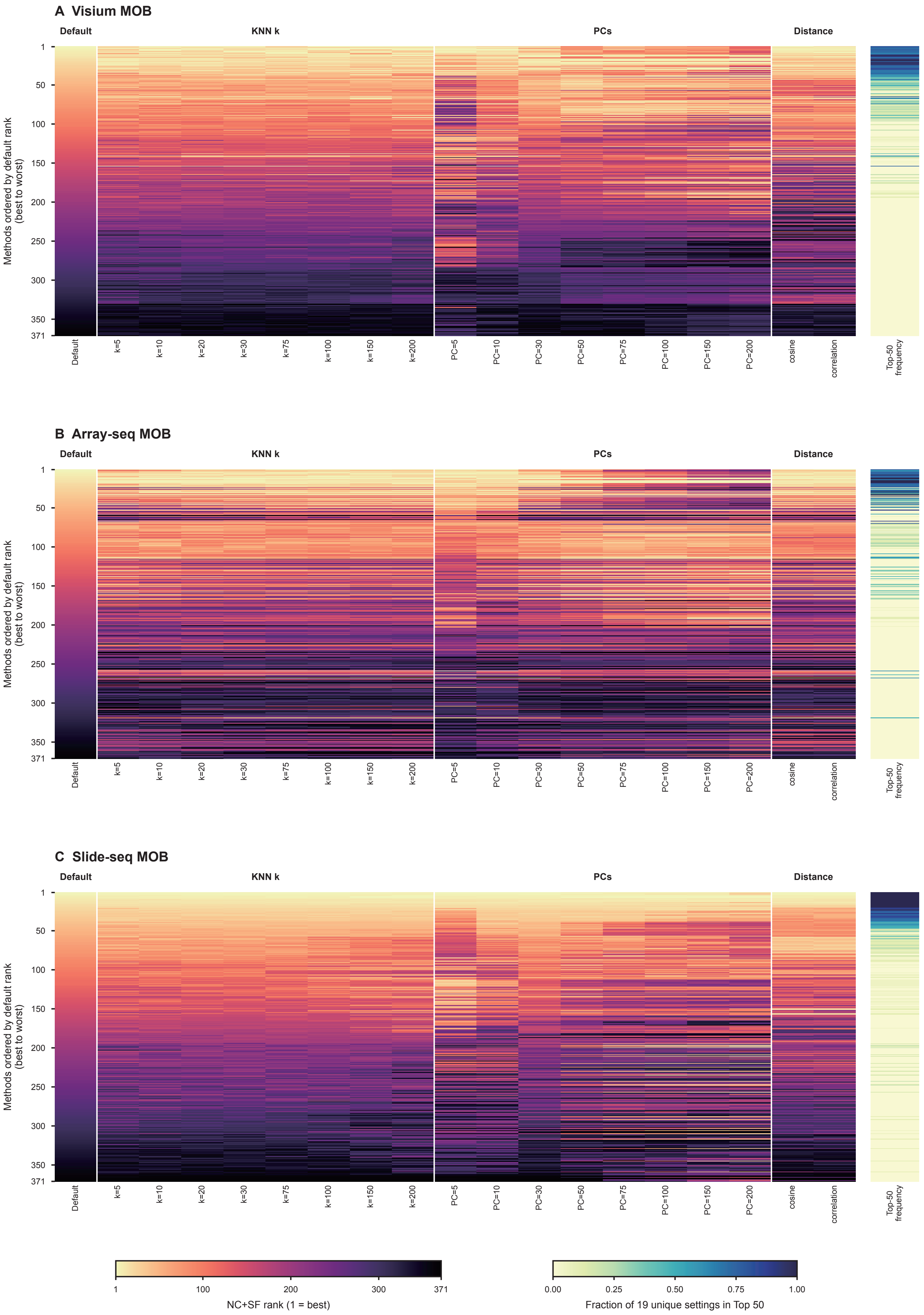


Default: k=50, PCs=20, Euclidean (L2). Top-50 frequency uses one default and 18 non-default perturbations.

#### Figure S33

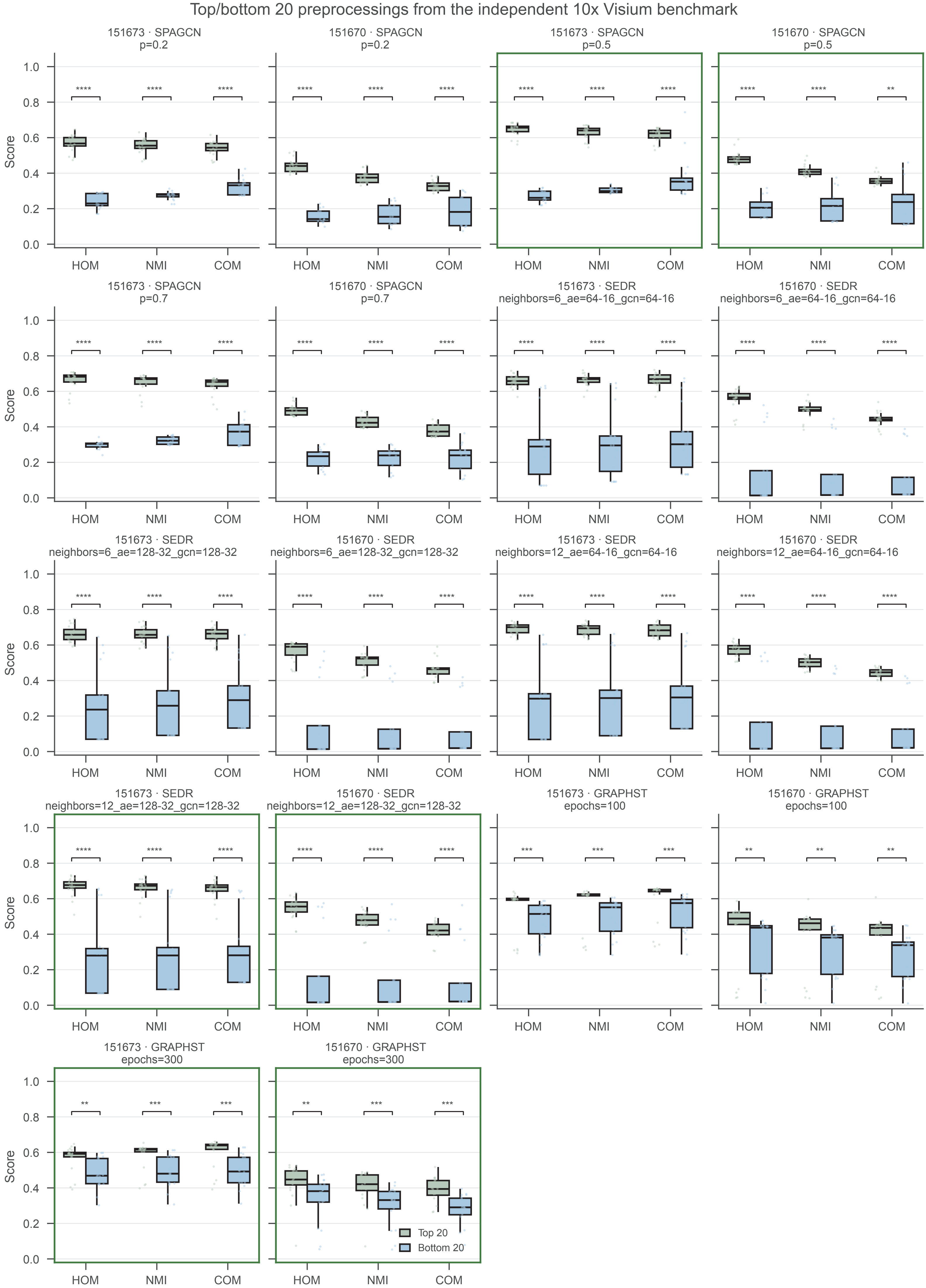
Top/bottom 20 preprocessings from the independent 10x Visium benchmark
151673 · SPAGCN p=0.2
151670 · SPAGCN p=0.2
151673 · SPAGCN p=0.5
151670 · SPAGCN p=0.5
151673 · SPAGCN p=0.7
151670 · SPAGCN p=0.7
151673 · SEDR neighbors=6_ae=64-16_gcn=64-16
151670 · SEDR neighbors=6_ae=64-16_gcn=64-16
151673 · SEDR neighbors=6_ae=128-32_gcn=128-32
151670 · SEDR neighbors=6_ae=128-32_gcn=128-32
151673 · SEDR neighbors=12_ae=64-16_gcn=64-16
151670 · SEDR neighbors=12_ae=64-16_gcn=64-16
151673 · SEDR neighbors=12_ae=128-32_gcn=128-32
151670 · SEDR neighbors=12_ae=128-32_gcn=128-32
151673 · GRAPHST epochs=100
151670 · GRAPHST epochs=100
151673 · GRAPHST epochs=300
151670 · GRAPHST epochs=300
Score
HOM
NMI
COM
Top 20
Bottom 20
